# Design of Cell-Permeable Inhibitors of Eukaryotic Translation Initiation Factor 4E (eIF4E) for Inhibiting Aberrant Cap-Dependent Translation in Cancer

**DOI:** 10.1101/2023.05.23.541912

**Authors:** Emilio L. Cárdenas, Rachel L. O’Rourke, Arya Menon, Jennifer Meagher, Jeanne Stuckey, Amanda L. Garner

**Affiliations:** Department of Medicinal Chemistry, College of Pharmacy, University of Michigan, Ann Arbor, Michigan 48109, United States; Life Science Institute, University of Michigan, Ann Arbor, Michigan 48109, United States; Rogel Cancer Center, University of Michigan, Ann Arbor, Michigan 48109, United States

**Author notes:** Corresponding Author: Amanda L. Garner - Department of Medicinal Chemistry, College of Pharmacy, University of Michigan, 1600 Huron Parkway, NCRC B520, Ann Arbor, Michigan 48109, USA. ORCID: orcid.org/0000-0002-0870-3347.

## Abstract

Eukaryotic translation initiation factor 4E (eIF4E) is an RNA-binding protein that binds to the m^7^GpppX-cap at the 5′ terminus of coding mRNAs to initiate cap-dependent translation. While all cells require cap-dependent translation, cancer cells become addicted to enhanced translational capacity, driving the production of oncogenic proteins involved in proliferation, evasion of apoptosis, metastasis, and angiogenesis among other cancerous phenotypes. eIF4E is the rate-limiting translation factor and its activation has been shown to drive cancer initiation, progression, metastasis, and drug resistance. These findings have established eIF4E as a translational oncogene and promising, albeit challenging, anti-cancer therapeutic target. Although significant effort has been put forth towards inhibiting eIF4E, the design of cell-permeable, cap-competitive inhibitors remains a challenge. Herein, we describe our work towards solving this long-standing challenge. By employing an acyclic nucleoside phosphonate prodrug strategy, we report the synthesis of cell-permeable inhibitors of eIF4E binding to capped mRNA to inhibit cap-dependent translation.

## INTRODUCTION

Protein synthesis is the penultimate step of gene expression, dynamically controlling the makeup of the proteome in response to cellular stimuli. While all cells require protein synthesis, cancer cells, in particular, become addicted to enhanced translational capacity, driving the production of proteins necessary for proliferation and survival, in addition to other hallmarks of cancer.^1–3^ Indeed, in cancer, changes in mRNA translation have been shown to be more extensive than changes in transcription, enabling the cell to use existing mRNA pools to quickly regulate cellular state and activity.^3, 4^ Although many mechanisms lead to dysregulation of translational control in cancer, the predominant paths are through hyperactivation of the PI3K-AKT-mTORC1 and RAS-RAF-MAPK, and MYC signaling pathways,^5, 6^ all of which converge on the regulation of cap-dependent mRNA translation, the major translational program leading to the translation of mRNAs encoding oncoproteins and growth and survival factors including c-Myc, cyclin Ds, and ODC among others (Figure 1). ^7–11^

**Figure 1.**
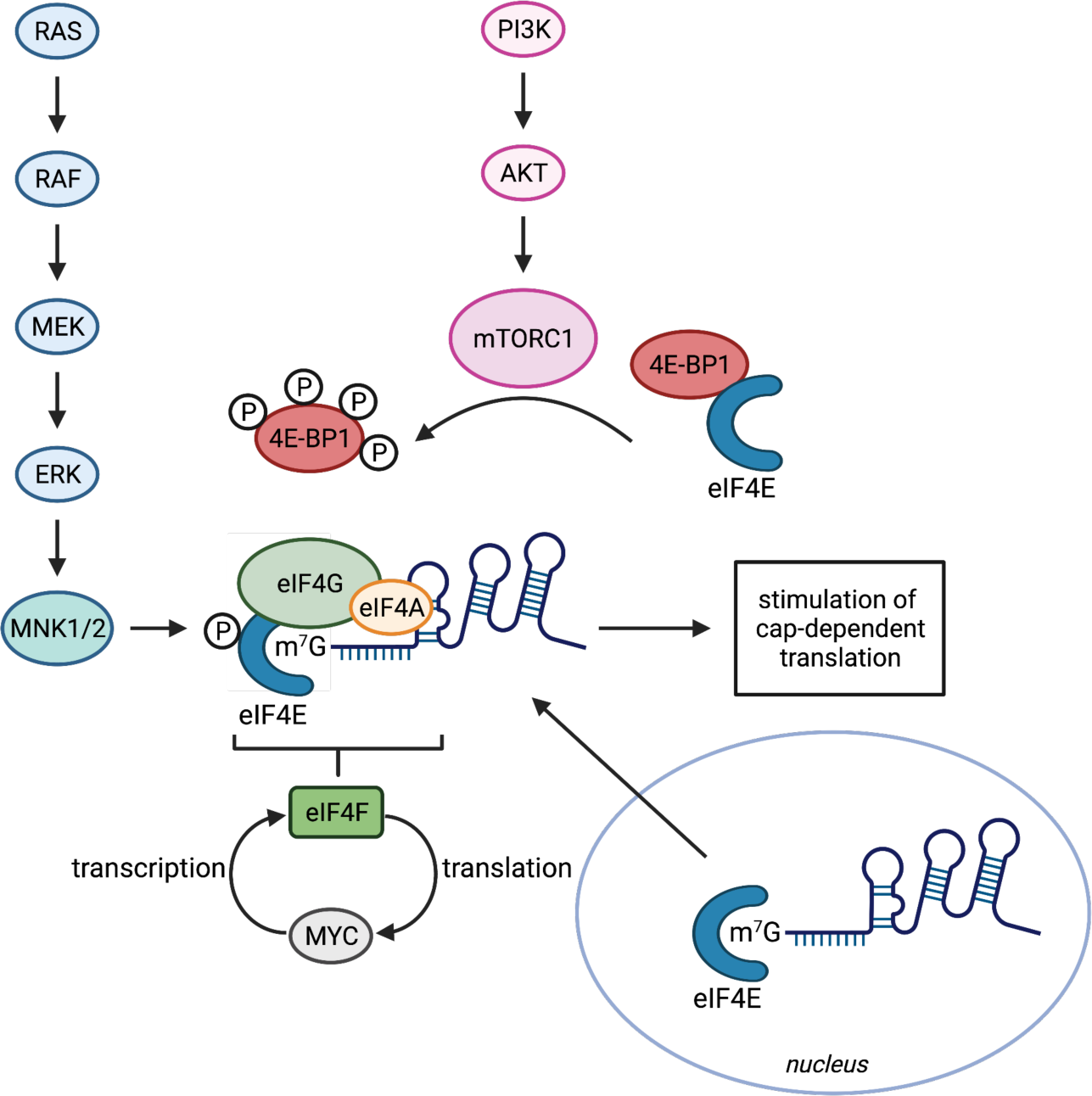
Activation of eIF4E and cap-dependent translation.

The initiation of cap-dependent mRNA translation is governed by the availability of eukaryotic translation initiation factor 4E (eIF4E). eIF4E is an RNA-binding protein that plays two important roles in the initiation of cap-dependent translation: (1) eIF4E binds to the m^7^GpppX-cap at the 5′ terminus of coding mRNAs to promote the assembly of the eIF4F translation initiation complex, and (2) it promotes the nuclear export of select cap-dependent transcripts (Figure 1).^8, 10–12^ Thus, eIF4E plays the critical role of both selecting and executing the translation of specific transcripts. Based on the significance of these functions in maintaining cellular homeostasis, eIF4E is the rate-limiting translation factor and its overexpression has been shown to induce tumorigenesis.^10^ Consequently, elevated levels are found in many human cancers, and in many cases, elevated eIF4E has been linked to advanced disease and reduced survival.^1–3, 8, 9, 13^

While cancer cells have been shown to rely on eIF4E overexpression to promote oncogenesis, importantly, a 50% reduction in eIF4E was found to impede cellular transformation and significantly decrease tumor growth in a KRAS-driven lung cancer model.^14^ Most notably, using this Eif4e haploinsufficient mouse model, it was revealed that knockdown of eIF4E was compatible with normal development and global protein synthesis.^14^ Further evidence for the tolerability of eIF4E reduction in normal tissues stems from studies using eIF4E-targeted antisense oligonucleotides (ASO), which were similarly found to be non-toxic.^15^ Combined, these findings indicate that eIF4E-directed therapies are likely to exhibit suitable therapeutic index despite the widespread expression and activity of this translation initiation factor, and make eIF4E a promising, albeit challenging target in cancer drug discovery.

Because eIF4E’s primary function is binding to the m^7^GpppX cap of mRNAs, significant effort has been invested in designing cap analogues for inhibiting this activity.^16–24^ Yet, due to reliance on nucleoside monophosphate- or diphosphate-based scaffolds for many of the reported compounds, adequate cell permeability has remained an impedance towards use as eIF4E-targeted chemical probes for target validation studies (Figure 2).^22, 24^ To overcome this challenge, which has long plagued the field, herein we describe the design and synthesis of a series of acyclic nucleoside phosphonate prodrugs resulting in the first cell-permeable, cap-competitive inhibitors of eIF4E based upon a nucleotide scaffold.

**Figure 2.**
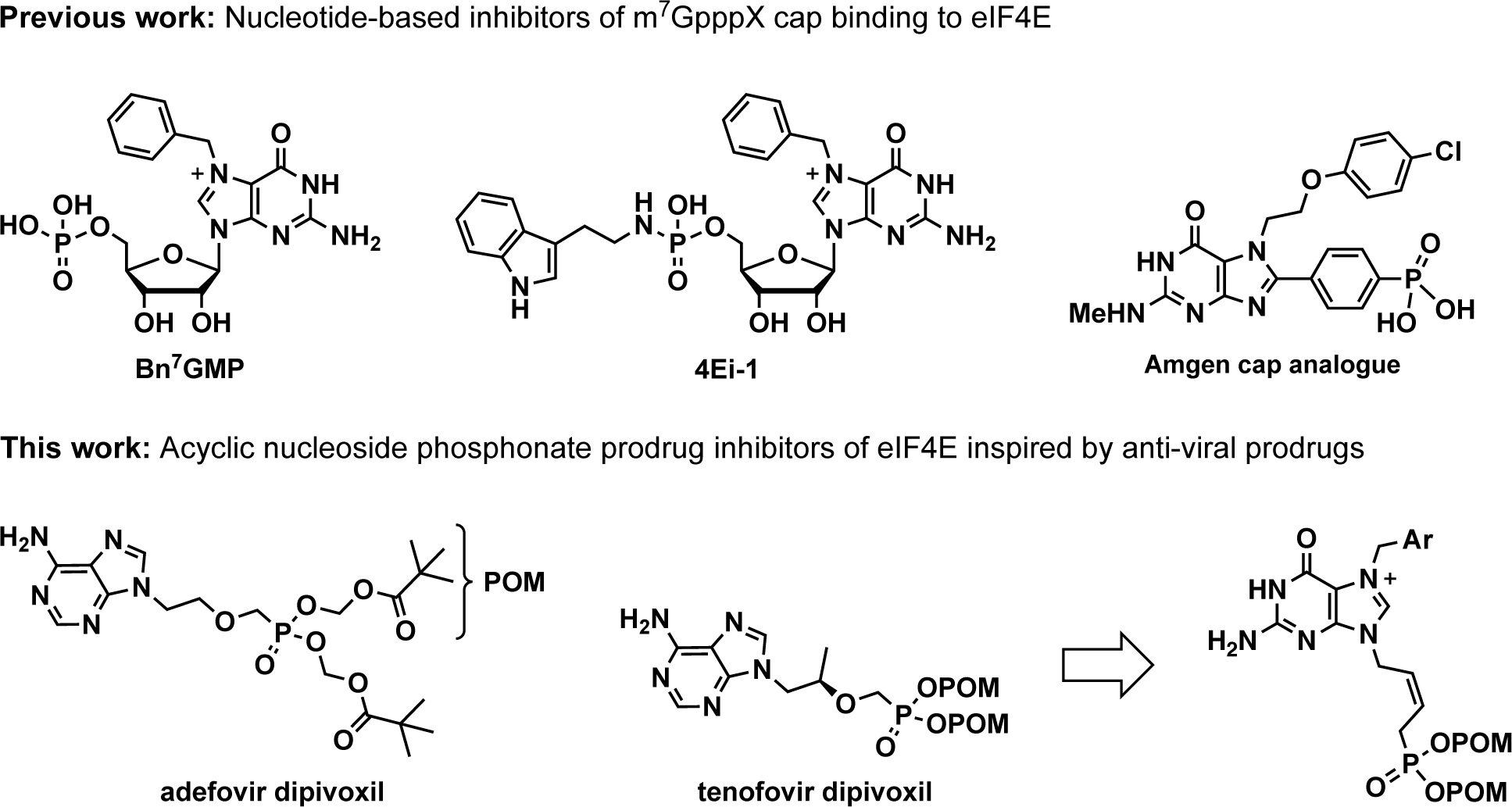
Examples of reported inhibitors of m^7^GpppX cap binding to eIF4E and proposed acyclic nucleoside phosphonate prodrugs inspired by the anti-viral drugs, adefovir and tenofovir dipivoxil.

## RESULTS AND DISCUSSION

### Design and synthesis of acyclic nucleoside phosphonate prodrug inhibitors of eIF4E

Based on crystal structures of eIF4E bound to several cap analogues,^25, 26^ we posited that the ribose ring, which has traditionally been included in the design of cap-based inhibitors like **4Ei-1** (Figure 2), serves merely as a linker between the nucleobase and phosphates, the key binding motifs within the cap structure, as it makes no contact with the protein. Indeed, Amgen’s cap inhibitor (Figure 2) explored a similar design strategy leading to improved inhibitory activity; however, this compound, which contains a negatively charged phosphonic acid, was found to lack cell penetrability.^26^

Inspired by nucleoside phosphonate prodrugs, upon which several important classes of anti-viral drugs have been centered (e.g., adefovir and tenofovir dipivoxil),^27–29^ we set out to design an analogous series of acyclic nucleoside phosphonate prodrugs targeting the eIF4E cap-binding pocket. As proof-of-concept for our prodrug approach, we took advantage of the bis-pivaloyloxymethyl (POM) ester protecting groups of adefovir/tenofovir (Figure 2) to mask the negative charge of the phosphonate motif for cell penetration, which through subsequent esterase-mediated deprotection in cells, would yield the bioactive compound.^27–29^ Using this design principle, we synthesized a series of prodrugs employing a common alkenyl phosphonate modification with varying N7 substitutions to initiate structure-activity relationship (SAR) studies (Figure 2 and Scheme 1). The linker strategy used was based on previous work by Agrofoglio and co-workers in their design of acyclic nucleoside phosphonate prodrugs containing a bis-POM (*E*)-4-phosphono-but-2-enyl linker as broad-spectrum antiviral agents.^30^ However, due to the concave shape of the cap-binding pocket of eIF4E, we envisioned that a (*Z*)-4-phosphono-but-2-enyl linker would appropriately position the phosphonate toward the positively-charged cleft that is occupied by the phosphate of Bn^7^GMP (Figure 3).^25^ Thus, we synthesized a library of *N*7-substituted cap analogues containing a bis-POM (*Z*)-4-phosphono-but-2-enyl group in place of the phosphorylated ribose motif of the canonical cap structure.

**Figure 3.**
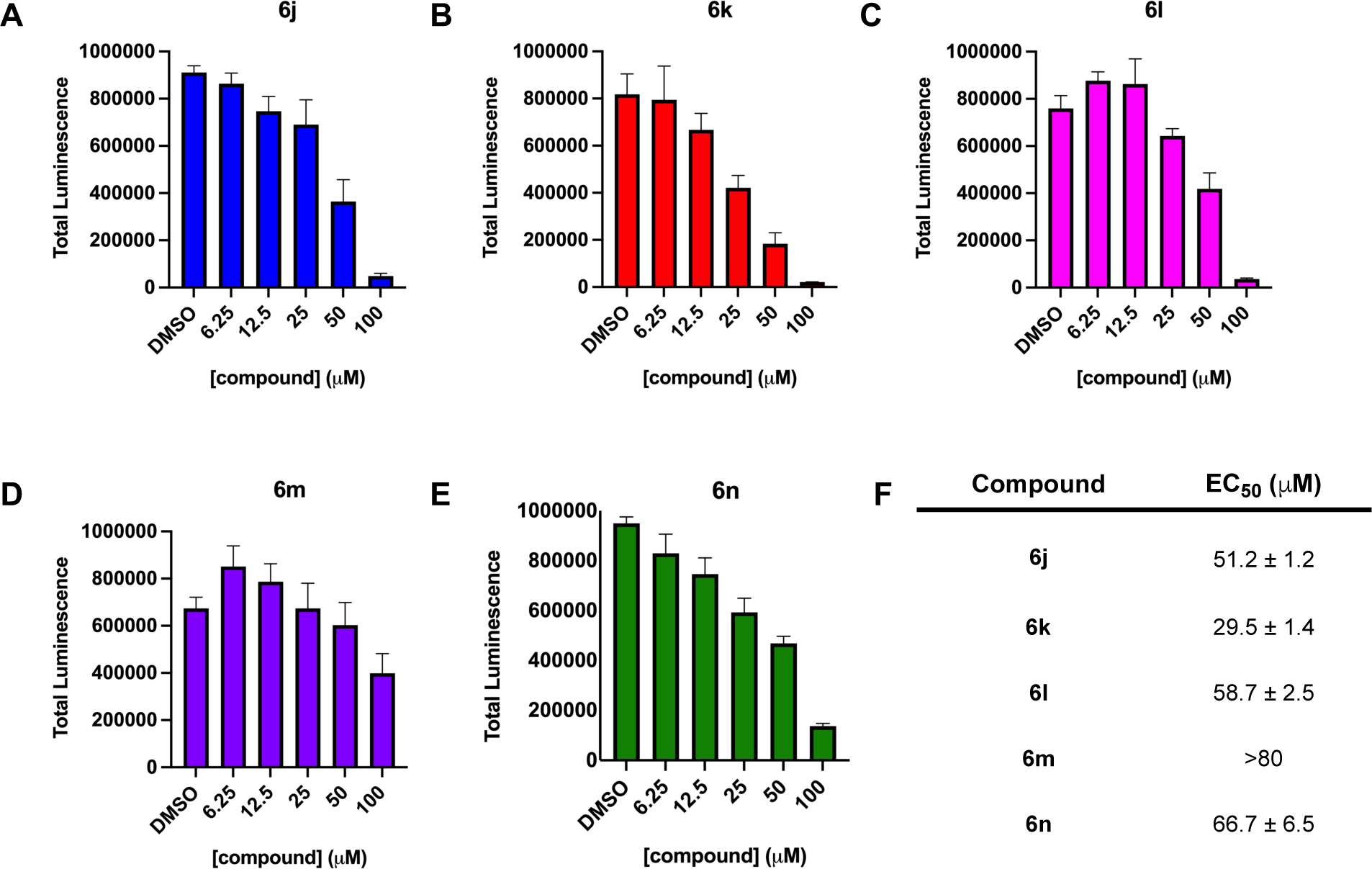
Anti-proliferative activity of bis-POM cap analogue prodrugs (A) **6j**, (B) **6k**, (C) **6l**, (D) **6m**, and (E) **6n**. (F) Table of EC_50_ values.

**Scheme 1.**
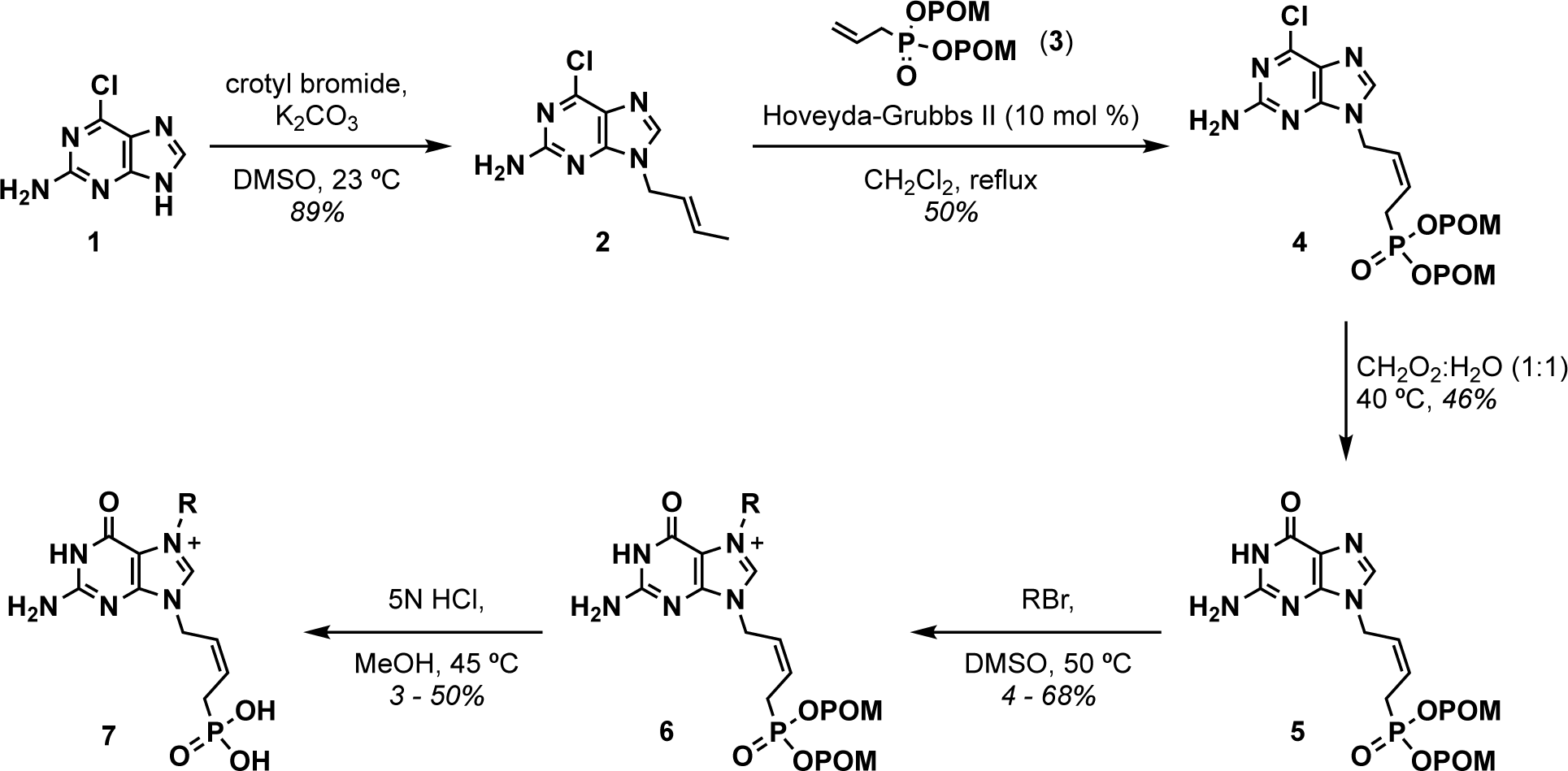
Synthetic scheme for the preparation of cap analogue prodrugs **6** and phosphonic acids 7.

Our synthetic route is detailed in Scheme 1 and began with the crotylation of 2-amino-6-chloropurine (**1**) to afford the *N*-9-crotyl purine (**2**) as a single regioisomer. Cross-metathesis of **2** with bis-POM allylphosphonate **3** catalyzed by Hoveyda-Grubbs 2^nd^ generation catalyst resulted in a mixture of *Z*/*E* isomers, which following purification by column chromatography, provided exclusively the *Z*-adduct **4** in moderate yield. The resulting 6-chloropurine analogue was then hydrolyzed in an aqueous solution of formic acid to provide guanine **5** in 46% yield. Alkylation of **5** with various alkyl bromides resulted in the desired bis-POM-substituted cap analogues **6a**−**6n** in 4−68% yield. Unprotected phosphonic acids, **7a**−**7n**, were also prepared for biochemical testing via hydrolysis with an aqueous solution of 5N HCl in MeOH.

### Biochemical inhibitory activity and target engagement

As preliminary characterization, unprotected phosphonic acids (**7**) were tested for competitive inhibition of eIF4E using a fluorescence polarization (FP) assay employing fluorescein-labeled m^7^GTP.^31^ Promising inhibitory activity was observed with a select group of compounds with measured IC_50_ values of 1−16 μM, in-line with those of m^7^GMP/m^7^GDP (Table 1 and Figure S1). The 7-methyl substituted analogue **7a** was found to be 2.5-fold less active than m^7^GMP with a measured IC_50_ value of 16 µM. In-line with previous observations by Rentmeister and co-workers in characterizing cap analogues for promoting cap-dependent mRNA translation,^32^ **7b** and **7c** were found be inactive. Contrary to previous studies,^26, 33^ **7d** and **7f** were also found to be inactive. Interestingly, the 4-chlorobenzyl-substituted derivative (**7e**) exhibited moderate activity (IC_50_ value of 58 µM) in comparison to the inactive benzyl analogue (**7d**), reiterating the significance of halogen-bonding interactions within the hydrophobic pocket distal to the guanine core.^34^ Further extension into this pocket by the biphenyl derivative **7g** markedly improved activity and this analogue exhibited a measured IC_50_ value of 10 µM. While cinnamyl-substituted analogue **7h** and 3-phenylpropyl derivative **7i** lacked activity, 4-chloro-phenoxyethyl-substituted^26^ **7j** yielded the most potent cap analogue with a measured IC_50_ value of 1.1 µM. A small library of *N*-7-methylarene-substituted compounds (**7k−7n**) were also prepared and tested. All were found to be active with the methyl(imidazopyridine)-(**7m**), methyl(indole)-(**7n**), and methyl(benzofuran)-(**7k**) most active with IC_50_ values of 4.0, 5.6, and 7.7 µM, respectively, followed by methyl(benzothiophene) analogue **7l** with an IC_50_ value of 16 µM.

**Table 1.**
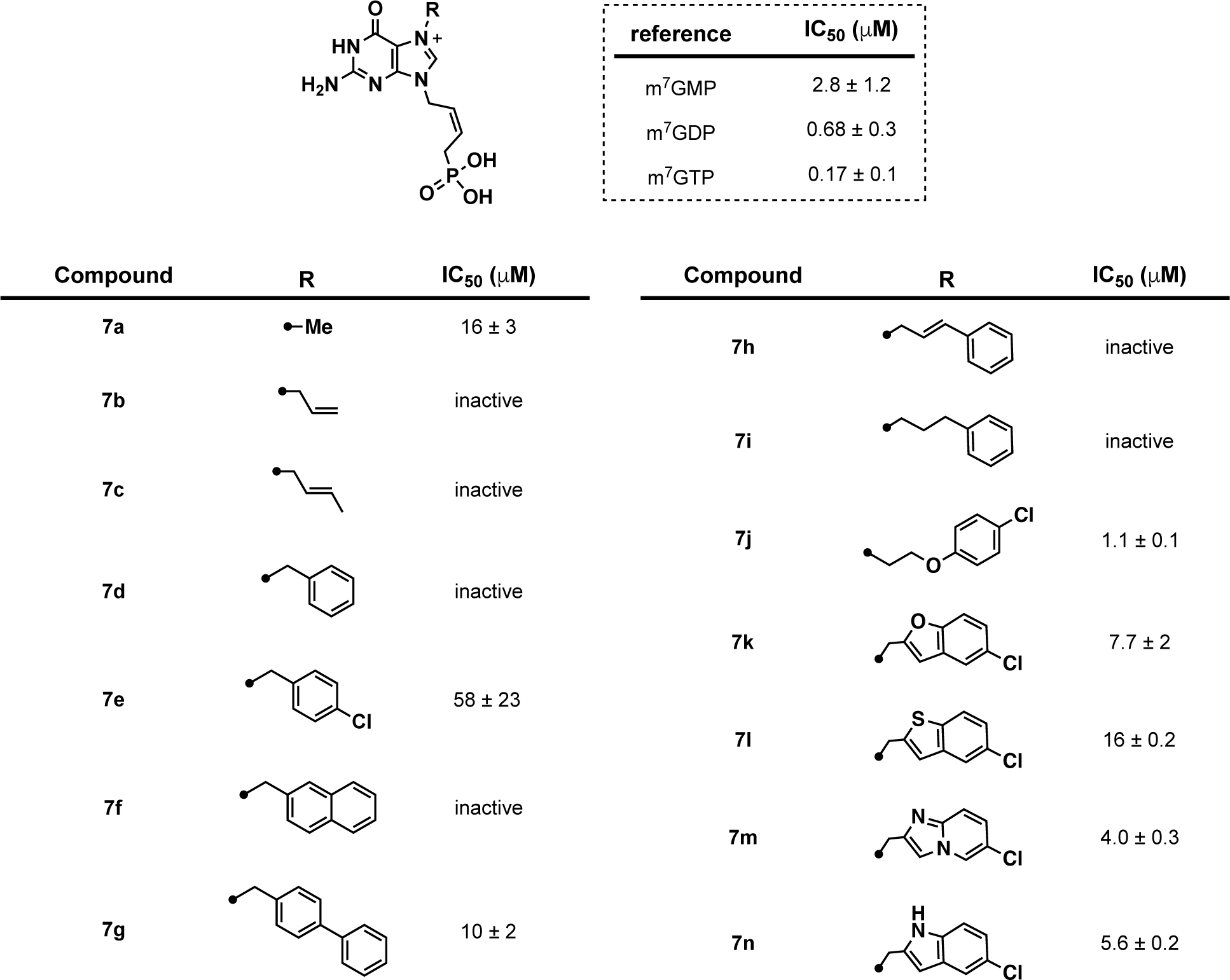
Biochemical inhibitory activities of phosphonic acids (**7**) measured using a FP assay.

Prior to cellular studies, we carried out an orthogonal eIF4E target engagement assay, the cellular thermal shift assay (CETSA).^35^ CETSA monitors the thermal stabilization of a protein induced upon binding to a ligand. To determine if cap analogues were able to engage with eIF4E in the complex cellular milieu, lysate from HeLa cells was incubated with **7j** or **7n** as models and subjected to thermal denaturation. As shown in Figure S2, both cap analogues showed significant stabilization of eIF4E indicating that these compounds can bind to eIF4E in cell lysate.

### Anti-proliferative activity correlative with inhibition of the translation of cap-dependent transcripts

Encouraged by these results, we profiled bis-POM-protected prodrugs **6j−6n** for anti-proliferative activity in MiaPaca-2 cells, a cell line whose growth is known to be affected by modulation of eIF4E and cap-dependent translation.^36, 37^ Cells were treated with compounds and anti-proliferative activity was measured using the CellTiter-Glo^®^ cell viability assay after 48 h (Figure 3). Of the compounds profiled, prodrug **6k** containing a methyl(benzofuran) substitution was found to be most potent with an EC_50_ value of 29 µM (Figures 3B and 3F). Analogues **6j**, **6l**, and **6n** exhibited similar activities with measured EC_50_ values of 51, 59, and 67 µM, respectively (Figure 3A, 3C, 3E, and 3F). Prodrug **6m** showed weak cellular activity possibility due to instability in cells (Figure 3D). Using prodrug **6n** as a model, we also demonstrated dose-dependent antiproliferative activity in additional pancreatic cancer cell lines (Figure S3). Moreover, we confirmed that these compounds do not serve as substrates of guanylate kinase, thereby prohibiting ability to serve as chain terminators analogous to anti-viral drugs (Figure S4).^38^

To provide further evidence to support on-target anti-proliferative activity in cells, we examined impact on the expression of select cap-dependent transcripts. Canonically, cap-dependent transcripts are defined as those mRNAs that contain long and highly structured 5′ untranslated regions (5′ UTR), a feature that is commonly found in many oncogenic transcripts.^2^ Examples of well-studied mRNAs whose translation is dependent upon eIF4E include c-Myc, ornithine decarboxylase 1 (ODC1), and cyclin D1, all of which play an important role in regulating cellular proliferation and become dysregulated in many cancers.^2^ Using the DepMap Portal, we confirmed that expression of these proteins is also correlative with eIF4E levels in pancreatic adenocarcinoma cell lines, including MiaPaCa-2 cells (Figure S5). Thus, we chose to measure protein levels of these cap-dependent transcripts following treatment of MiaPaCa-2 cells with **6n** (Figure 4A) using Western blot analysis. Excitingly, as shown in Figures 4B and 4C, significant and dose-dependent decreases were observed in ODC1 and cyclin D1 following treatment. A modest decrease was observed in c-Myc levels (Figure 4B), which was expected as its translation is driven by both cap-dependent and -independent processes due its unique 5′ UTR that contains an internal ribosome entry site.^39^ Overall, in combination with our target engagement studies, these results provide confidence that our design has resulted in the development of cell-permeable eIF4E inhibitors suitable for further development to validate eIF4E as a potential cancer therapeutic target.

**Figure 4.**
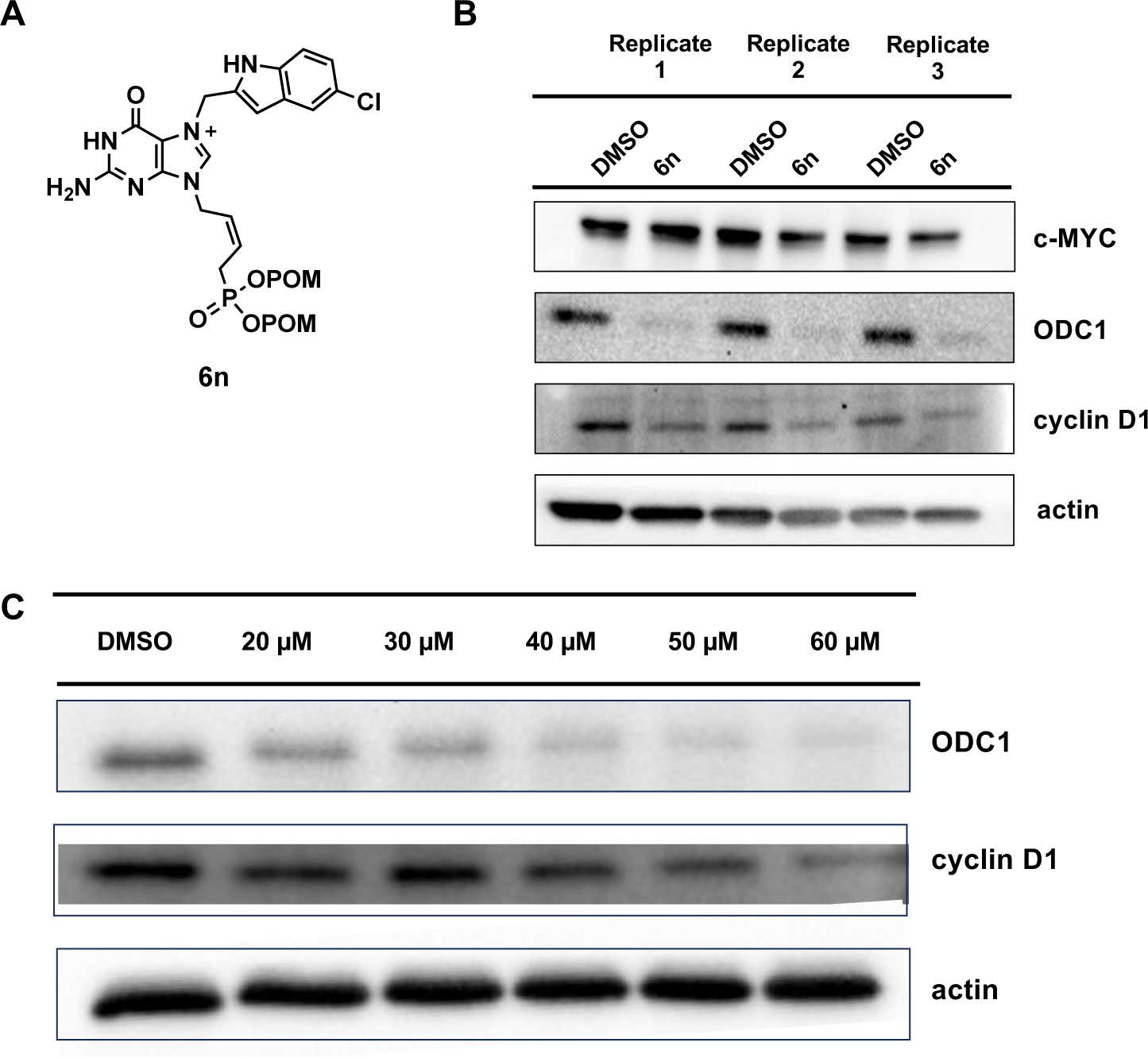
Effect of **6n** on the expression level of select cap-dependent transcripts. MiaPaCa-2 cells were treated for 6 h and protein levels were measured using Western blot. (A) Structure of **6n**. (B) Protein levels following treatment with 60 μM compound. (C) Dose-dependent inhibition of ODC1 and cyclin D1 expression.

### Crystallographic analysis of 7n bound to eIF4E

To catalyze further development of this scaffold and better understand the binding mode of our acyclic nucleoside phosphonate-based cap analogues, an X-ray crystal structure of **7n** bound to the cap-binding site of eIF4E was resolved with a resolution of 2.0 Å (Figure 5). As expected, the guanine core formed cation-π, π stacking interactions with W56 and W102, and made key hydrogen bonding interactions with the carboxylate side chain of E103 and the backbone NH group of W102. The methylene unit linking the guanine to the indole substituent formed van der Waals interactions with Trp166. The indole group of **7n** was situated in a large hydrophobic pocket lined by V153, F48, L60 and P100. W56 formed an edge-on π − π stacking interaction with the ring to complete the pocket. The chlorine atom on the indole extended further into the pocket interacting with the hydroxyl side chain of S92 (Figure 5). Interestingly, Walkinshaw and co-workers demonstrated that binding of cap analogues containing larger *N*-7-substituents, such as Bn^7^GMP, are accommodated by a conformational change that results in a 180° flip of the side chain of W102 (Figure S6A).^25^ Although **7n** accesses the same hydrophobic pocket, the *N*-7-methyl(indole) substituent is situated below the guanine core, presumably precluding the same conformational change observed in co-crystal structures with Bn^7^GMP (Figure S6B). The (*Z*)-4-phosphono-but-2-enyl linker positioned the phosphonate in proximity to the cationic residues known to interact with the triphosphate of the cap structure, resulting in formation of bifurcated hydrogen bonds between two of the phosphonate oxygens of **7n** and the side chain of R157 (Figure 5). Elucidation of the binding mode of **7n** within the cap-binding domain of eIF4E provides valuable insight for further inhibitor design of acyclic nucleoside derivatives that could take advantage of additional interactions located in the positively charged cleft accessed by the phosphonate of **7n**, as well as explore alternative non-nucleobase scaffolds. Efforts toward this goal will be reported in due course.

**Figure 5.**
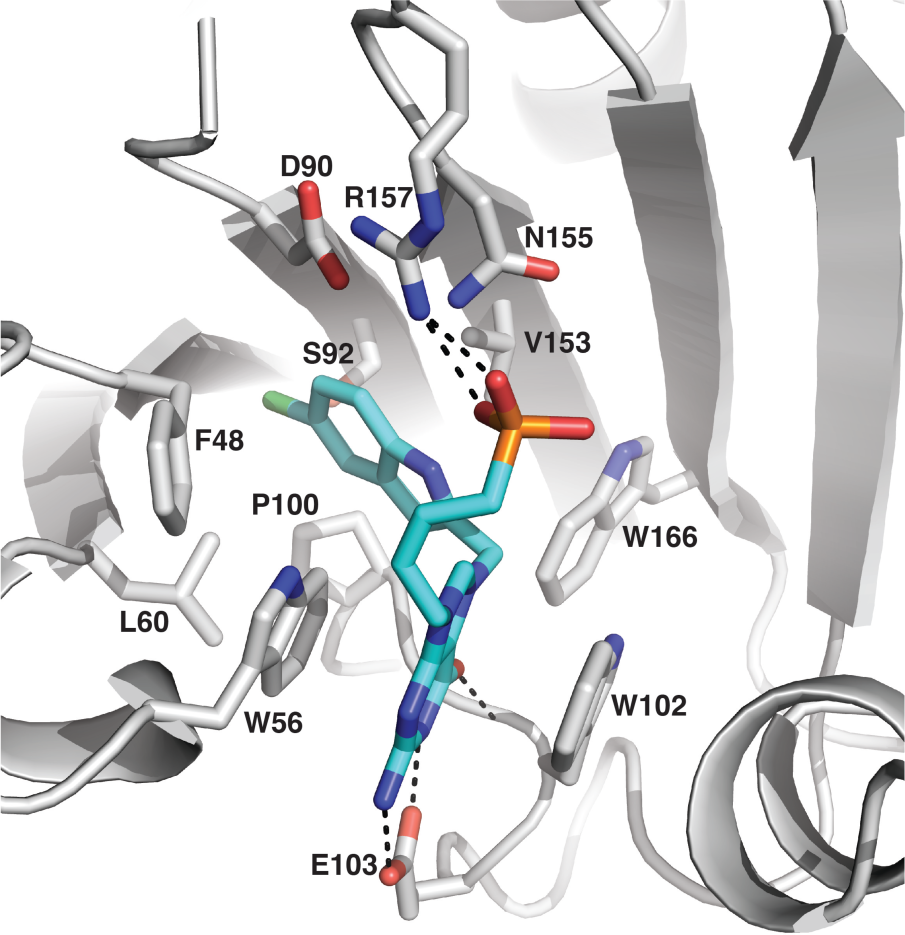
eIF4E bound to **7n**. The eIF4E protein backbone is shown as a grey ribbon with residues that interact with **7n** shown as sticks. **7n** is shown as sticks with cyan carbons. Nitrogen atoms are colored blue, oxygens in red and phosphorus in orange. Dashed lines represent hydrogen bonds. (PDB: 8SX4)

## CONCLUSIONS

Despite its well-established significance in cancer biology, eIF4E has remained an elusive target for drug discovery. Using rational design strategies, including principles gleaned from anti-viral drugs, we have developed cap-competitive acyclic nucleoside phosphonate prodrug-based inhibitors of eIF4E, resulting in the generation of cell-active compounds with promising on-target anti-proliferative activity. These compounds, in addition to recently reported small molecule eIF4E inhibitors reported by eFFECTOR Therapeutics,^40^ will provide important tools for validating eIF4E as an anti-cancer therapeutic target, as well as provide new insights into the role of cap-dependent translation in cancer and other relevant human diseases.

## EXPERIMENTAL SECTION

### General Chemistry Materials and Methods

All purchased solvents and reagents were used without further purification. Reactions were monitored by thin-layer chromatography (TLC) carried out on 0.25-mm SiliCycle silica gel plates (60F-254) using UV-light (254 nm). Flash chromatography was performed using SiliaFlash P60 silica gel. Analytical RP-HPLC was performed using an Agilent 1260 Infinity HPLC equipped with an ZORBAX Eclipse XDB-C18 colun (4.6 x 150 mm; 5 µm) with detection at 254 nm. Semi-preparative HPLC was carried out on Agilent 1260 Infinity HPLC equipped with a PrepHT XDB-C18 column (21.2 x 150 mm; 5 µm) with detection at 254 nm). NMR spectra were performed on a 300 MHz Bruker and 400 MHz Bruker instrument calibrated using a solvent peak as an internal reference. Chemical shifts (*δ* values) are reported in parts per million and are referenced to the deuterated residual solvent peak. Mass spectrometry (MS) was performed using an Agilent 6230 TOF LC/MS spectrometer using ESI ionization with an accuracy of 2 ppm. All compounds were found to be >95% pure by HPLC analysis.

*(E)-9-(but-2-en-1-yl)-6-chloro-9H-purin-2-amine (**2**)*. 2-amino-6-chloro-purine **1** (1.39 g, 8.26 mmol) was diluted in DMSO (0.66 M, 13 mL) and then treated with K_2_CO_3_ (6.15 g, 44.6 mmol) and 85% crotyl bromide (0.50 mL, 4.13 mmol) dropwise at 23 °C. After 3 h, the reaction mixture was quenched with H_2_O (10 mL) at 23 °C. The resulting reaction mixture was extracted with EtOAc (300 mL). The combined organic layers were washed with brine (200 mL) and then dried over Na_2_SO_4_. The resulting organic layer was concentrated under reduced pressure and purified by column chromatography (50% ethyl acetate/hexanes) to yield the N-9 crotyl purine **2** as a single regioisomer (805 mg, 89%). ^1^H NMR (400 MHz, MeOD-d_4_) δ 8.00 (s, 1H), 5.74 (m, 1H), 5.65 (m, 1H), 4.63 (d, *J* = 8 Hz, 2H), 1.66 (d, *J* = 4 Hz, 3H); ^13^C NMR (101 MHz, MeOD-d_4_) δ 160.10, 153.54, 150.02, 142.82, 141.28, 130.75, 124.46, 44.86, 16.47.

*((Allylphosphoryl)bis(oxy))bis(methylene) bis(2,2-dimethylpropanoate) (**3**)*. Diethylallyl phosphonate (112.2 mmol, 20 g) was diluted in CH_2_Cl_2_ (0.93 M, 120 mL) and then treated with trimethylsilyl bromide (450 mmol, 60 mL) dropwise at 23 °C. After 24 h, the mixture was concentrated to dryness, diluted in MeOH (0.3 M, 370 mL), and stirred for 1 h at 23 °C. The reaction mixture was concentrated to dryness and the crude diethylallyl phosphonic acid was used without further purification. The diethylallyl phosphonic acid was diluted in DMF (0.3 M, 225 mL) and then treated with Et_3_N (672 mmol, 93 mL), POM-Cl (672 mmol, 97 mL), and sodium iodide (11.2 mmol, 1.7 g) and warmed to 60 °C. After 5 days, the reaction was concentrated to dryness, diluted in H_2_O (300 mL), and extracted with Et_2_O (500 mL). The organic layer was dried over Na_2_SO_4_, concentrated to dryness, and purified by column chromatography (30% ethyl acetate/hexanes) to provide the desired diethylallyl bis-POM phosphonate **3** as a yellow oil (31.2 g, 83%). ^1^H NMR (400 MHz, CDCl_3_) δ 5.63 (m, 1H), 5.57 (dd, *J* = 12 Hz, 4H), 5.14 (m, 1H), 2.61 (ddt, *J* = 22, 7, 1 Hz, 2H), 1.13 (s, 18H); ^13^C NMR (101 MHz, CDCl_3_) δ 176.63, 125.94, 121.01, 81.45, 38.59, 32.78, 26.74.

*(Z)-(((4-(2-Amino-6-chloro-9H-purin-9-yl)but-2-en-1-yl)phosphoryl)bis(oxy))bis(methylene)bis (2,2-dimethylpropanoate) (**4**)*. *N*9-Crotyl purine **2** (780 mg, 3.49 mmol) was diluted in CH_2_Cl_2_ (0.04 M, 90 mL) and then subsequently treated with bis-POM allyl phosphonate (4.3 g, 12.9 mmol) and Hoveyda-Grubbs 2^nd^ Generation Catalyst (218 mg, 0.349 mmol). The resulting mixture was warmed to reflux and stirred for 48 h. The reaction mixture was cooled to 23 °C and concentrated under reduced pressure. The crude residue was loaded directly onto silica and purified by column chromatography (1−3% MeOH/CH_2_Cl_2_) to provide purine (*Z*)-bis-POM phosphonate **4** (741 mg) and (*E*)-bis-POM-phosphonate (180 mg) with a 50% yield; (*Z*) ^1^H NMR (400 MHz, CDCl_3_) δ 7.76 (s, 1H), 5.85 (m, 1H), 5.65 (m, 5H), 5.41 (bs, 2H), 4.66 (t, *J* = 5 Hz, 2H), 2.71 (dd, *J* = 24, 8 Hz, 2H), 1.20 (s, 18H); ^13^C NMR (101 MHz, CDCl_3_) δ 176.85, 159.24, 153.60, 151.26, 142.00, 129.47, 125.06, 123.69, 123.57, 81.70, 44.99, 38.70, 31.40, 30.01, 26.82; (*E*) ^1^H NMR (400 MHz, CDCl_3_) δ 7.86 (s, 1H), 5.79 (m, 1H), 5.68 (m, 5H), 5.47 (bs, 2H), 4.75 (t, *J* = 4 Hz, 2H), 2.96 (dd, *J* = 24, 8 Hz, 2H), 1.21 (s, 18H); ^13^C NMR (101 MHz, CDCl_3_) δ 176.98, 159.28, 153.67, 151.14, 142.06, 128.50, 125.02, 122.41, 122.29, 81.62, 38.73, 27.25, 26.83, 25.86; LRMS (ESI^+^) 532.1718 [M]^+^.

*(Z)-(((4-(2-amino-6-oxo-1,6-dihydro-9H-purin-9-yl)but-2-en-1yl)phosphoryl)bis(oxy))bis(methylene)bis(2,2-dimethylpropanoate) (**5**)*. Purine bis-POM phosphonate **4** (900 mg, 1.69 mmol) was diluted in an aqueous mixture of formic acid (1:1, CH_2_O_2_:H_2_O, 0.06 M, 28 mL) and then warmed to 40 °C. After 20 h, the reaction mixture was cooled to 23 °C and concentrated to dryness via lyophilization. The crude residue was purified by column chromatography (8% MeOH/CH_2_Cl_2_) to yield the guanine analog **5** (400 mg, 46%). ^1^H NMR (300 MHz, DMSO-d_6_) δ 10.55 (s, 1H), 7.60 (s, 1H), 6.44 (s, 2H), 5.85 (m, 1H), 5.57 (m, 4H), 5.43 (m, 1H), 4.55 (t, *J* = 4 Hz, 2H), 2.85 (dd, *J* = 28, 8 Hz, 2H), 1.15 (s, 18H); ^13^C NMR (101 MHz, DMSO-d_6_) δ 176.64, 157.28, 154.12, 151.43, 137.33, 131.24, 122.15, 116.97, 81.97, 44.23, 38.63, 30.72, 29.36, 26.94; LRMS (ESI^+^) 514.2056 [M]^+^.

*(Z)-2-Amino-9-(4-(bis((pivaloyloxy)methoxy)phosphoryl)but-2-en-1-yl)-7-methyl-6-oxo-6,9-dihydro-1H-purin-7-ium (**6a**)*. The guanine analogue **5** (36 mg, 0.070 mmol) was diluted in DMF (0.13 M, 0.50 mL) and then treated with iodomethane (30 mg, 0.21 mmol) dropwise at 23 °C. After 24 h, the reaction mixture was loaded directly onto silica. The crude residue was immediately purified by column chromatography (2−6% MeOH/CH_2_Cl_2_) followed by reverse phase semi−preparative HPLC (30−60% MeCN/H_2_O with 0.1% FA) to yield bis-POM cap analogue **6a** (25 mg, 68% yield). ^1^H NMR (400 MHz, DMSO-d_6_) δ 11.80 (s, 1H), 9.19 (s, 1H), 7.24 (bs, 2H), 5.89 (m, 1H), 5.61 (m, 4H), 4.75 (t, *J* = 4 Hz, 2H), 4.01 (s, 3H), 2.87 (dd, *J* = 24, 8 Hz, 2H), 1.16 (s, 18H); ^13^C NMR (101 MHz, DMSO-d_6_) δ 176.60, 156.36, 154.30, 149.87, 138.15, 128.66, 125.09, 107.79, 82.06, 46.10, 38.66, 30.90, 29.54, 26.93; LRMS (ESI^+^) 528.2204 [M]^+^.

*(Z)-2-Amino-7-benzyl-9-(4-(bis((pivaloyloxy)methoxy)phosphoryl)but-2-en-1-yl)-6-oxo-6,9-dihydro-1H-purin-7-ium (**6b**)*. Guanine analogue **5** (70 mg, 0.13 mmol) was diluted in DMSO (0.06 M, 2 mL) and then treated with benzyl bromide (0.16 mmol, 19 µL) at 23 °C for 24 h. The crude residue was immediately purified by column chromatography (2−6% MeOH/CH_2_Cl_2_) followed by reverse phase semi−preparative HPLC (30−60% MeCN/H_2_O with 0.1% FA) to yield bis-POM cap analogue **6b** (30 mg, 43%). ^1^H NMR (400 MHz, DMSO-d_6_) δ 12.23 (bs, 1H), 9.40 (s, 1H), 7.48 (m, 2H), 7.40 (m, 3H), 5.90 (m, 1H), 5.63 (s, 2H), 5.60 (m, 4H), 4.77 (t, *J* = 4 Hz, 2H), 2.85 (dd, *J* = 20, 8 Hz, 2H), 1.15 (s, 18H); ^13^C NMR (101 MHz, DMSO-d_6_) δ 177.29, 163.97, 159.00, 150.72, 139.35, 135.73, 129.94, 129.11, 128.69, 123.24, 106.98, 82.81, 51.19, 46.32, 38.64, 31.54, 29.53, 25.74; LRMS (ESI^+^) 604.3136 [M]^+^.

*(Z)-2-Amino-9-(4-(bis((pivaloyloxy)methoxy)phosphoryl)but-2-en-1-yl)-7-(4-chlorobenzyl)-6-oxo-6,9-dihydro-1H-purin-7-ium (**6c**)*. Guanine analogue **5** (70 mg, 0.058 mmol) was diluted in DMSO (0.06 M, 0.40 mL) and then treated with 4-Cl-benzyl bromide (0.23 mmol, 47 mg) at 50 °C for 24 h. The crude residue was immediately purified by column chromatography (2−6% MeOH/CH_2_Cl_2_) followed by reverse phase semi−preparative HPLC (30−60% MeCN/H_2_O with 0.1% FA) to yield bis-POM cap analogue **6c** (6.4 mg, 17%). ^1^H NMR (400 MHz, DMSO-d_6_) δ 12.23 (bs, 1H), 9.39 (s, 1H), 7.51 (d, *J* = 8 Hz, 2H), 7.47 (d, *J* = 8 Hz, 2H), 5.90 (m, 1H), 5.62 (s, 2H), 5.59 (m, 4H), 4.75 (t, *J* = 4 Hz, 2H), 2.85 (dd, *J* = 20, 8 Hz, 2H), 1.15 (s, 18H); ^13^C NMR (101 MHz, DMSO-d_6_) δ 176.58, 150.34, 137.61, 134.10, 133.86, 130.74, 129.27, 128.70, 128.54, 125.16, 125.05, 106.93, 81.98, 50.96, 45.88, 38.64, 30.88, 29.53, 26.92; LRMS (ESI^+^) 638.2128 [M]^+^.

*(Z)-7-([1,1’-Biphenyl]-4-ylmethyl)-2-amino-9-(4-(bis((pivaloyloxy)methoxy)phosphoryl)but-2-en-1-yl)-6-oxo-6,9-dihydro-1H-purin-7-ium (**6d**)*. Guanine analogue **5** (30 mg, 0.058 mmol) was diluted in DMSO (0.06 M, 1 mL) and then treated with 4-(bromomethyl)biphenyl (0.24 mmol, 60 mg) at 50 °C for 24 h. The crude residue was immediately purified by column chromatography (2−6% MeOH/CH_2_Cl_2_) followed by reverse phase semi−preparative HPLC (30−60% MeCN/H_2_O with 0.1% FA) to yield bis-POM cap analogue **6d** (6.3 mg, 16%). ^1^H NMR (400 MHz, DMSO-d6) δ 12.30 (bs, 1H), 9.48 (s, 1H), 7.66 (m, 5H), 7.58 (d, *J* = 8 Hz, 2H), 7.47 (t, *J* = 8 Hz, 2H), 7.39 (m, 2H), 5.93 (m, 1H), 5.72 (m, 1H), 5.68 (s, 2H), 5.58 (m, 1H), 4.77 (t, *J* = 4 Hz, 2H), 2.85 (dd, *J* = 24, 8 Hz, 2H), 1.14 (s, 18H); ^13^C NMR (101 MHz, DMSO-d6) δ 176.58, 150.31, 140.99, 139.94, 137.58, 134.27, 129.44, 129.42, 128.73, 128.57, 128.19, 127.59, 127.19, 125.19, 125.07, 106.98, 82.04, 81.98, 51.39, 46.33, 38.63, 30.89, 29.54, 26.91; LRMS (ESI^+^) 680.2821 [M]^+^.

*(Z)-2-Amino-9-(4-(bis((pivaloyloxy)methoxy)phosphoryl)but-2-en-1-yl)-7-(naphthalen-2-ylmethyl)-6-oxo-6,9-dihydro-1H-purin-7-ium (**6e**)*. Guanine analogue **5** (30 mg, 0.058 mmol) was diluted in DMSO (0.06 M, 1 mL) and then treated with 2-(bromomethyl)naphthalene (0.24 mmol, 64 mg) at 50 °C for 24 h. The crude residue was immediately purified by column chromatography (2−6% MeOH/CH_2_Cl_2_) followed by reverse phase semi−preparative HPLC (30−60% MeCN/H_2_O with 0.1% FA) to yield bis-POM cap analogue **6e** (4.1 mg, 11%). ^1^H NMR (400 MHz, DMSO-d6) δ 12.42 (bs, 1H), 9.41 (s, 1H), 7.94 (m, 4H), 7.61 (dd, *J* = 12, 4 Hz, 1H), 7.55 (m, 2H), 7.43 (bs, 1H), 5.91 (m, 1H), 5.80 (s, 2H), 5.68 (m, 1H), 5.57 (m, 4H), 4.76 (t, *J* = 8 Hz, 2H), 2.84 (dd, *J* = 24, 8 Hz, 2H), 1.14 (s, 18H); ^13^C NMR (101 MHz, DMSO-d6) δ 176.57, 158.02, 150.32, 137.33, 133.21, 133.19, 132.68, 129.96, 129.06, 128.39, 128.11, 127.90, 127.13, 127.06, 126.20, 125.04, 124.93, 107.16, 82.03, 81.97, 51.79, 46.27, 38.63, 30.88, 29.51, 26.90; LRMS (ESI^+^) 654.2981 [M]^+^.

*(Z)-7-allyl-2-amino-9-(4-(bis((pivaloyloxy)methoxy)phosphoryl)but-2-en-1-yl)-6-oxo-6,9-dihydro-1H-purin-7-ium (**6f**)*. Guanine analogue **5** (20 mg, 0.04 mmol) was diluted in DMSO (0.06 M, 0.70 mL) and then treated with allyl bromide (0.38 mmol, 33 µL) at 50 °C for 24 h. The crude residue was immediately purified by column chromatography (2−6% MeOH/CH_2_Cl_2_) followed by reverse phase semi−preparative HPLC (30−60% MeCN/H_2_O with 0.1% FA) to yield bis-POM cap analogue **6f** (7.8 mg, 35%). ^1^H NMR (400 MHz, DMSO-d_6_) δ 12.12 (s, 1H), 7.45 (bs, 1H), 6.09 (m, 1H), 5.91 (m, 1H), 5.69 (m, 1H), 5.59 (m, 4H), 5.34 (m, 1H), 5.04 (d, *J* = 4 Hz, 1H), 4.77 (t, *J*= 4 Hz, 2H), 2.85 (dd, *J* = 24, 8 Hz), 1.16 (s, 18H); ^13^C NMR (101 MHz, DMSO-d_6_) δ 176.59, 157.10, 154.70, 150.17, 137.49, 131.93, 128.72, 125.12, 120.46, 107.05, 82.05, 50.77, 46.26, 46.24, 38.65, 30.90, 26.93; LRMS (ESI^+^) 554.2479 [M]^+^.

*2-Amino-9-((Z)-4-(bis((pivaloyloxy)methoxy)phosphoryl)but-2-en-1-yl)-7-((E)-but-2-en-1-yl)-6-oxo-6,9-dihydro-1H-purin-7-ium (**6g**)*. Guanine analogue **5** (30 mg, 0.058 mmol) was diluted in DMSO (0.06 M, 1.0 mL) and then treated with crotyl bromide (0.23 mmol, 24 µL) at 50 °C for 24. The crude residue was immediately purified by column chromatography (2−6% MeOH/CH_2-_ Cl_2_) followed by reverse phase semi−preparative HPLC (30−60% MeCN/H_2_O with 0.1% FA) to yield bis-POM cap analogue **6g** (12.2 mg, 38%). ^1^H NMR (400 MHz, DMSO-d6) δ 12.02 (s, 1H), 9.34 (s, 1H), 7.44 (bs, 1H), 5.89 (m, 1H), 5.70 (m, 1H), 5.58 (m, 1H), 4.95 (d, *J* = 8 Hz, 1H), 4.76 (t, *J* = 4 Hz, 2H), 2.84 (dd, *J* = 24, 8 Hz, 2H), 1.70 (dd, *J* = 4, 2 Hz, 3 H), 1.16 (s, 18H); ^13^C NMR (101 MHz, DMSO-d6) δ 176.58, 156.64, 150.13, 137.26, 133.19, 131.71, 128.72, 125.15, 124.32, 123.20, 106.99, 82.05, 81.99, 50.40, 46.26, 38.64, 30.90, 29.54, 27.12, 26.92, 26.82, 17.99; LRMS (ESI^+^) 568.2518 [M]^+^.

*2-Amino-9-((Z)-4-(bis((pivaloyloxy)methoxy)phosphoryl)but-2-en-1-yl)-7-cinnamyl-6-oxo-6,9-dihydro-1H-purin-7-ium (**6h**)*. Guanine analogue **5** (30 mg, 0.058 mmol) was diluted in DMSO (0.06 M, 1.0 mL) and then treated with cinnamyl bromide (0.23 mmol, 46 mg) at 50 °C for 24 h. The crude residue was immediately purified by column chromatography (2−6% MeOH/CH_2_Cl_2_) followed by reverse phase semi−preparative HPLC (30−60% MeCN/H_2_O with 0.1% FA) to yield bis-POM cap analogue **6h** (21.3 mg, 65%). ^1^H NMR (400 MHz, DMSO-d6) δ 11.90 (bs, 1H), 9.34 (s, 1H), 7.46 (m, 2H), 7.35 (m, 4H), 6.78 (d, *J* = 24 Hz, 1H), 6.52 (m, 1H), 5.91 (m, 1H), 5.68 (m, 1H), 5.58 (m, 4H), 5.20 (d, *J* = 8 Hz, 2H), 4.76 (t, *J* = 4 Hz, 2H), 2.84 (dd, *J* = 28, 12 Hz, 2H), 1.15 (s, 18H); ^13^C NMR (101 MHz, DMSO-d6) δ 176.57, 160.70, 142.94, 140.13, 135.87, 131.37, 129.33, 129.18, 128.25, 128.19, 127.42, 127.26, 127.16, 82.00, 76.33, 70.24, 69.70, 51.40, 38.66, 26.91; LRMS (ESI^+^) 630.2707 [M]^+^.

*(Z)-2-amino-9-(4-(bis((pivaloyloxy)methoxy)phosphoryl)but-2-en-1-yl)-6-oxo-7-(3-phenylpropyl)-6,9-dihydro-1H-purin-7-ium (**6i**)*. Guanine analogue **5** (200 mg, 0.38 mmol) was diluted in DMF (0.09 M, 4 mL) and then treated with 3-phenylpropyl bromide (1.16 mmol, 177 µL) at 70 °C for 24 h. The crude residue was immediately purified by column chromatography (2−6% MeOH/CH_2_Cl_2_) followed by reverse phase semi−preparative HPLC (30−60% MeCN/H_2_O with 0.1% FA) to yield bis-POM cap analogue **6i** (7.0 mg, 31%). ^1^H NMR (400 MHz, DMSO-d6) δ 11.97 (bs, 1H), 9.31 (s, 1H), 7.43 (bs, 2H), 7.26 (m, 2H), 7.20 (m, 3H), 5.90 (m, 1H), 5.71 (m, 1H), 5.61 (m, 4H), 4.75 (d, *J* = 24 Hz, 2H), 4.41 (t, *J* = 8 Hz, 2H), 2.66 (t, *J* = 8 Hz, 2H), 2.22 (m, 2H), 1.15 (s, 18H); ^13^C NMR (101 MHz, DMSO-d6) δ 176.59, 156.87, 153.43, 150.17, 140.96, 137.57, 128.75, 128.63, 126.44, 124.89, 107.21, 82.05, 81.99, 49.11, 46.15, 43.47, 38.64, 32.12, 30.96, 27.50, 26.92; LRMS (ESI^+^) 632.3069 [M]^+^.

*(Z)-2-Amino-9-(4-(bis((pivaloyloxy)methoxy)phosphoryl)but-2-en-1-yl)-7-(2-(4-chlorophenoxy)ethyl)-6-oxo-6,9-dihydro-1H-purin-7-ium (**6j**)*. Guanine analogue **5** (60 mg, 0.116 mmol) was diluted in DMF (0.06 M, 2 mL) and then treated with 1-(2-bromoethoxy)-4-chlorobenzene (0.584 mmol, 136 mg) at 70 °C for 24 h. The crude residue was immediately purified by column chromatography (2−6% MeOH/CH_2_Cl_2_) followed by reverse phase semi−preparative HPLC (30−60% MeCN/H_2_O with 0.1% FA) to yield bis-POM cap analogue **6j** (2.7 mg, 4%). ^1^H NMR (400 MHz, DMSO-d6) δ 10.86 (s, 1H), 9.25 (s, 1H), 7.32 (d, *J* = 8 Hz, 2H), 6.58 (d, *J* = 8 Hz, 2H), 6.52 (bs, 2H), 5.86 (m, 1H), 5.60 (m, 5H), 4.78 (t, *J* = 8 Hz, 2H), 4.56 (t, *J* = 8 Hz, 2H), 4.43 (t, *J* = 8 Hz, 2H), 2.80 (dd, *J* = 24, 8 Hz, 2H), 1.15 (s, 18H); ^13^C NMR (101 MHz, DMSO-d6) δ 176.65, 157.41, 157.05, 154.19, 151.48, 137.35, 131.31, 129.75, 125.43, 122.20, 116.96, 107.13, 81.97, 66.12, 55.38, 44.23, 38.64, 30.72, 29.36, 26.94; LRMS (ESI^+^) 668.2229 [M]^+^.

*(Z)-2-Amino-9-(4-(bis((pivaloyloxy)methoxy)phosphoryl)but-2-en-1-yl)-7-((5-chlorobenzofuran-2-yl)methyl)-6-oxo-6,9-dihydro-1H-purin-7-ium (**6k**)*: Guanine analogue **5** (30 mg, 0.058 mmol) was diluted in DMSO (0.06 M, 1 mL) and then treated with 2-(bromomethyl)-5-chlorobenzofuran (0.24 mmol, 60 mg) at 50 °C to 24 h. The crude residue was immediately purified by column chromatography (2−6% MeOH/CH_2_Cl_2_) followed by reverse phase semi−preparative HPLC (30−60% MeCN/H_2_O with 0.1% FA) to yield bis-POM cap analogue **6k** (7.4 mg, 24%). ^1^H NMR (500 MHz, DMSO-d6) δ 12.10 (s, 1H), 9.37 (s, 1H), 7.77 (d, *J* = 1 Hz, 1H), 7.61 (d, *J* = 10 Hz, 1H), 7.37 (m, 3H), 7.08 (s, 1H), 6.83 (d, *J* = 10 Hz, 1H), 5.89 (m, 1H), 5.86 (s, 2H), 5.67 (m, 1H), 5.58 (m, 1H), 4.76 (t, *J* = 8 Hz, 2H), 2.83 (dd, *J* = 20, 8 Hz, 2H), 1.14 (s, 18H); ^13^C NMR (126 MHz, DMSO-d6) δ 176.58, 158.45, 151.70, 133.43, 130.36, 125.56, 122.60, 122.48, 119.84, 112.55, 107.54, 81.96, 72.72, 50.99, 38.63, 30.37, 27.20, 26.95, 26.90; LRMS (ESI^+^) 678.2070 [M]^+^.

*(Z)-2-Amino-9-(4-(bis((pivaloyloxy)methoxy)phosphoryl)but-2-en-1-yl)-7-((5-chlorobenzo[b] thiophen-2-yl)methyl)-6-oxo-6,9-dihydro-1H-purin-7-ium (**6l**)*. Guanine analogue **5** (30 mg, 0.058 mmol) was diluted in DMSO (0.06 M, 1 mL) and then treated with 2-(bromomethyl)-5-chlorobenzo[*b*]thiophene (0.175 mmol, 46 mg) at 50 °C for 24 h. The crude residue was immediately purified by column chromatography (2−6% MeOH/CH_2_Cl_2_) followed by reverse phase semi−preparative HPLC (30−60% MeCN/H_2_O with 0.1% FA) to yield bis-POM cap analogue **6l** (8.8 mg, 22%). ^1^H NMR (400 MHz, DMSO-d6) δ 12.12 (s, 1H), 9.49 (s, 1H), 8.02 (d, *J* = 8 Hz, 1H), 7.98 (d, *J* = 4 Hz, 1H), 7.59 (s, 1H), 7.41 (dd, *J* = 8, 4 Hz, 1H), 5.95 (s, 2H), 5.89 (m, 1H), 5.68 (m, 1H), 5.57 (m, 4H), 4.78 (t, *J* = 8 Hz, 2H), 2.84 (dd, *J* = 24, 8 Hz), 1.14 (s, 18H); ^13^C NMR (101 MHz, DMSO-d6) δ 176.57, 150.14, 140.60, 139.90, 138.76, 138.03, 130.28, 128.41, 125.61, 125.24, 124.85, 123.88, 106.87, 82.03, 47.18, 38.63, 30.87, 29.51, 26.90; LRMS (ESI^+^) 694.1851 [M]^+^.

*(Z)-2-Amino-9-(4-(bis((pivaloyloxy)methoxy)phosphoryl)but-2-en-1-yl)-7-((6-chloroimidazo[1,2-a]pyridin-2-yl)methyl)-6-oxo-6,9-dihydro-1H-purin-7-ium (**6m**)*. Guanine analogue **5** (30 mg, 0.058 mmol) was diluted in DMSO (0.06 M, 1 mL) and then treated with 2-(bromomethyl)-6-chloroimidazo[1,2-*a*]pyridine (0.175 mmol, 43 mg) at 50 °C for 24 h. The crude residue was immediately purified by column chromatography (2−6% MeOH/CH_2_Cl_2_) followed by reverse phase semi−preparative HPLC (30−60% MeCN/H_2_O with 0.1% FA) to yield bis-POM cap analogue **6m** (7.7 mg, 20%). ^1^H NMR (400 MHz, DMSO-d6) δ 9.11 (s, 1H), 8.88 (s, 1H), 8.07 (s, 1H), 7.75 (d, *J* = 12 Hz, 1H), 7.32 (dd, *J* = 16, 4 Hz, 1H), 5.89 (m, 1H), 5.76 (s, 2H), 5.58, (m, 4H), 5.40 (m, 1H), 4.73 (t, *J* = 4 Hz, 2H), 2.83 (dd, *J* = 24, 8 Hz, 2H), 1.14 (s, 18H); ^13^C NMR (101 MHz, DMSO-d6) δ 176.56, 157.00, 150.07, 143.44, 140.19, 137.19, 126.82, 125.69, 125.05, 119.82, 118.04, 113.39, 107.15, 82.04, 55.40, 46.51, 38.62, 30.85, 29.49, 26.90; LRMS (ESI^+^) 678.2300 [M]^+^.

*(Z)-2-Amino-9-(4-(bis((pivaloyloxy)methoxy)phosphoryl)but-2-en-1-yl)-7-((5-chloro-1H-indol-2-yl)methyl)-6-oxo-6,9-dihydro-1H-purin-7-ium (**6n**)*. Guanine analogue **5** (50 mg, 0.097 mmol) was diluted in DMSO (0.06 M, 2 mL) and then treated with *tert*-butyl 2-(bromomethyl)-5-chloro-1*H*-indole-1-carboxylate (0.485 mmol, 166 mg) at 50 °C for 24 h. The crude residue was immediately purified by column chromatography (2−6% MeOH/CH_2_Cl_2_) to yield the desired cap analogue precursor (30 mg, 40%). ^1^H NMR (400 MHz, MeOH-d4) δ 8.05 (d, *J* = 8 Hz, 1H), 7.53 (d, *J* = 4 Hz, 1H), 7.31 (dd, *J* = 12, 4 Hz, 1H), 6.70 (s, 1H), 6.08 (s, 2H), 5.86 (m, 1H), 5.63 (m, 4H), 5.53 (m, 1H), 2.85 (dd, *J* = 24, 8 Hz, 2H), 1.71 (s, 9H), 1.20 (s, 18H); ^13^C NMR (101 MHz, MeOH-d4) δ 176.80, 166.66, 157.34, 154.93, 149.95, 134.99, 134.08, 129.71, 128.52, 125.50, 124.83, 120.13, 116.72, 110.44, 85.87, 81.81, 38.32, 30.47, 29.15, 27.00, 25.84; LRMS (ESI^+^) 777.2751 [M]^+^.

The cap analog precursor (30 mg, 0.038 mmol) was diluted in CH_2_Cl_2_ (300 µL) and then treated with TFA (145 µL) at 23 °C. After 45 minutes, the reaction mixture was concentrated to dryness. The crude residue was then purified by reverse phase semi−preparative HPLC (30−60% MeCN/H_2_O with 0.1% FA) to yield bis-POM cap analogue **6n** (4.7 mg, 39% yield). ^1^H NMR (400 MHz, DMSO-d6) δ 12.17 (s, 1H), 9.05 (s, 1H), 7.58 (d, *J* = 4Hz, 1H), 7.41 (d, *J* = 8 Hz, 1H), 7.10 (dd, *J* = 8, 4 Hz, 1H), 6.53 (s, 1H), 6.38 (bs, 2H), 5.88 (m, 1H), 5.77 (s, 2H), 5.66 (m, 1H), 5.58 (m, 4H), 4.71 (t, *J* = 4 Hz, 2H), 2.82 (dd, *J* = 24, 8 Hz, 2H), 1.13 (s, 18H); ^13^C NMR (101 MHz, DMSO-d6) δ 176.58, 156.62, 153.64, 150.11, 135.86, 135.16, 133.62, 128.05, 125.22, 124.41, 120.66, 113.61, 86.18, 82.02, 46.45, 38.63, 30.60, 29.43, 26.89; LRMS (ESI^+^) 677.2249 [M]^+^.

*(Z)-2-Amino-7-methyl-6-oxo-9-(4-phosphonobut-2-en-1-yl)-6,9-dihydro-1H-purin-7-ium (**7a**)*. Bis-POM cap analogue **6a** (30 mg, 0.056 mmol) was diluted in MeOH (0.21 M, 7 µL) and 5N HCl_(aq)_ (0.014 M, 1mL) at 23 °C. The resulting reaction mixture was warmed to 45 °C and stirred for 24 h. The reaction mixture was then cooled to 23 °C and diluted with H_2_O (20 mL). The mixture was frozen and concentrated via lyophilization. The crude reside was then purified by reverse-phase semi-preparative HPLC (5−80% MeCN with 0.1% FA) to yield **7a** (4.7 mg, 29%). LRMS (ESI^+^) 300.0847 [M]^+^.

*(Z)-2-Amino-7-benzyl-6-oxo-9-(4-phosphonobut-2-en-1-yl)-6,9-dihydro-1H-purin-7-ium (**7b**)*. Bis-POM cap analogue **6b** (10 mg, 0.016 mmol) was diluted in MeOH (0.21 M, 70 µL) and 5 N HCl_(aq)_ (0.014 M, 1 mL) and warmed to 45 °C for 24 h. The crude reside was then purified by reverse-phase semi-preparative HPLC (5−80% MeCN with 0.1% FA) to provide cap analogue **7b** (3 mg, 50%). LRMS (ESI^+^) 376.1178 [M]^+^.

*(Z)-2-Amino-7-(4-chlorobenzyl)-6-oxo-9-(4-phosphonobut-2-en-1-yl)-6,9-dihydro-1H-purin-7-ium (**7c**)*. Bis-POM cap analogue **6c** (30 mg, 0.047 mmol) was diluted in MeOH (0.21 M, 0.22 mL) and 5 N HCl_(aq)_ (0.014 M, 4 mL) and warmed to 45 °C for 24 h. The crude reside was then purified by reverse-phase semi-preparative HPLC (5−80% MeCN with 0.1% FA) to provide cap analogue **7c** (1 mg, 3%), LRMS (ESI^+^) 410.0781 [M]^+^.

*(Z)-7-([1,1’-Biphenyl]-4-ylmethyl)-2-amino-6-oxo-9-(4-phosphonobut-2-en-1-yl)-6,9-dihydro-1H-purin-7-ium (**7d**)*. Bis-POM cap analogue **6d** (15 mg, 0.022 mmol) was diluted in MeOH (0.21 M, 0.10 mL) and 5 N HCl_(aq)_ (0.014 M, 2 mL) and warmed to 45 °C for 24 h. The crude reside was then purified by reverse-phase semi-preparative HPLC (5−80% MeCN with 0.1% FA) to provide cap analogue **7d** (1.5 mg, 15%). LRMS (ESI^+^) 452.0517 [M]^+^.

*(Z)-2-Amino-7-(naphthalen-2-ylmethyl)-6-oxo-9-(4-phosphonobut-2-en-1-yl)-6,9-dihydro-1H-purin-7-ium (**7e**)*. Bis-POM cap analogue **6e** (10 mg, 0.015 mmol) was diluted in MeOH (0.21 M, 71 µL) and 5 N HCl_(aq)_ (0.014 M, 1 mL) and warmed to 45 °C for 24 h. The crude reside was then purified by reverse-phase semi-preparative HPLC (5−80% MeCN with 0.1% FA) to provide cap analogue **7e** (1.0 mg, 16%). LRMS (ESI^+^) 426.1353 [M]^+^.

*(Z)-7-Allyl-2-amino-6-oxo-9-(4-phosphonobut-2-en-1-yl)-6,9-dihydro-1H-purin-7-ium (**7f**)*. Bis-POM cap analogue **6f** (20 mg, 0.036 mmol) was diluted in MeOH (0.21 M, 0.20 mL) and 5 N HCl_(aq)_ (0.014 M, 3 mL) and warmed to 45 °C for 24 h. The crude reside was then purified by reverse-phase semi-preparative HPLC (5−80% MeCN with 0.1% FA) to provide cap analogue **7f** (2.3 mg, 21%). LRMS (ESI^+^) 326.0195 [M]^+^.

*2-Amino-7-((E)-but-2-en-1-yl)-6-oxo-9-((Z)-4-phosphonobut-2-en-1-yl)-6,9-dihydro-1H-purin-7-ium (**7g**)*. Bis-POM cap analogue **6g** (35 mg, 0.061 mmol) was diluted in MeOH (0.21 M, 0.30 mL) and 5 N HCl_(aq)_ (0.014 M, 3 mL) and warmed to 45 °C for 24 h. The crude reside was then purified by reverse-phase semi-preparative HPLC (5−80% MeCN with 0.1% FA) to provide cap analogue **7g** (1.0 mg, 8%). LRMS (ESI^+^) 340.0335 [M]^+^.

*2-Amino-7-cinnamyl-6-oxo-9-((Z)-4-phosphonobut-2-en-1-yl)-6,9-dihydro-1H-purin-7-ium (**7h**)*. Bis-POM cap analogue **6h** (35 mg, 0.079 mmol) was diluted in MeOH (0.21 M, 0.30 mL) and 5 N HCl_(aq)_ (0.014 M, 6 mL) and warmed to 45 °C for 24 h. The crude reside was then purified by reverse-phase semi-preparative HPLC (5−80% MeCN with 0.1% FA) to provide cap analogue **7h** (1.0 mg, 3%). LRMS (ESI^+^) 402.1327 [M]^+^.

*(Z)-2-Amino-6-oxo-7-(3-phenylpropyl)-9-(4-phosphonobut-2-en-1-yl)-6,9-dihydro-1H-purin-7-ium (**7i**)*. Bis-POM cap analogue **6i** (30 mg, 0.047 mmol) was diluted in MeOH (0.2 M, 0.22 mL) and 5 N HCl_(aq)_ (0.014 M, 4 mL) and warmed to 45 °C for 24 h. The crude reside was then purified by reverse-phase semi-preparative HPLC (5−80% MeCN with 0.1% FA) to provide cap analogue **7i** (1.3 mg, 7%). LRMS (ESI^+^) 404.0568 [M]^+^.

*(Z)-2-Amino-7-(2-(4-chlorophenoxy)ethyl)-6-oxo-9-(4-phosphonobut-2-en-1-yl)-6,9-dihydro-1H-purin-7-ium (**7j**)*. Bis-POM cap analogue **6j** (6 mg, 8.90 µmol) was diluted in MeOH (0.2 M, 40 µL) and 5 N HCl_(aq)_ (0.014 M, 1 mL) and warmed to 45 °C for 24 h. The crude reside was then purified by reverse-phase semi-preparative HPLC (5−80% MeCN with 0.1% FA) to provide cap analogue **7j** (2.0 mg, 51%). LRMS (ESI^+^) 440.0874 [M]^+^.

*(Z)-2-Amino-7-((5-chlorobenzofuran-2-yl)methyl)-6-oxo-9-(4-phosphonobut-2-en-1-yl)-6,9-dihydro-1H-purin-7-ium (**7k**)*. Bis-POM cap analogue **6k** (30 mg, 0.044 mmol) was diluted in MeOH (0.2 M, 0.20 mL) and 5 N HCl_(aq)_ (0.014 M, 5 mL) and warmed to 45 °C for 24 h. The crude reside was then purified by reverse-phase semi-preparative HPLC (5−80% MeCN with 0.1% FA) to provide cap analogue **7k** (5.5 mg, 29%). LRMS (ESI^+^) 450.0717 [M]^+^.

*(Z)-2-Amino-7-((5-chlorobenzo[b]thiophen-2-yl)methyl)-6-oxo-9-(4-phosphonobut-2-en-1-yl)-6,9-dihydro-1H-purin-7-ium (**7l**)*. Bis-POM cap analogue **6l** (40 mg, 0.057 mmol) was diluted in MeOH (0.2 M, 0.30 mL) and 5 N HCl_(aq)_ (0.014 M, 4 mL) and warmed to 45 °C for 24 h. The crude reside was then purified by reverse-phase semi-preparative HPLC (5−80% MeCN with 0.1% FA) to provide cap analogue **7l** (2.5 mg, 10%). LRMS (ESI^+^) 466.0488 [M]^+^.

*(Z)-2-Amino-7-((6-chloroimidazo[1,2-a]pyridin-2-yl)methyl)-6-oxo-9-(4-phosphonobut-2-en-1-yl)-6,9-dihydro-1H-purin-7-ium (**7m**)*. Bis-POM cap analogue **6m** (8 mg, 0.011 mmol) was diluted in MeOH (0.2 M, 50 µL) and 5 N HCl_(aq)_ (0.014 M, 2 mL) and warmed to 45 °C for 24 h. The crude reside was then purified by reverse-phase semi-preparative HPLC (5−80% MeCN with 0.1% FA) to provide cap analogue **7m** (1.9 mg, 38%). LRMS (ESI^+^) 450.0814 [M]^+^.

*(Z)-2-Amino-7-((5-chloro-1H-indol-2-yl)methyl)-6-oxo-9-(4-phosphonobut-2-en-1-yl)-6,9-dihydro-1H-purin-7-ium (**7n**)*. Bis-POM cap analogue **6n** (43 mg, 0.08 mmol) was diluted in MeOH (0.2 M, 0.4 mL) and 5 N HCl_(aq)_ (0.014 M, 6 mL) and warmed to 45 °C for 24 h. The crude reside was then purified by reverse-phase semi-preparative HPLC (5−80% MeCN with 0.1% FA) to provide cap analogue **7n** (9.6 mg, 27%). LRMS (ESI^+^) 449.0831 [M]^+^.

### General Biology Materials

EDA-m7GTP-5-FAM (NU-824-5FM) was purchased from Jena Bioscience and used as received. The following antibodies were used in this study: eIF4E (Cell Signaling Technology, 9742), eIF4G (Cell Signaling Technology, 2498), actin-HRP (Santa Cruz Biotechnology, sc-47778), c-Myc (Cell Signaling Technology, 13987), ODC1 (Novus Biologicals, 2878R), and cyclin D1 (Cell Signaling Technology, 2978).

### Data and Statistical Analysis

All data was analyzed using GraphPad Prism version 9.5.1 for Mac OS (GraphPad Software, www.graphpad.com). Graphs show mean ± standard deviation as described in the figure legends.

### Protein Expression and Purification

Human eIF4E subcloned into a pET19b vector modified to include a 10× His tag and a PreScission protease cut-site between the tag and the beginning of the eIF4E gene was used for expression.^41^ pET19bpp-eIF4E plasmid was transformed into BL21(DE3) cells. Cells were grown at 37 °C to an OD_600_ of 0.6−0.8, induced with 1 mM IPTG, and grown for 16 h at 18 °C. The cells were pelleted and lysed through sonication in lysis buffer (50 mM Tris-HCl, pH 8, 500 mM NaCl, 50 mM imidazole, 20 mM β-mercaptoethanol, 5 mM DTT, and 1% Tween-20 with protease inhibitors). The cell lysate was centrifuged at 38,000g for 2 h and then incubated with Ni-NTA resin for 1 h at 4 °C. The resin was washed with 15 mL of lysis buffer (3−4×, 15 min each, in 50 mM Tris-HCl, pH 8, 500 mM NaCl, 25 mM imidazole, 5 mM DTT). eIF4E was eluted with 5-mL aliquots (total ∼ 25 mL) of elution buffer (10 mM Tris-HCl, pH 8, 500 mM NaCl, 100 mM imidazole, 5 mM DTT). Protein was then dialyzed overnight at 4 °C in dialysis buffer (20 mM Tris buffer, pH 7.4, 100 mM NaCl, 2 mM DTT). After dialysis, protein was concentrated and purified using a FPLC Superdex 75 16/10 preparative column. Pure protein was aliquoted and stored in −80 °C. The yield was ∼1 mg from 1 L of cell culture.

### FP Assay

FP measurements were performed on a Biotek Cytation 3 microplate reader equipped with excitation (485 ± 20 nm) and emission (528 ± 20 nm) polarization filters. Experiments were carried out at room temperature in 384-well non-binding low volume black microplates with sample volume of 20 μL per well. Each condition was done in quadruplicate and each experiment in triplicate. eIF4E FP buffer was 50 mM HEPES (pH 7.2), 100 mM KCl, 0.5 mM EDTA, 1 mM DTT, 0.0025% Tween 20 and 0.05 mg/mL BSA. To determine K_d_ values, 15 nM EDA-m7GTP-5-FAM was mixed with increasing concentration of His10-eIF4E (0−10 μM). Before FP measurements, the plate was shaken at 150 rpm at room temperature for 40 min. For competition experiments, a mixture containing 15 nM probe and 95 nM protein in FP buffer was incubated with the increasing concentrations of compound **7** (0−80 μM). Plates were shaken for 40 min at room temperature and FP was measured. Values were plotted using GraphPad Prism.

### General Cell Culture

MiaPaca-2 and HeLa cell lines were grown in DMEM, 10% FBS, 2 mM L-glutamine at 37 °C with 5% CO_2_ in a humidified incubator.

### CETSA and Western Blot

Cultured HeLa were trypsinized and washed with PBS. Cells were then diluted in PBS supplemented with protease inhibitors. Cell suspensions were freeze-thawed 3× using liquid nitrogen. The soluble fraction (lysate) was separated from the cell debris by centrifugation at 15,000 rpm for 25 min at 4°C. Cell lysates (150 μg total protein) were then divided and treated with either control (water) or compounds **7** (100 μM). After 1 h incubation at room temperature, lysates were divided into 90-μL aliquots and heated at 42 °C, 49 °C, or 52 °C for 3 min followed by cooling at 20 °C for 3 min. Heated lysates were centrifuged at 15,000 rpm for 25 min at 4°C to separate the soluble fractions from precipitates. Supernatants were transferred to new tubes and analyzed by SDS-PAGE. Samples were run on a 4−12% Bis-Tris gel and transferred to PVDF membrane in Towbin’s Buffer. The membrane was blocked in 5% milk for 1 h at 25 °C, and then incubated with a primary antibody (overnight at 4 °C) and secondary antibody (1 h at 25 °C). Proteins were visualized using a Biorad ChemiDoc imaging system.

### Densitometry

Scanned Western blot images were processed using ImageJ software. The pixels in each band gave a raw reading. The baseline was subtracted from each raw reading. Raw readings were normalized to actin, and then ratios of treatment to control were calculated. Graphs were plotted in GraphPad Prism.

### Cell Viability Assay

The Cell Titer-Glo^®^ assay kit was purchased from Promega and performed according to the manufacturer’s instructions. Briefly, 2,000 MiaPaCa-2 cells were plated in a white, 96-well tissue culture-treated plate. Cells were treated with prodrugs **6** in quadruplicate and incubated for 48 h. After 48 h, the cell culture media was replaced with 70 µl of OptiMEM and then lysed with 70 µl of Cell Titer-Glo^®^ reagent. Total luminescence was read within 1 h using a BioTek Cytation 3 reader. Data was normalized and processed in GraphPad Prism.

### Total Cell Lysate and Western Blot

MiaPaCa-2 cells were grown in a 6-well plate to 50% confluency. After 16−24 h, cells were treated with 60 μM of compound (the EC_50_ determined from dose-response studies) or DMSO control and incubated for 6 h. After 6 h, cells were harvested in 200−350 μL of RIPA buffer (10 mM Tris-HCl, 150 mM NaCl, 1% Triton, 1% sodium deoxycholate, 0.1% SDS, pH 7.2). Total protein was quantified using the BCA assay and analyzed by Western blot.

### eIF4E expression and purification for crystallography

The gene for eIF4E (residues 27−217) was cloned into a pET19b expression vector and transformed into Rosetta^TM^2 cells. The protein was expressed and purified as described in Papadopoulos *et al*.^42^ with the following modification. Supernatant was incubated with m^7^GDP cap resin^24^ overnight and eluted the following day. After each elution, the protein was buffer exchanged into 10 mM HEPES, pH 7.5, 125 mM NaCl, and 1 mM TCEP and concentrated to 0.5−1.0 mg/mL and flash frozen in liquid nitrogen.

### Crystallization and Structure Determination

For crystallization, the protein was concentrated to 4.6 mg/mL. Native crystals grew from drops containing 2 μL protein and 2 μL well solution (16−26% PEG 3350, 0.1 M MES pH 6.0, 10% isopropanol, and 2 mM CaCl_2_). Small, crystals appeared after 1 week and continued to grow over the next few weeks. To achieve the inhibitor bound structure, powdered compound **7n** was added directly to the well containing crystals and incubated overnight. The following day, the crystals were cryoprotected in a solution of 26% PEG 3350, 0.1M MES pH 6.0, 2mM CaCl_2_, and 20% glycerol, and flash frozen for data collection. Diffraction data were collected at beamline 21-ID-D at the Advance Photon Source and processed with HKL2000.^43^ The structure was solved by molecular replacement in Phaser^44^ using the protein component of 4TQC as a search model. Iterative rounds of model building and refinement were completed using Coot^45^ and Buster.^46^ Coordinates and geometric restraints for the ligand and inhibitor were created using Grade.^46^ The crystals grew in the P2 space group with two molecules in the asymmetric unit. After molecular replacement, difference electron density corresponding to **7n** was observed in the binding site of the A chain, while density for m^7^GDP was observed in the B chain (Figure S7). In chain A, there was clear density for residues 33−217, except for the disordered region containing residues 83−87 and 208−211. In the B chain, residues 30−217 were apparent in electron density, except for residues 208−209. The structure was validated using Molprobity^47^ and ligand statistics were obtained from the PBD validation server. Data refinement statistics are given in Table S1.

## ASSOCIATED CONTENT

### Supporting Information

The Supporting Information is available free of charge at…

Supplemental figures, ^1^H and ^13^C NMR spectra, HPLC spectra, and crystallography data collection and refinement statistics (PDF)

Molecular formula strings (CSV)

### Accession Codes

eIF4E bound to **7n**, PDB: 8SX4

## ACKNOWLEDGMENTS

This work was supported by the NIH (R01 CA202018 and R01 GM132341 to A.L.G.) and the Rogel Cancer Center. This research used resources of the Advanced Photon Source, a U.S. Department of Energy (DOE) Office of Science User Facility operated for the DOE Office of Science by Argonne National Laboratory under Contract No. DE-AC02-06CH11357. Use of the LS-CAT Sector 21 was supported by the Michigan Economic Development Corporation and the Michigan Technology Tri-Corridor (Grant 085P1000817). This study is supported, in part, by the Cancer Center Support Grant (P30CA046592) from the National Cancer Institute, National Institutes of Health, to Rogel Cancer Center of the University of Michigan. Figure 1 was created using Biorender.

## ABBREVIATIONS

eIF4E: eukaryotic translation initiation factor 4E
POM: pivaloyloxymethyl
FP: fluorescence polarization
ODC1: ornithine decarboxylase 1
UTR: untranslated region.

## TABLE OF CONTENTS GRAPHIC

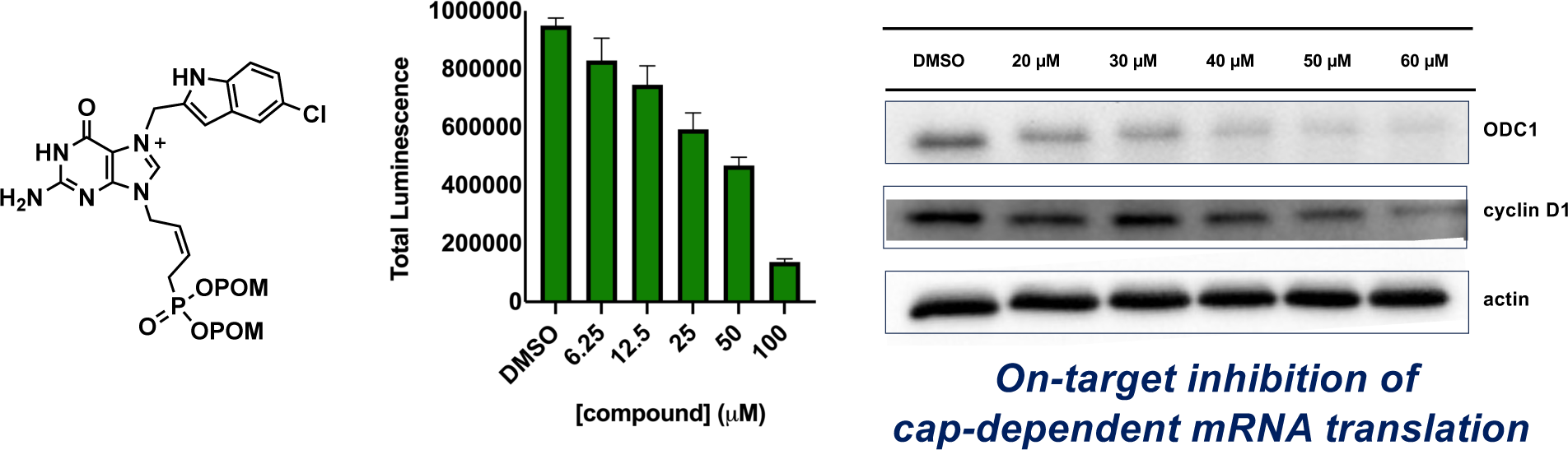

