## Supporting Information for "Design of Cell-Permeable Inhibitors of Eukaryotic Translation Initiation Factor 4E (eIF4E) for Inhibiting Aberrant Cap-Dependent Translation in Cancer"

**A. Supplemental Figures** Pages S2−S5

**B. $^{1}$H and $^{13}$C NMR Spectra** Pages S6−S24

**C. HPLC Spectra** Pages S25−S34

**D. X-ray Crystallography Data** Page S35

### A. Supplemental Figures

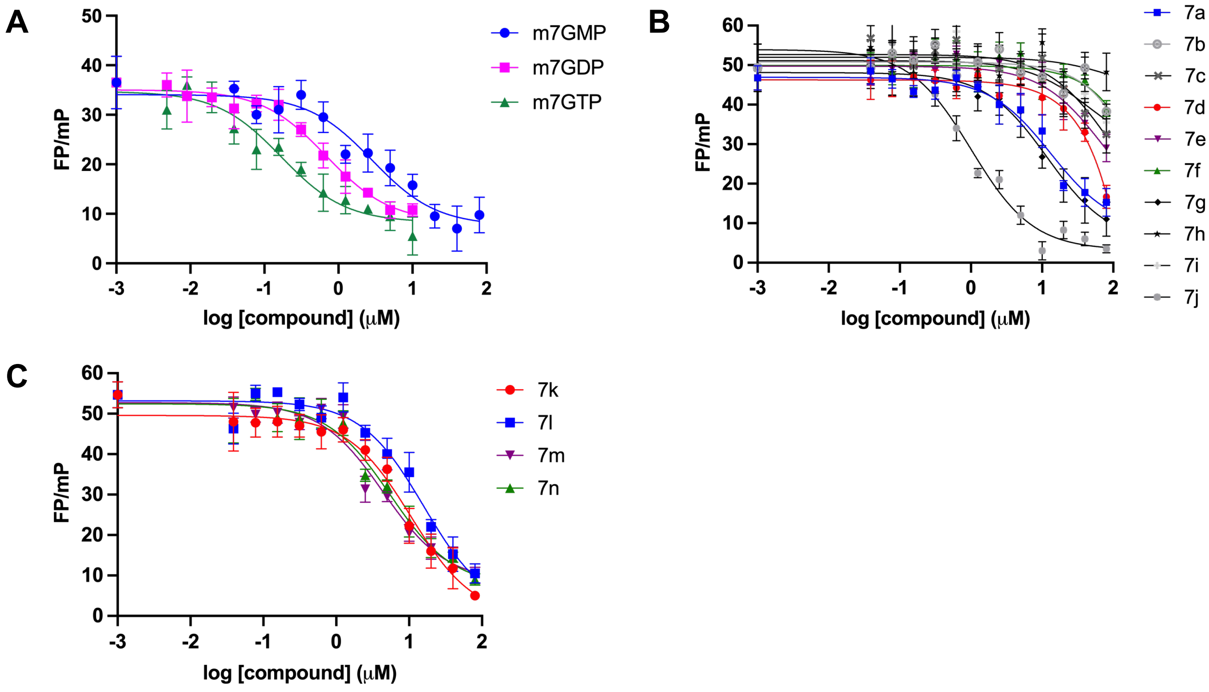


**Figure S1.** Inhibition curves for (A) m$^7$GMP, m$^7$GDP, and m$^7$GTP, and (B) and (C) unprotected cap analogues **7a–7n** measured using a fluorescence polarization assay.

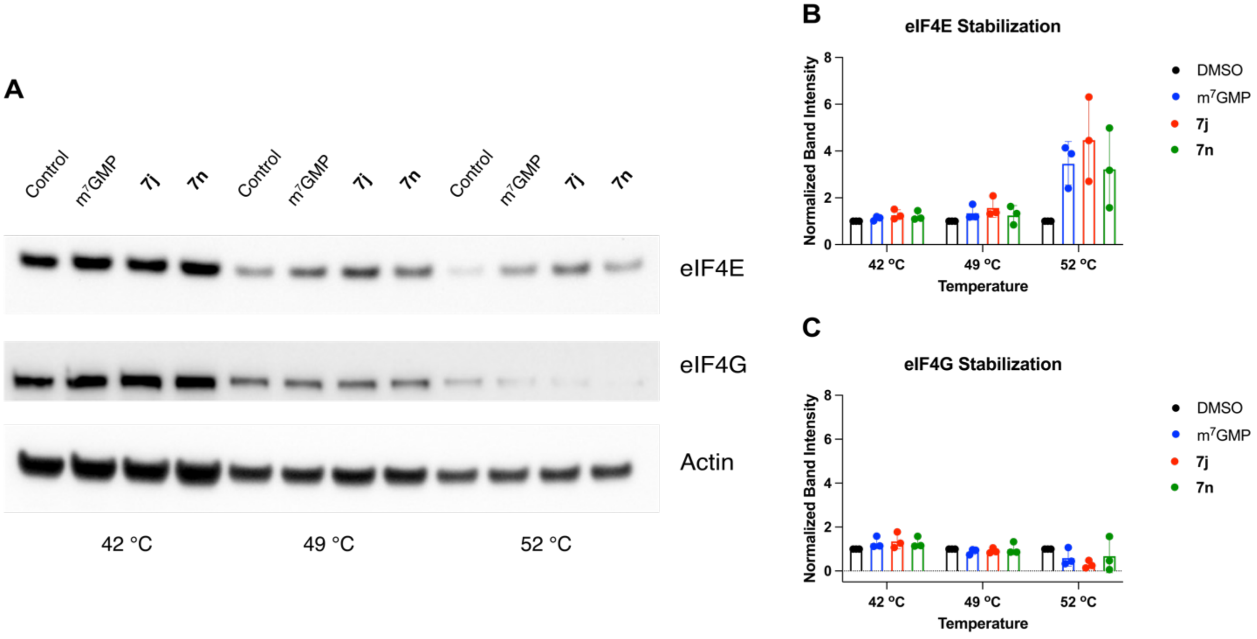


**Figure S2.** CETSA. (A) eIF4E target engagement measured using CETSA in HeLa cell lysate. eIF4E was detected via Western blot. (B) Quantitation in comparison to DMSO as a control.

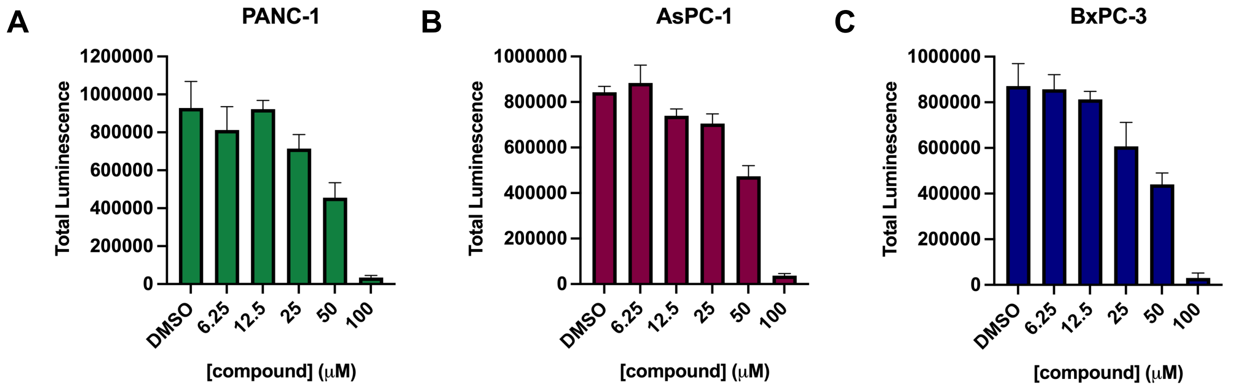


D

| Cell Line | $EC_{50}$ (μM) |
|---|---|
| PANC-1 | 69.5 ± 1.6 |
| AsPC-1 | 59.0 ± 1.5 |
| BxPC-3 | 148 ± 3 |

**Figure S3.** Anti-proliferative activity of **6n** in additional pancreatic cancer cell lines (A) PANC-1, (B) AsPC-1, and (C) BxPC-3. (D) Table of $EC_{50}$ values.

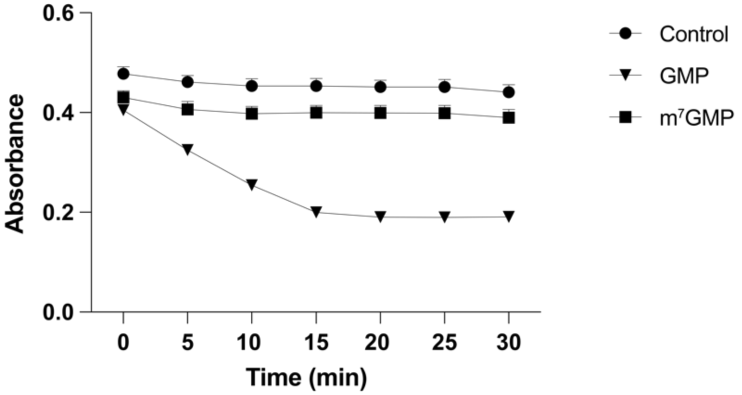


**Figure S4.** Alkylation of the $N^7$ position abrogates GMPK activity. GMPK assays were carried out as described by Khan *et al.* (*J. Biol. Chem.* **2019**, *294*, 11920−11933) using 1 nM GMPK and 100 μM GMP or $m^7$GMP.

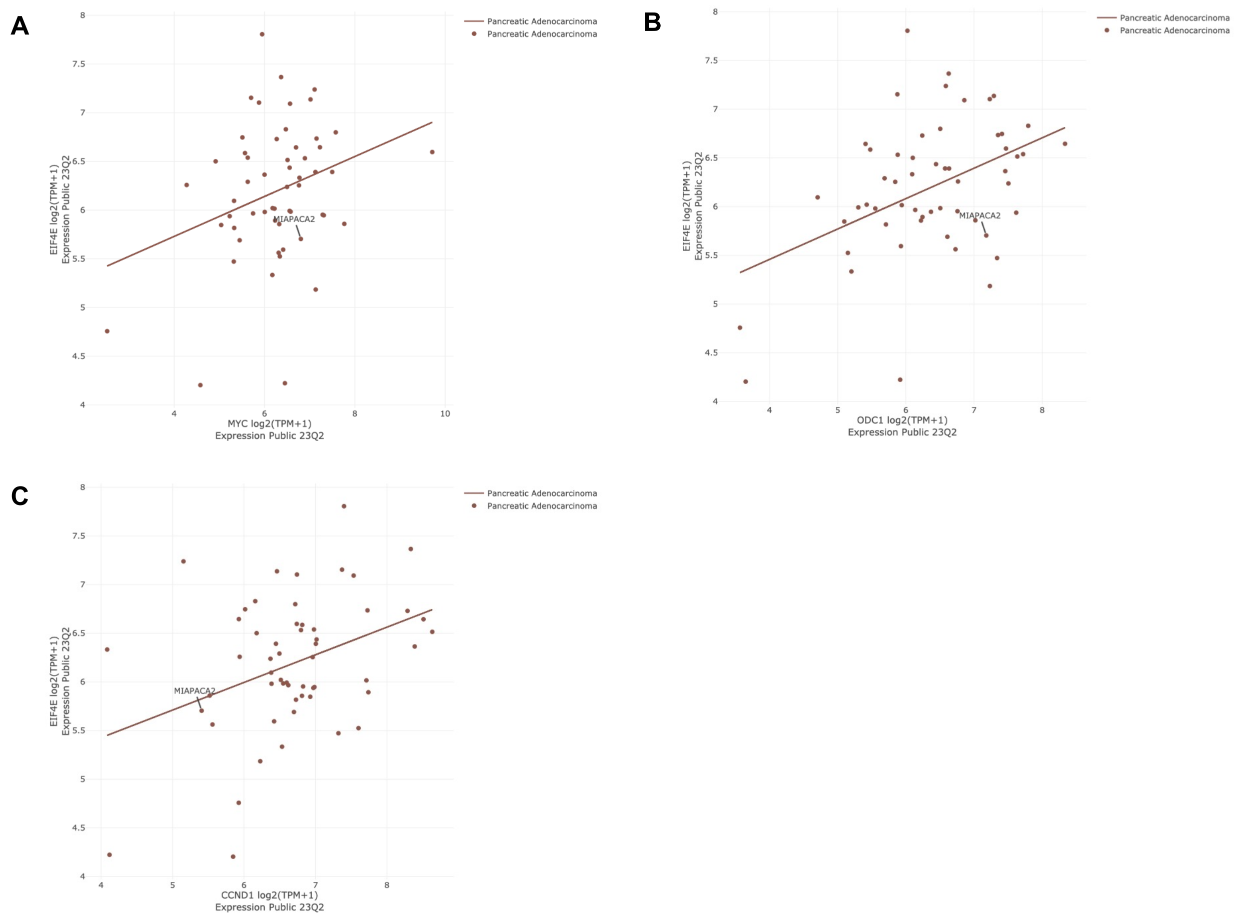


**Figure S5.** Correlation of eIF4E expression (*y*-axis) with the expression of select cap-dependent transcripts (*x*-axis) in pancreatic cancer cells generated using the DepMap Portal. MiaPaca-2 cells used in this study are indicated in each plot. (A) c-Myc. (B) ODC1. (C) Cyclin D1.

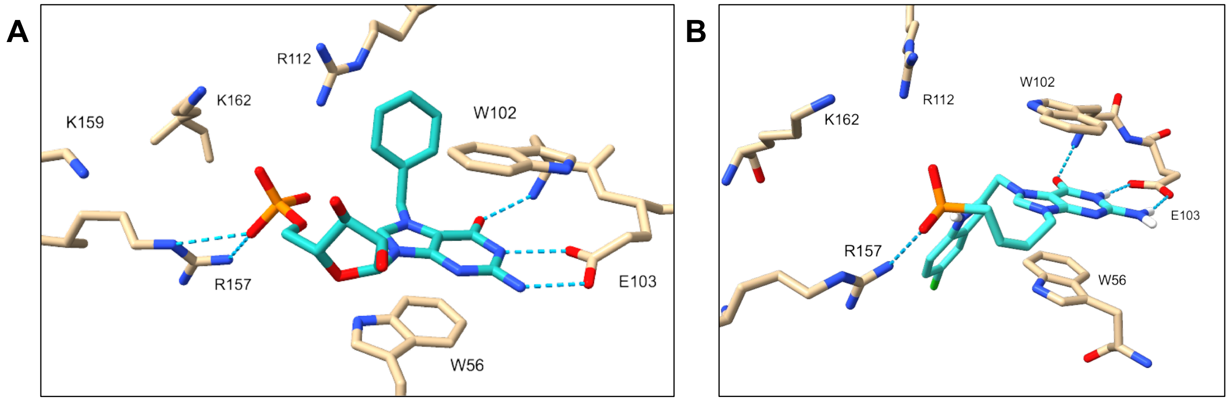


**Figure S6.** Comparison of the binding modes of (A) $Bn^7GMP$ (PDB: 2V8X) and (B) **7n** (PDB: 8SX4) within eIF4E's cap-binding pocket.

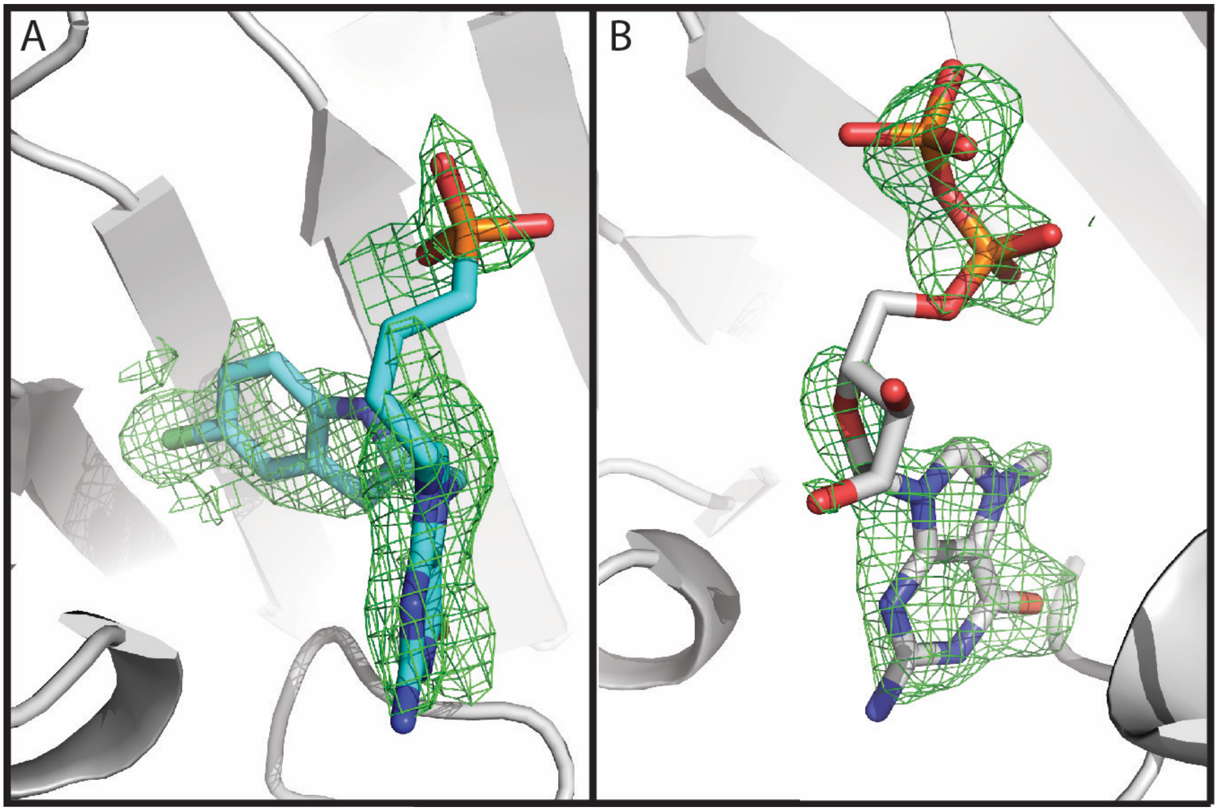


**Figure S7.** Difference electron density map produced from the molecular replacement results. The difference map (Fo-Fc) is shown as a green grid contoured at 2.5s. (A) Density representing **7n** was noted in chain A, while (B) density for $m^7GDP$ was apparent in chain B. (PDB: 8SX4)

### B. $^{1}H$ and $^{13}C$ NMR Spectra

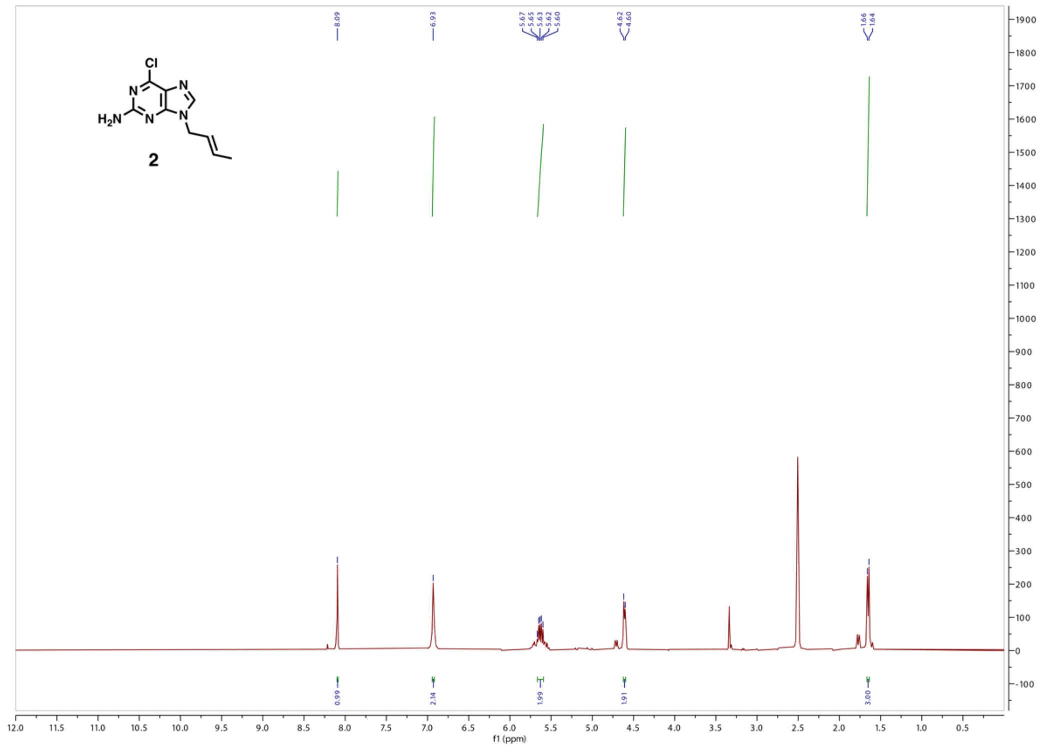


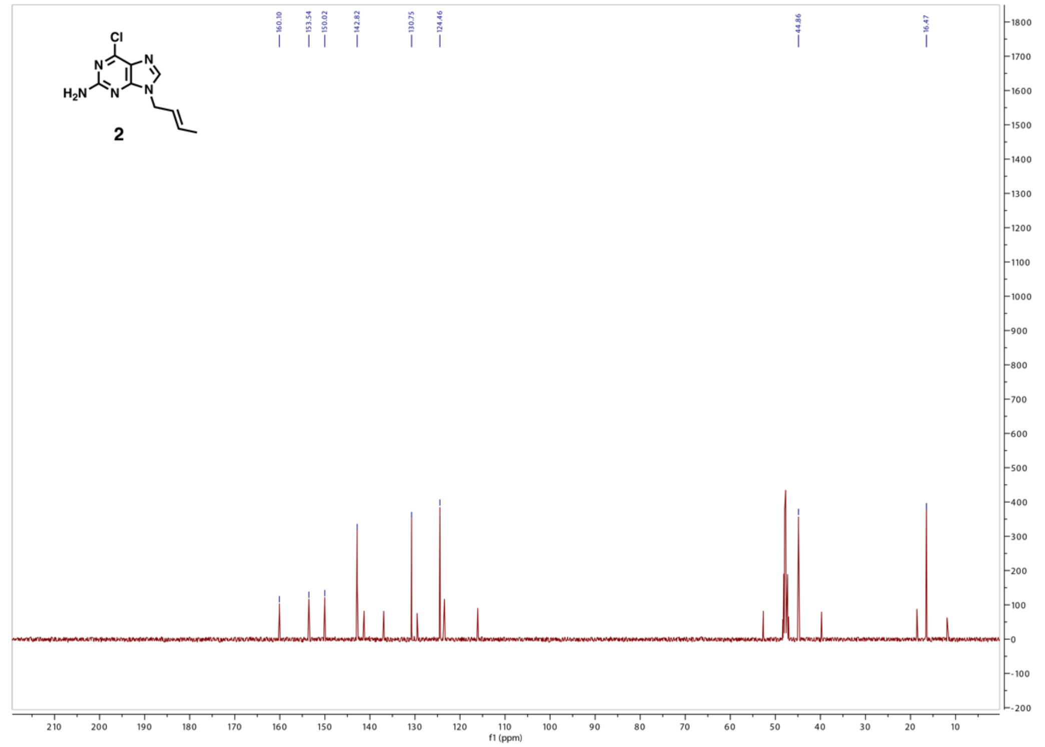

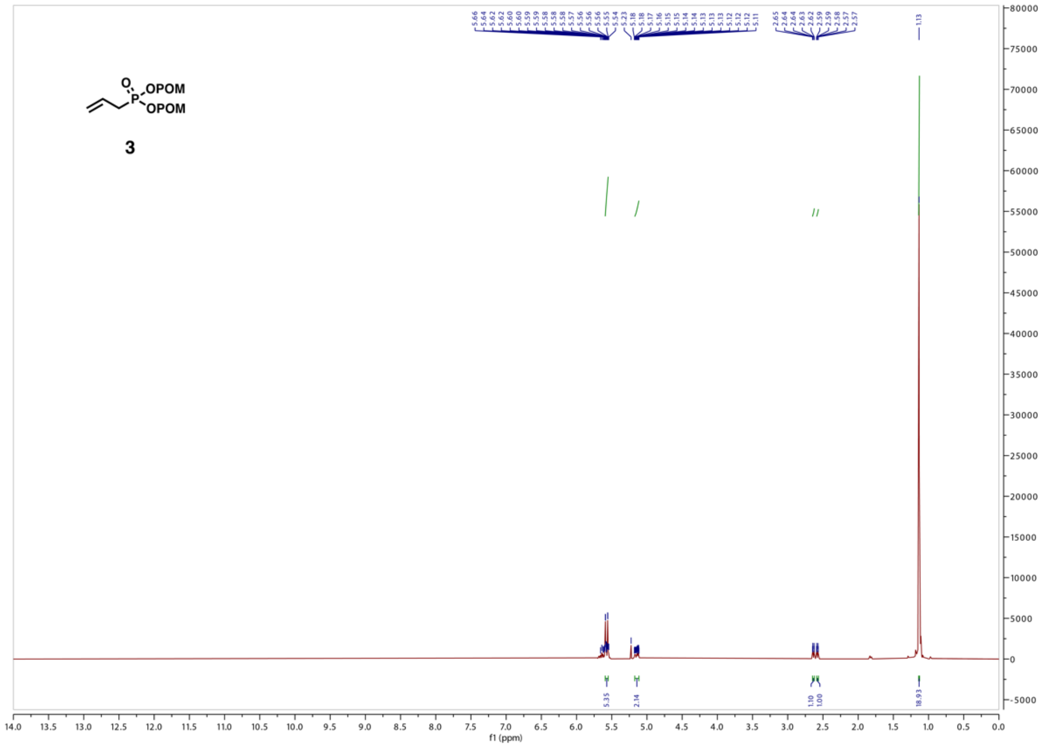

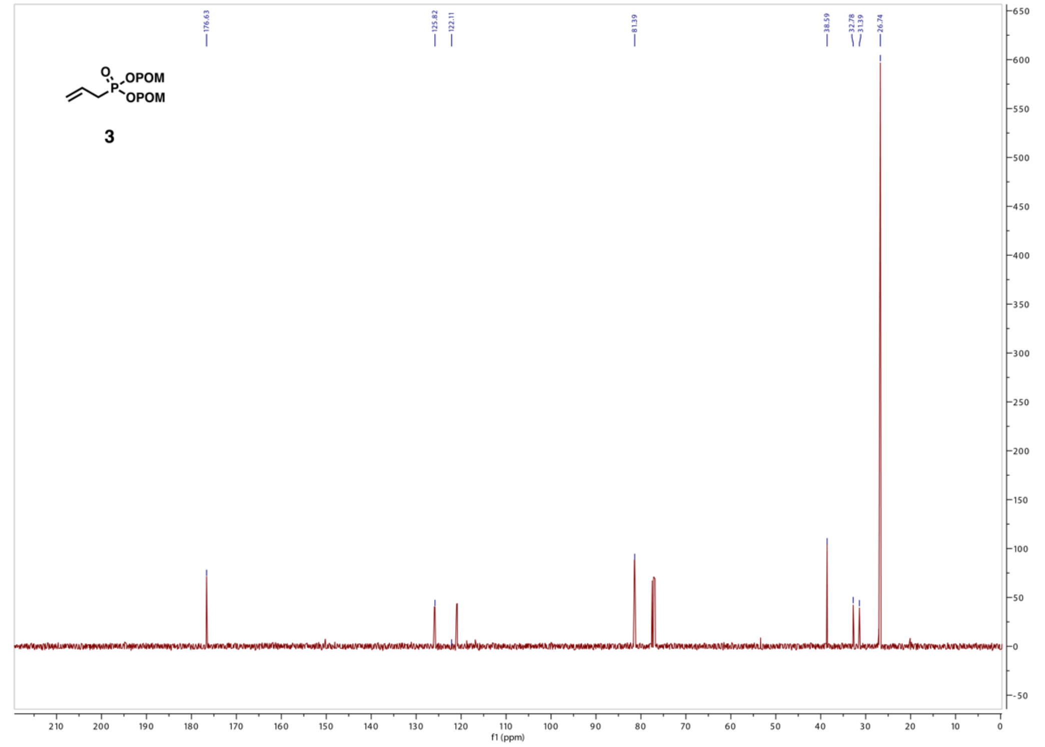

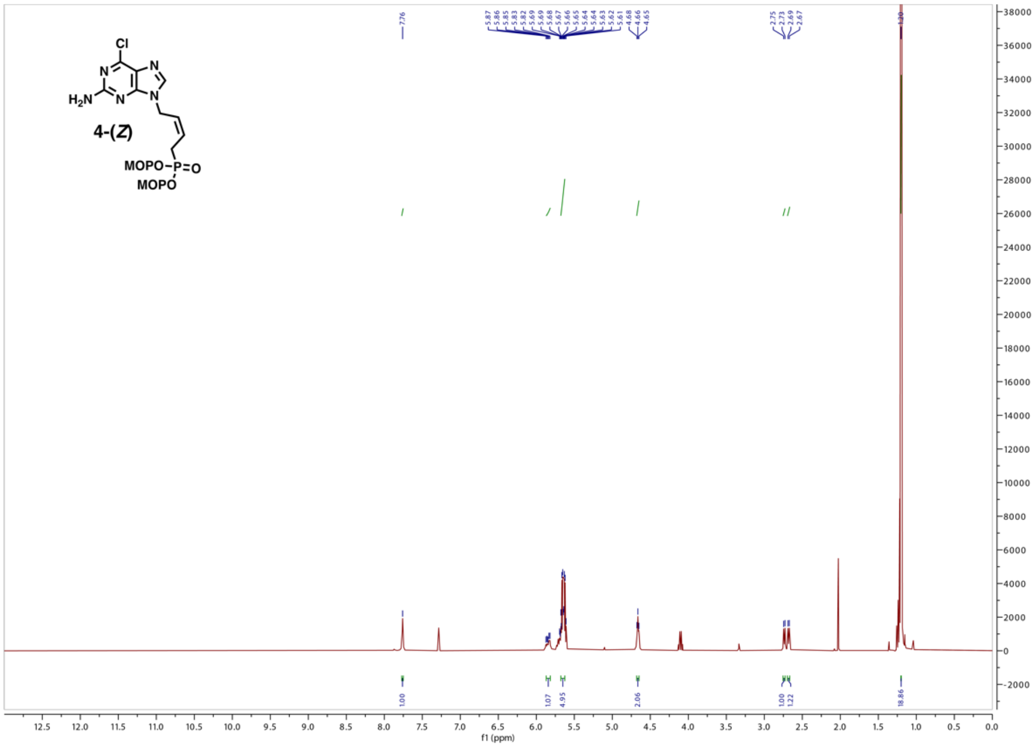
Cl
N
N
H2N
N
N
4-(Z)
MOPO–P=O
MOPO
f1 (ppm)

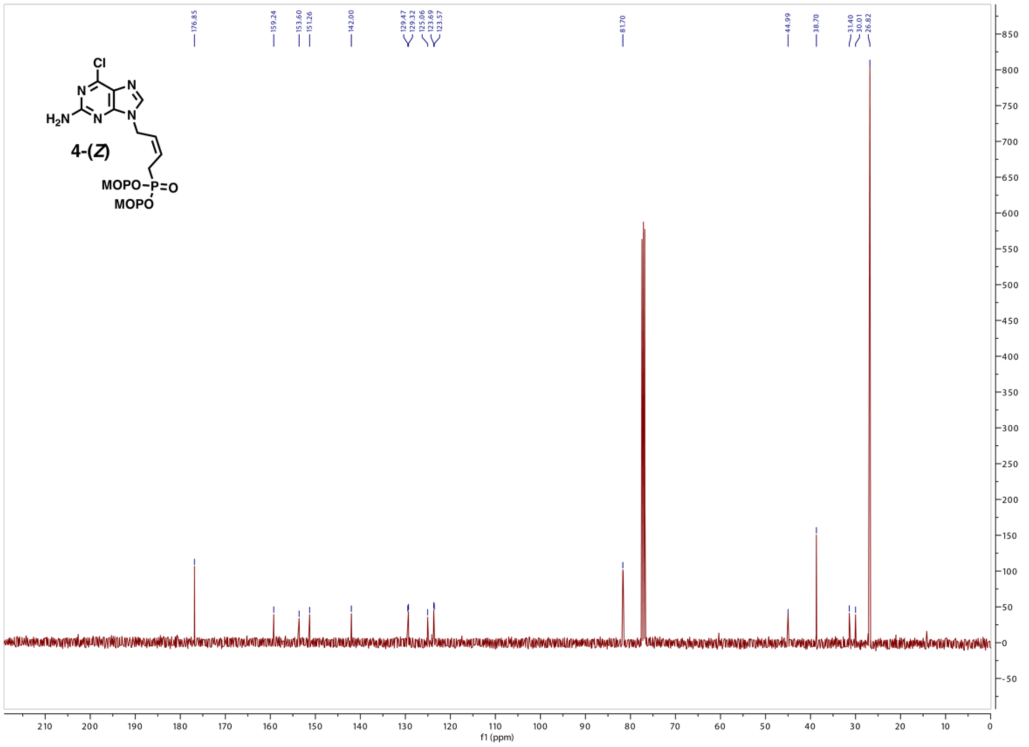
Cl
N
N
H2N
N
N
4-(Z)
MOPO–P=O
MOPO
f1 (ppm)

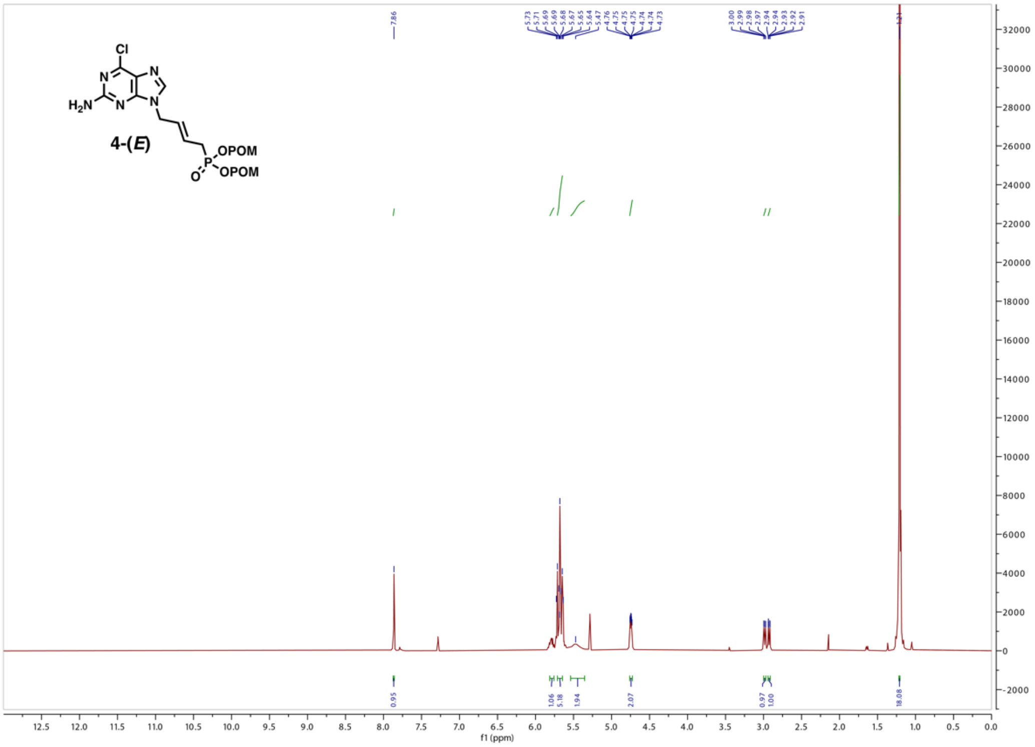
4-(E)
f1 (ppm)

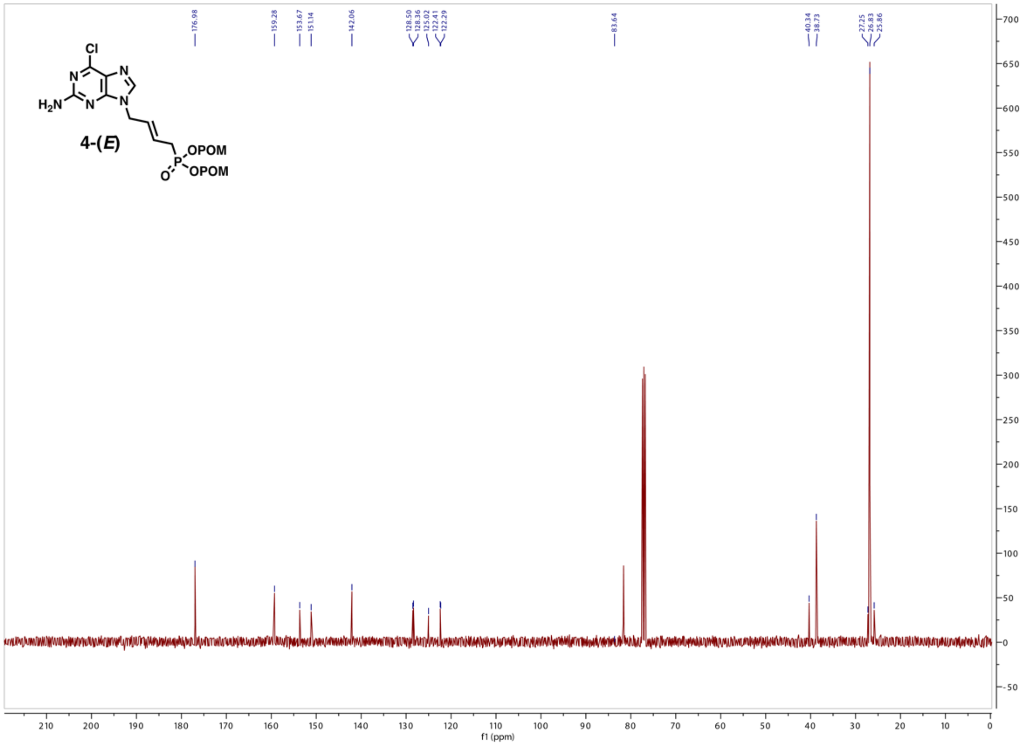
4-(E)
f1 (ppm)

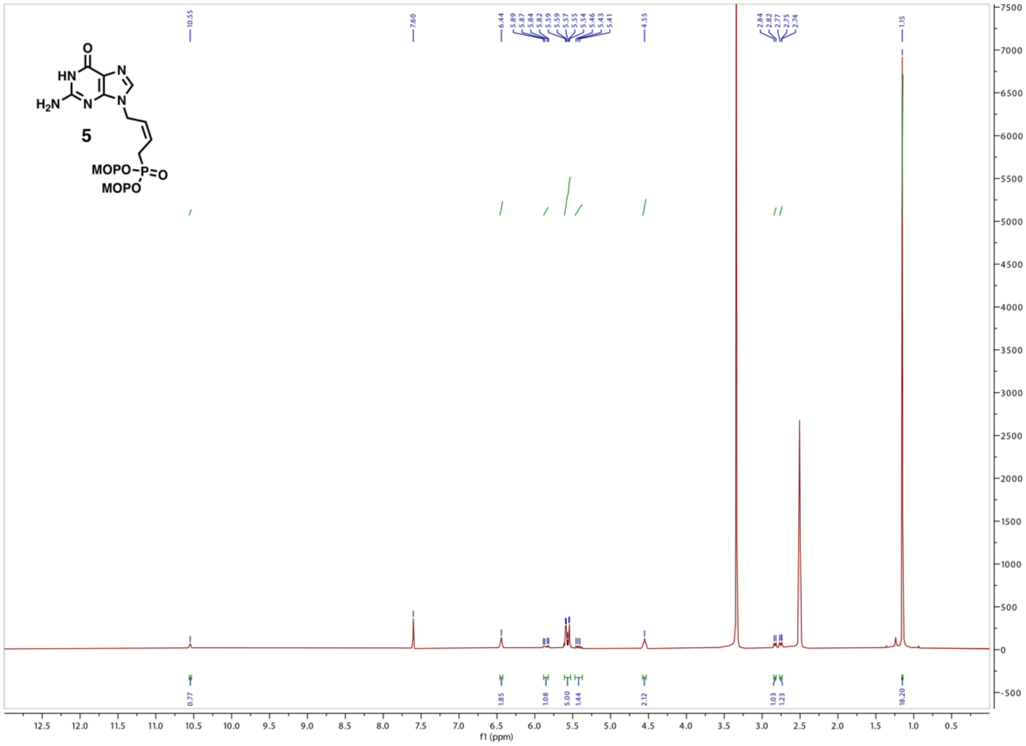

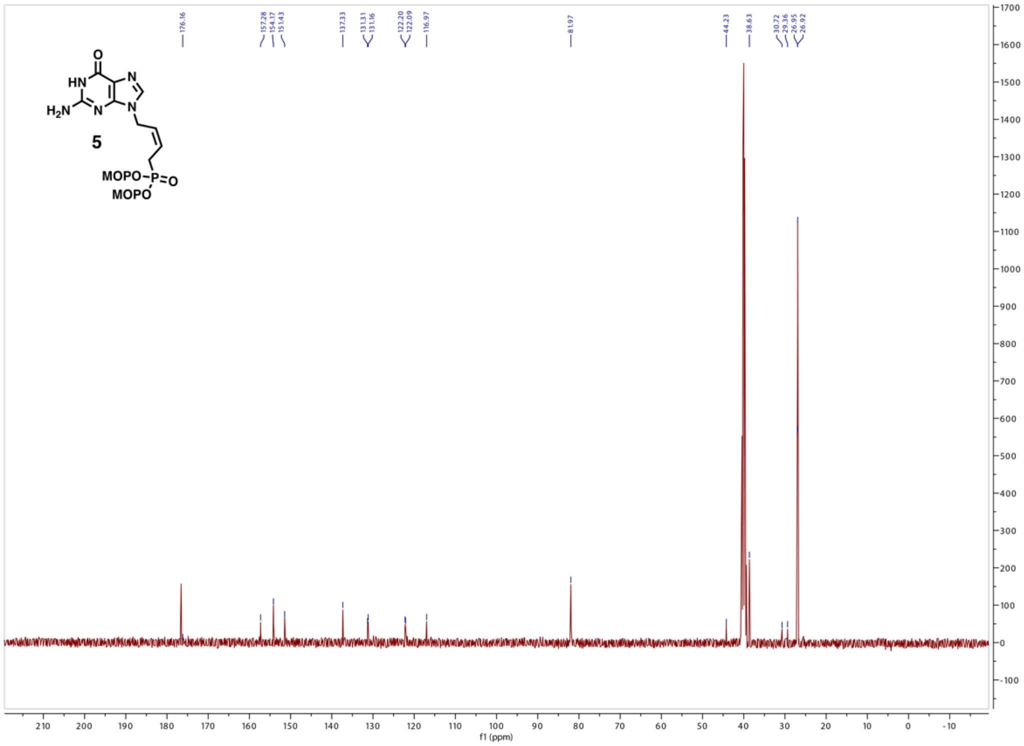

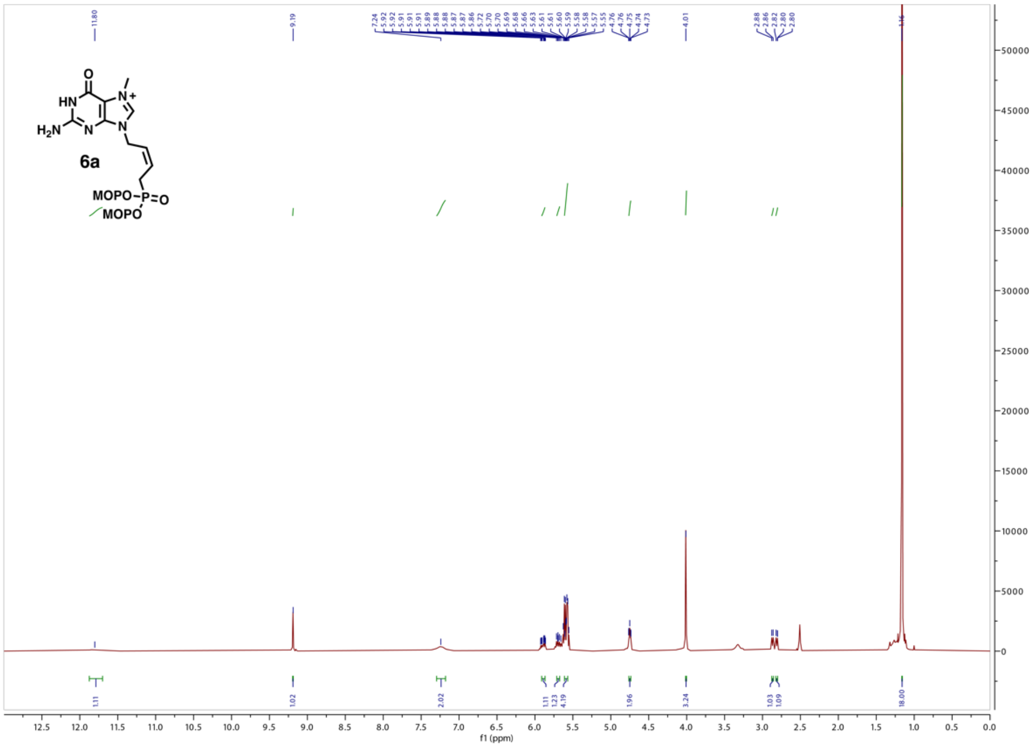
6a
MOPO
MOPO
f1 (ppm)

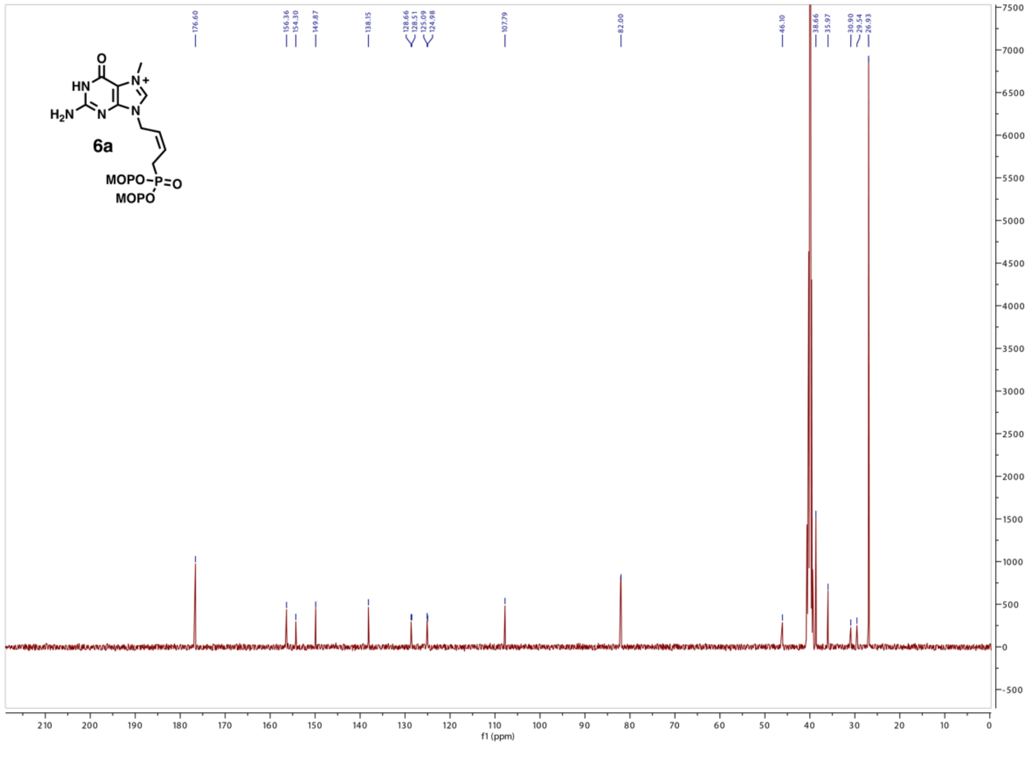
6a
MOPO
MOPO
f1 (ppm)

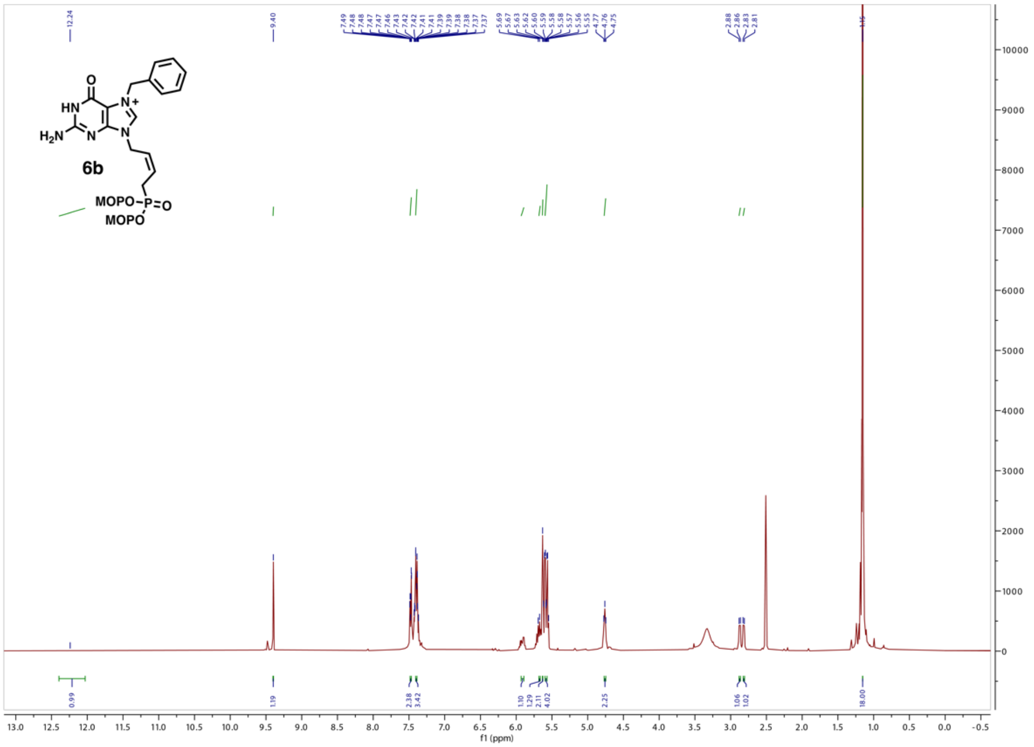

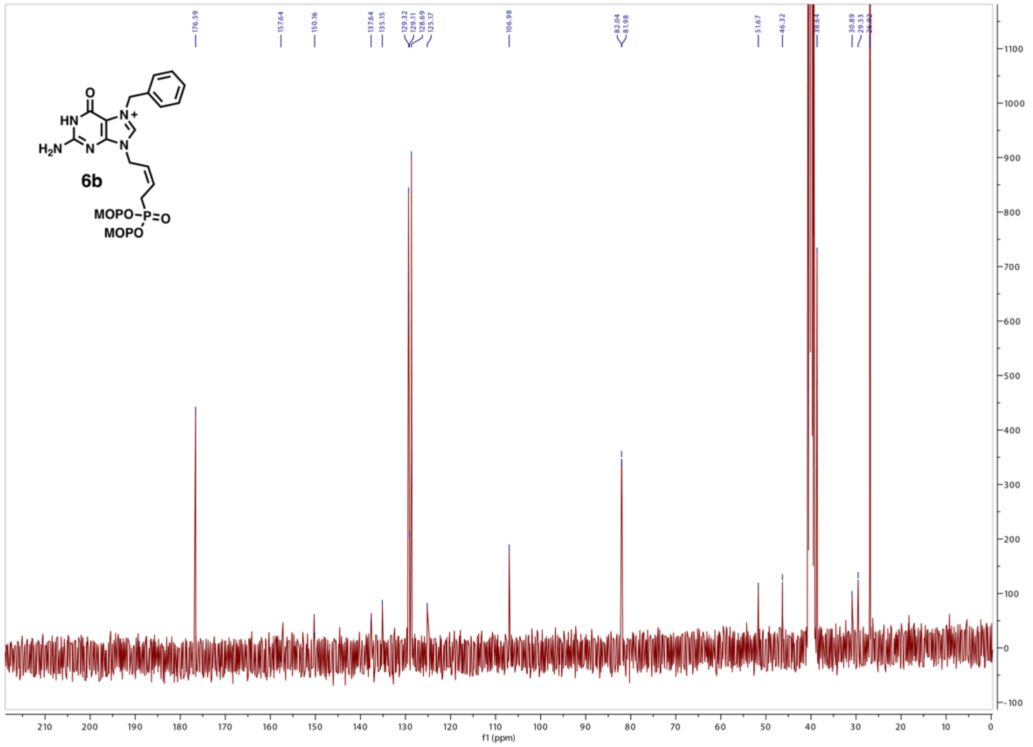

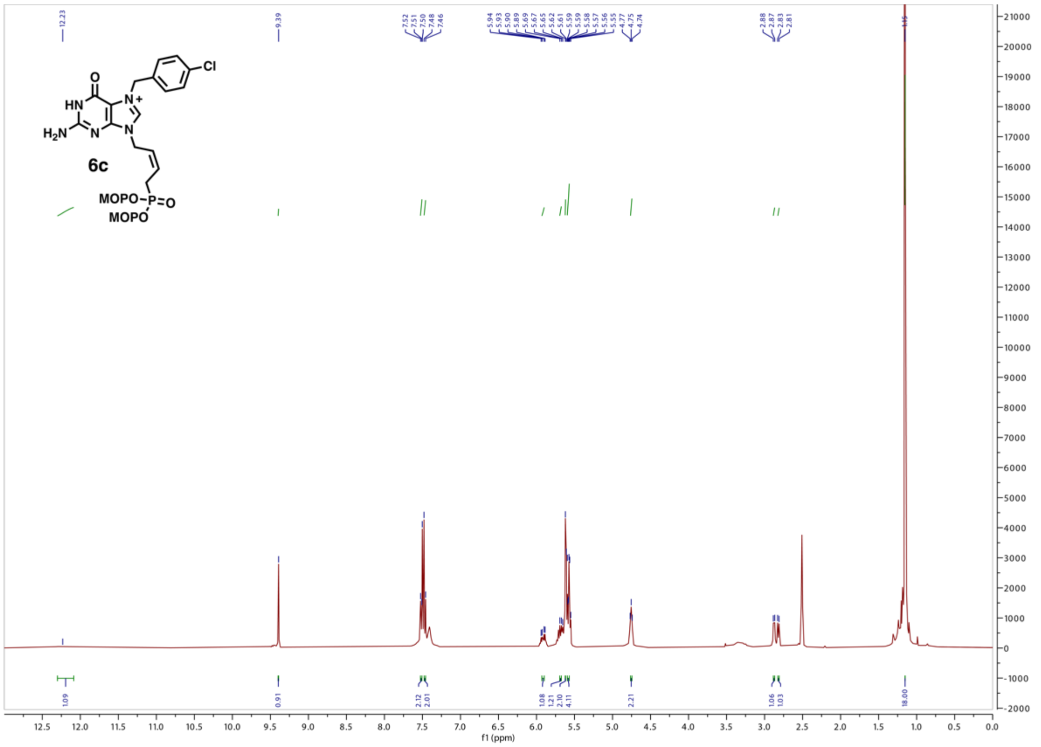

6c
MOPO
MOPO
f1 (ppm)


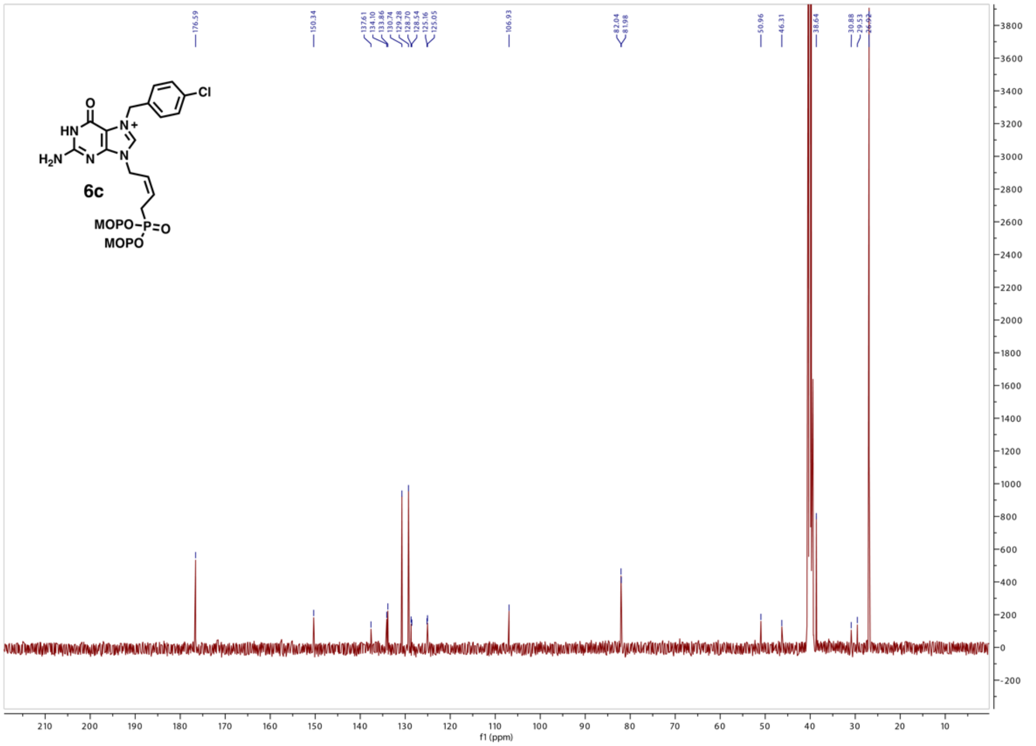

6c
MOPO
MOPO
f1 (ppm)

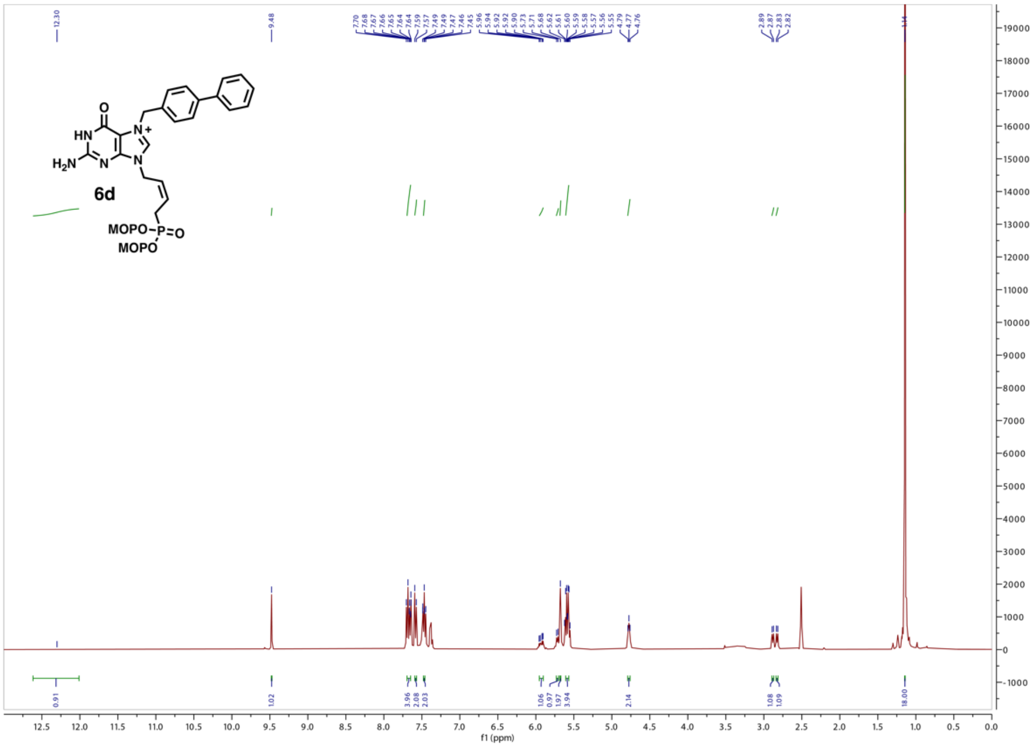

O
HN
N+
H2N
N
N
6d
MOPO-P=O
MOPO
f1 (ppm)


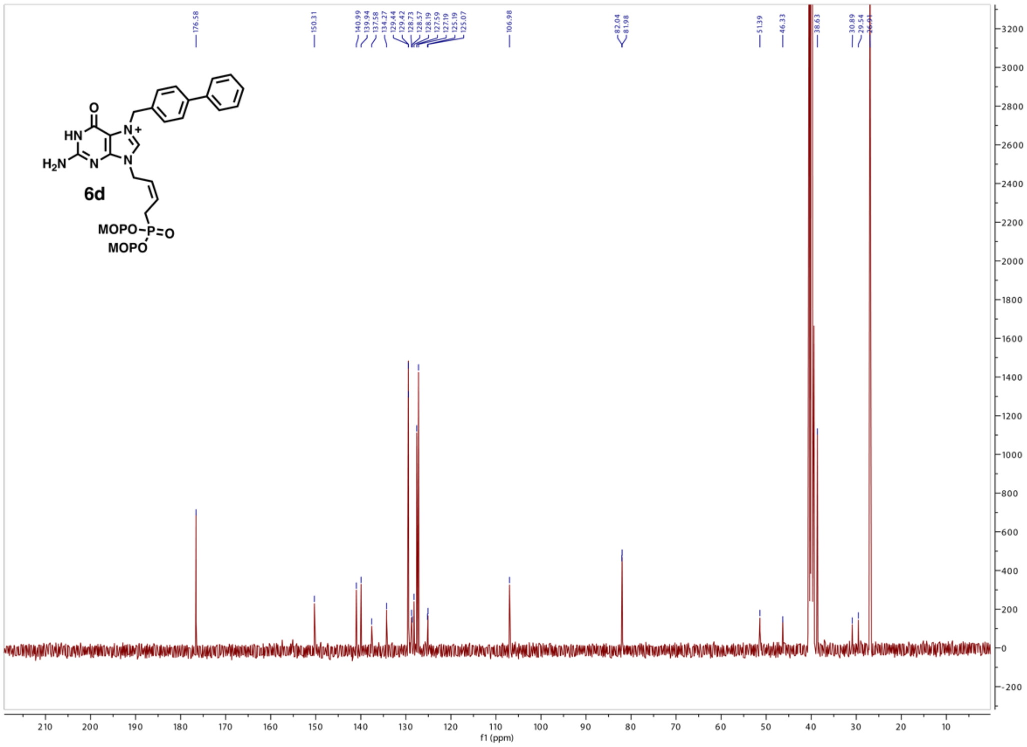

O
HN
N+
H2N
N
N
6d
MOPO-P=O
MOPO
f1 (ppm)

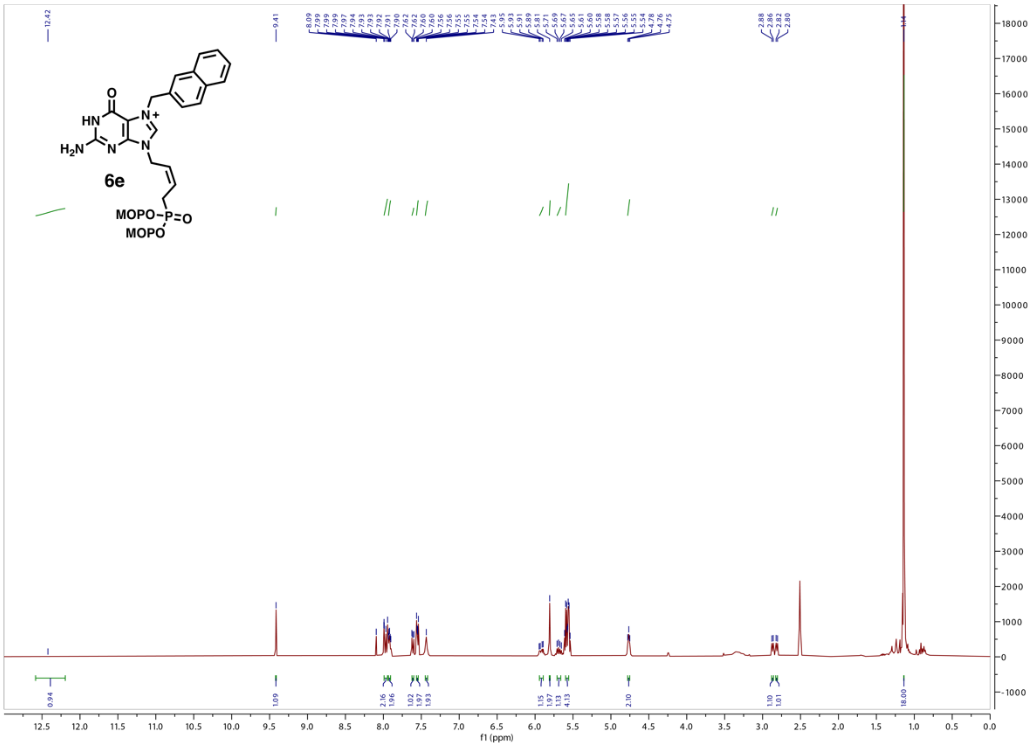

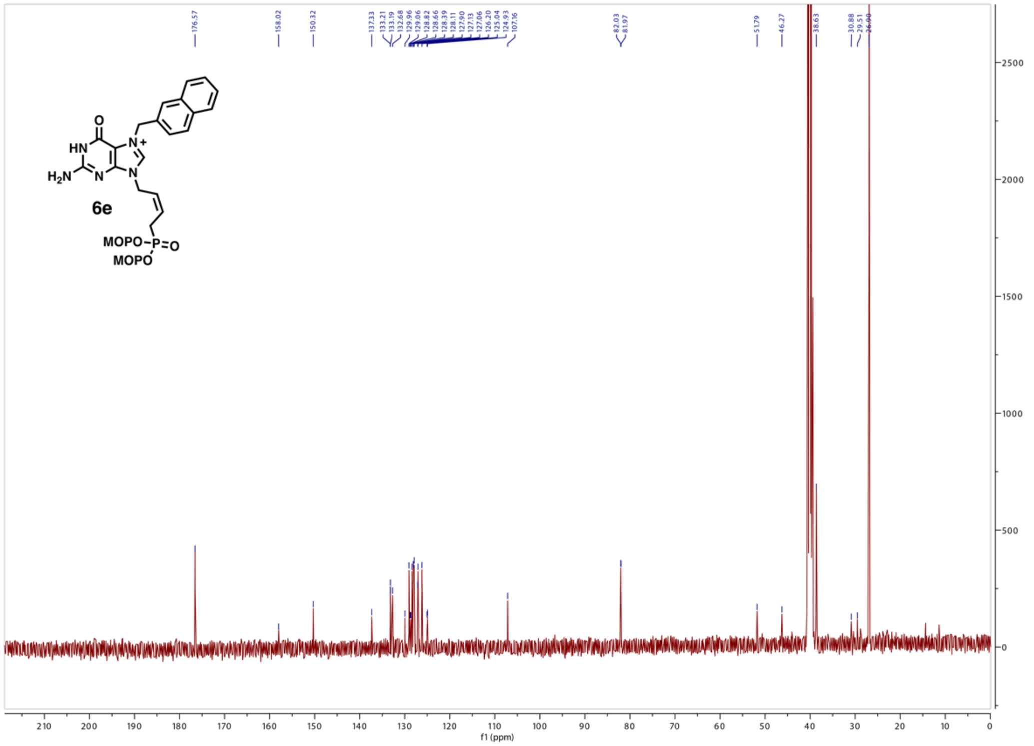

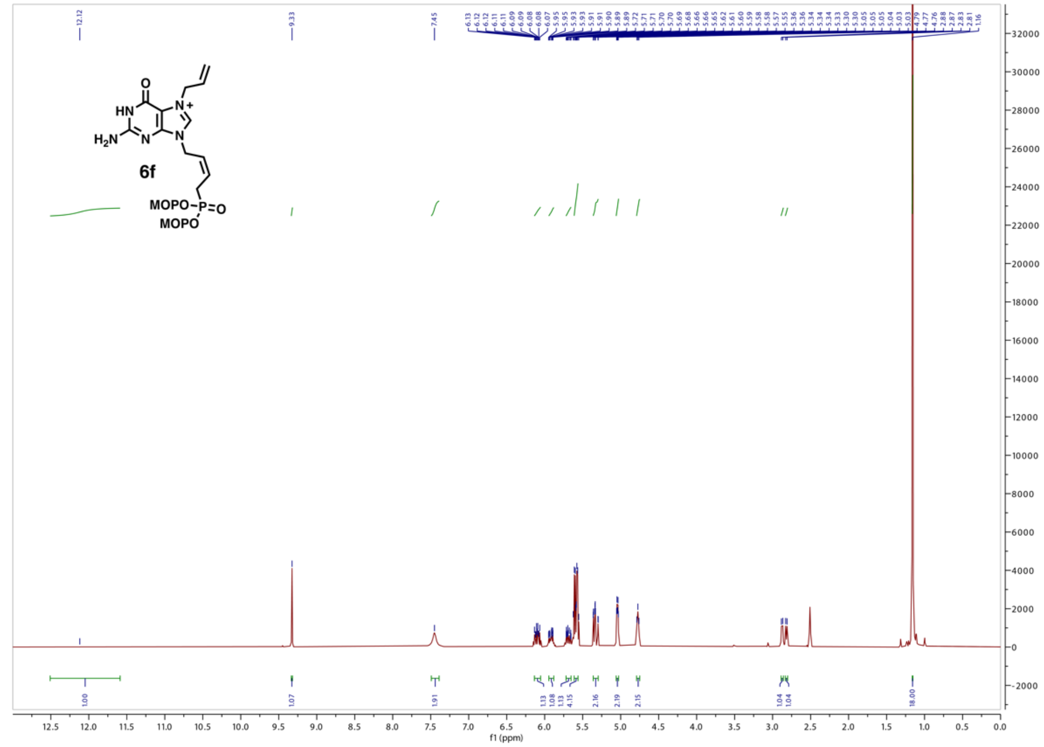

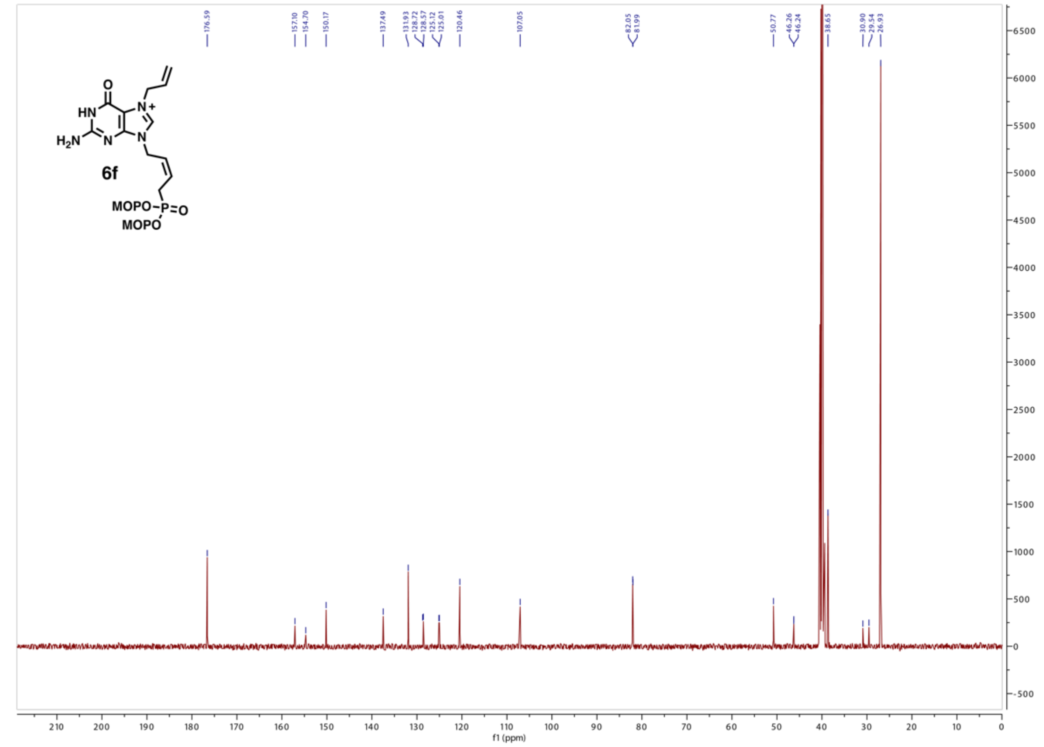

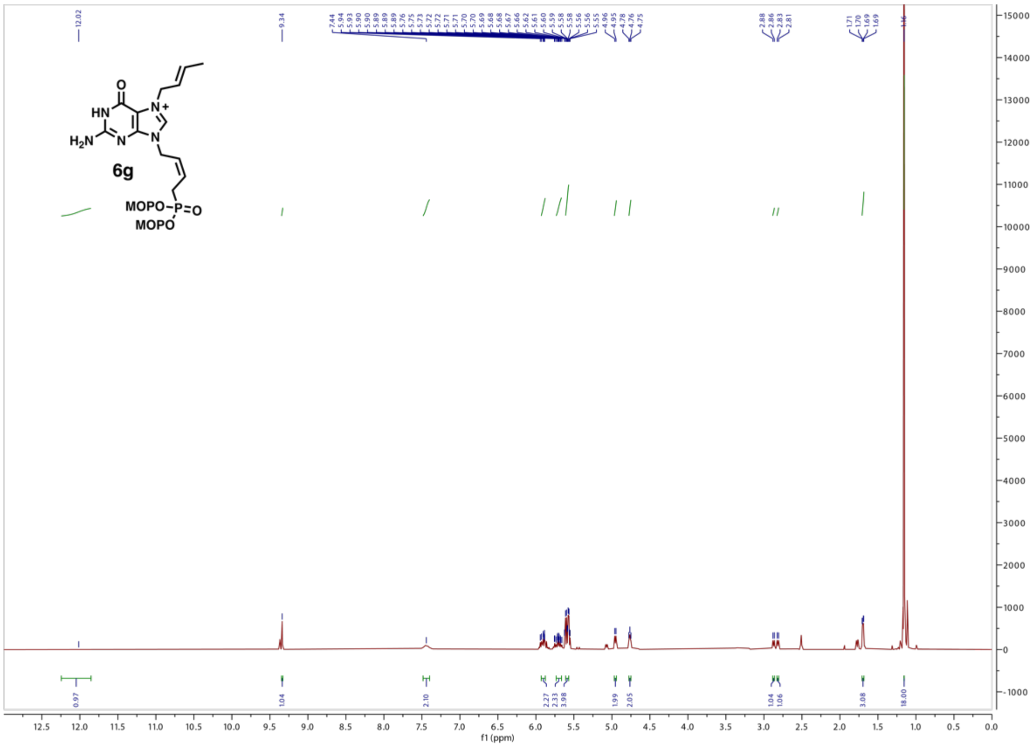
6g
MOPO
MOPO
f1 (ppm)

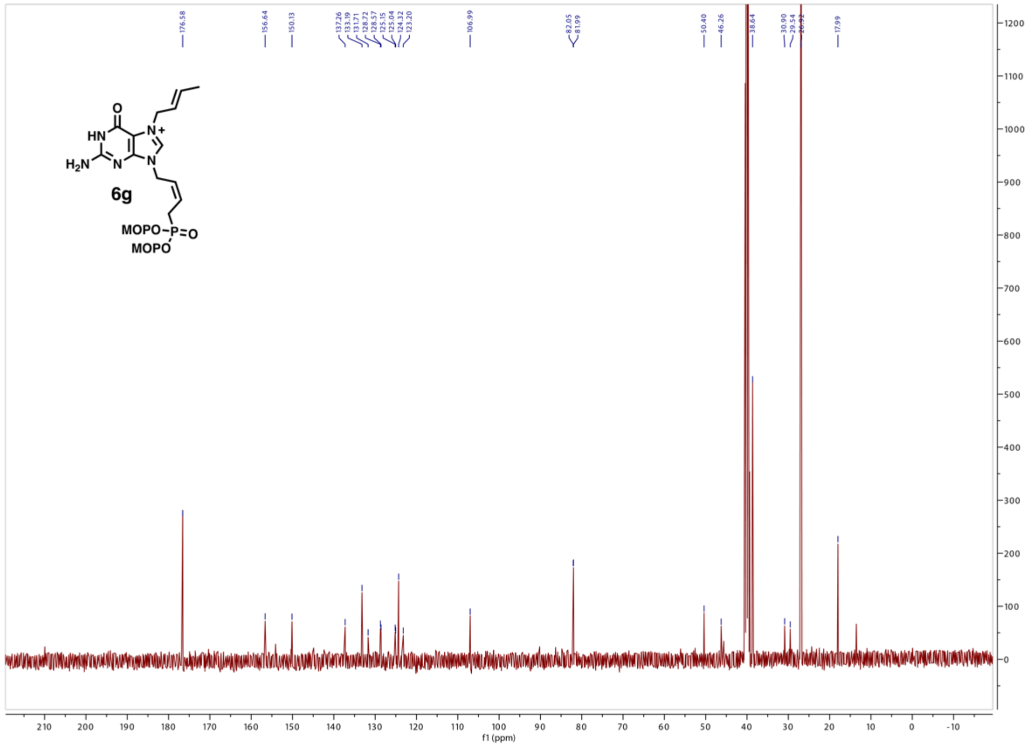
6g
MOPO
MOPO
f1 (ppm)

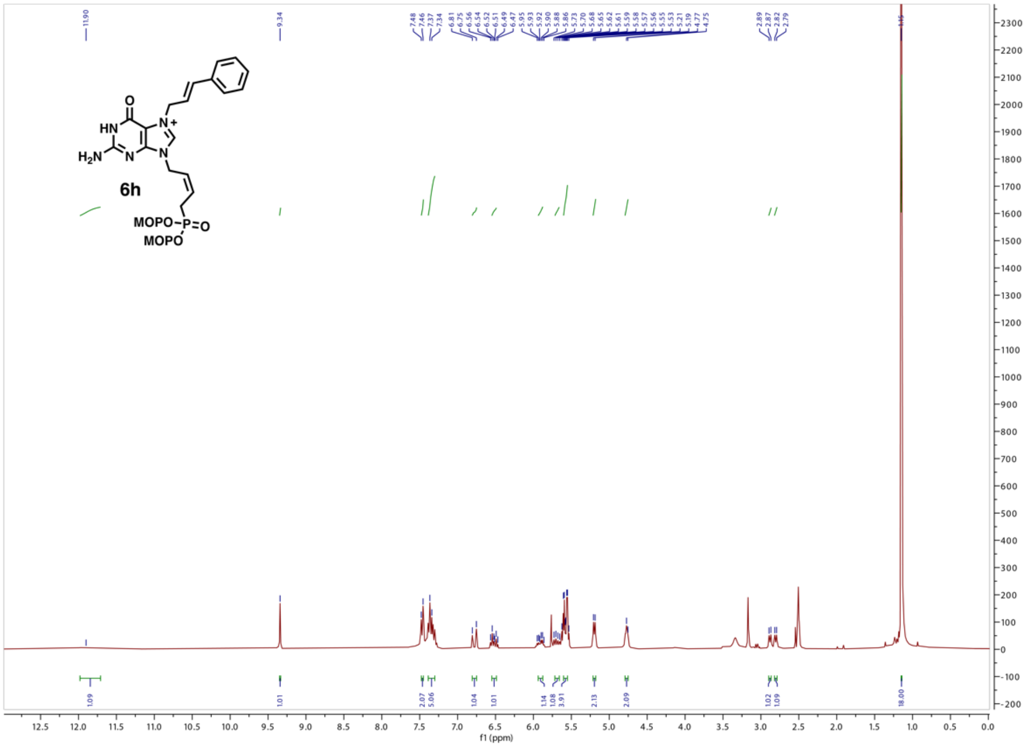

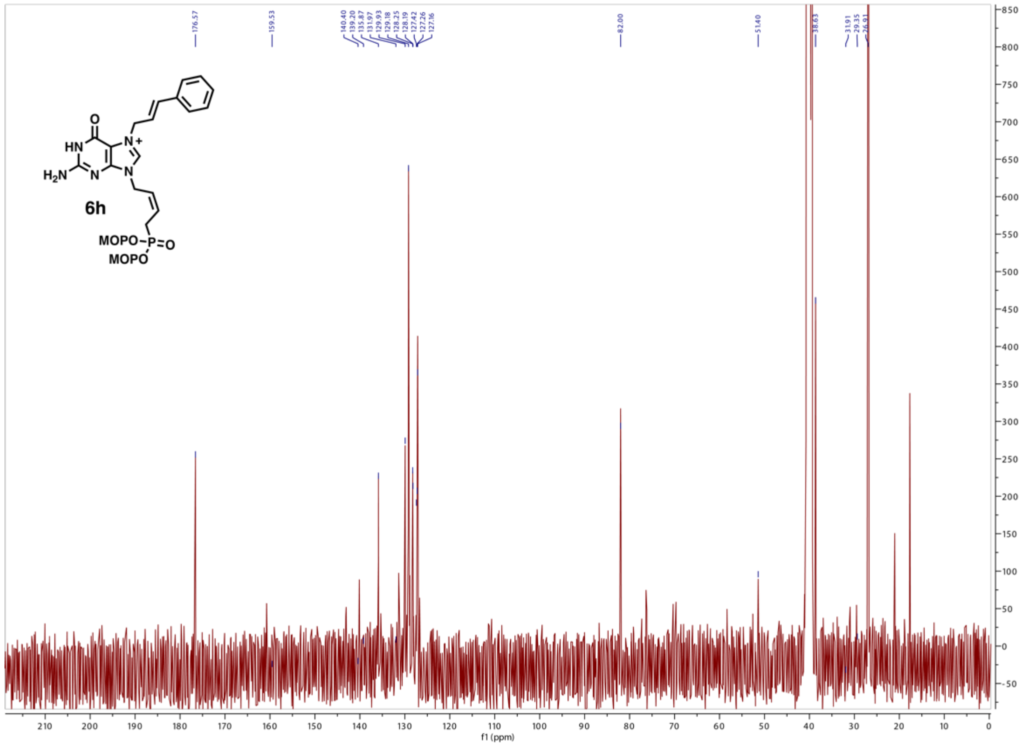

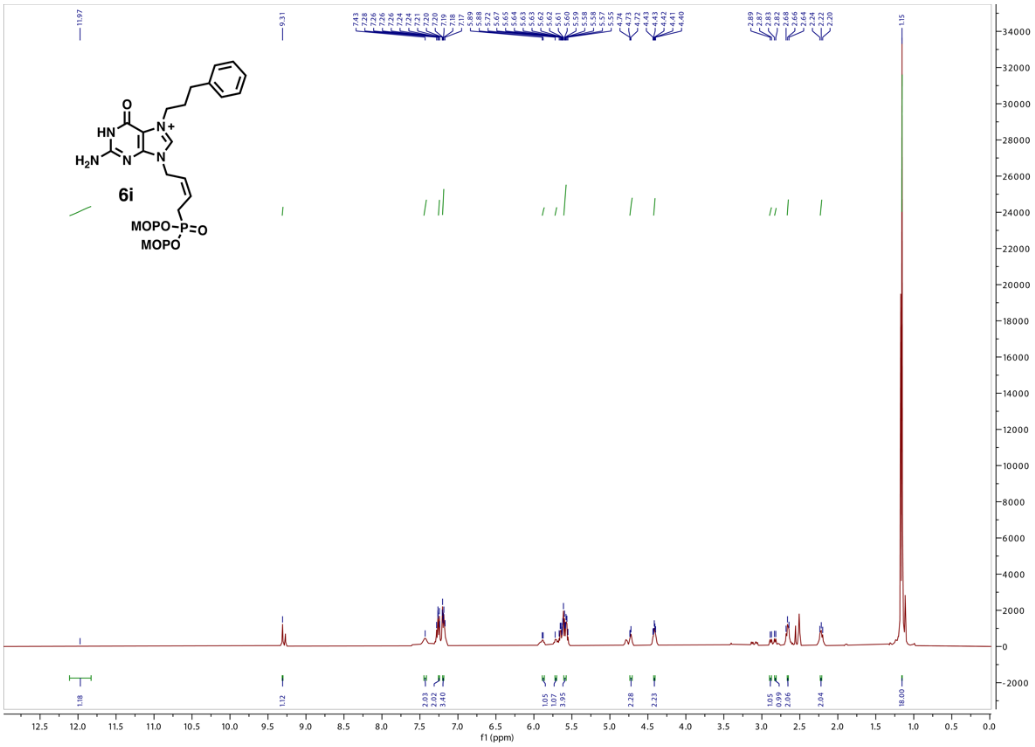

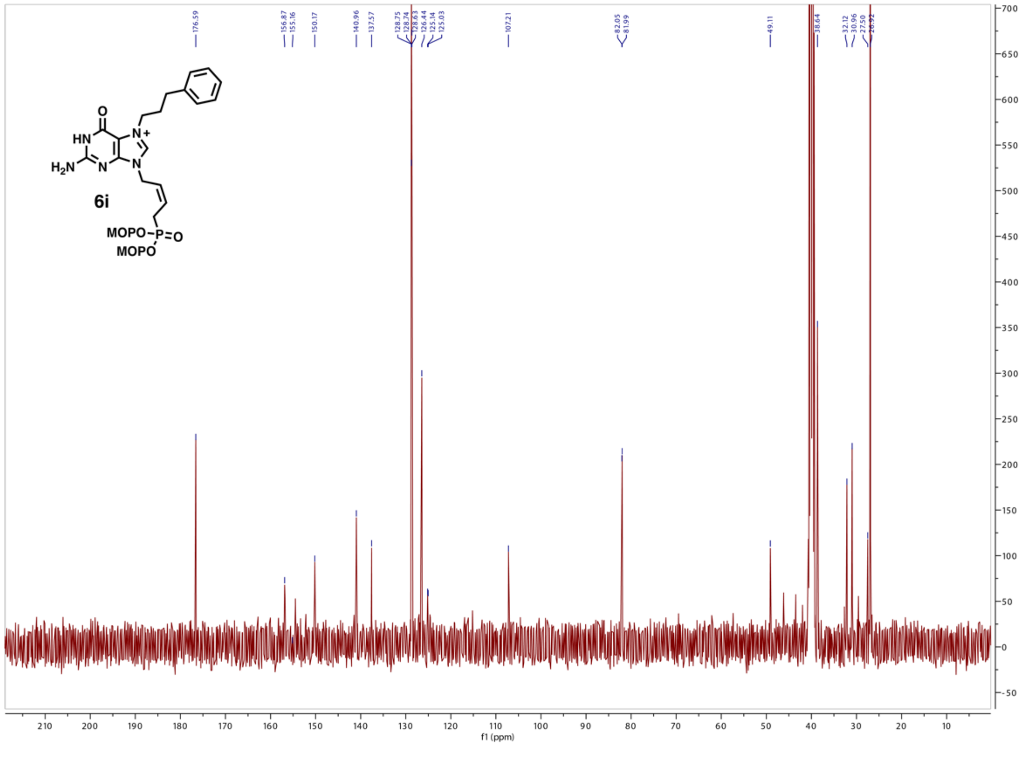

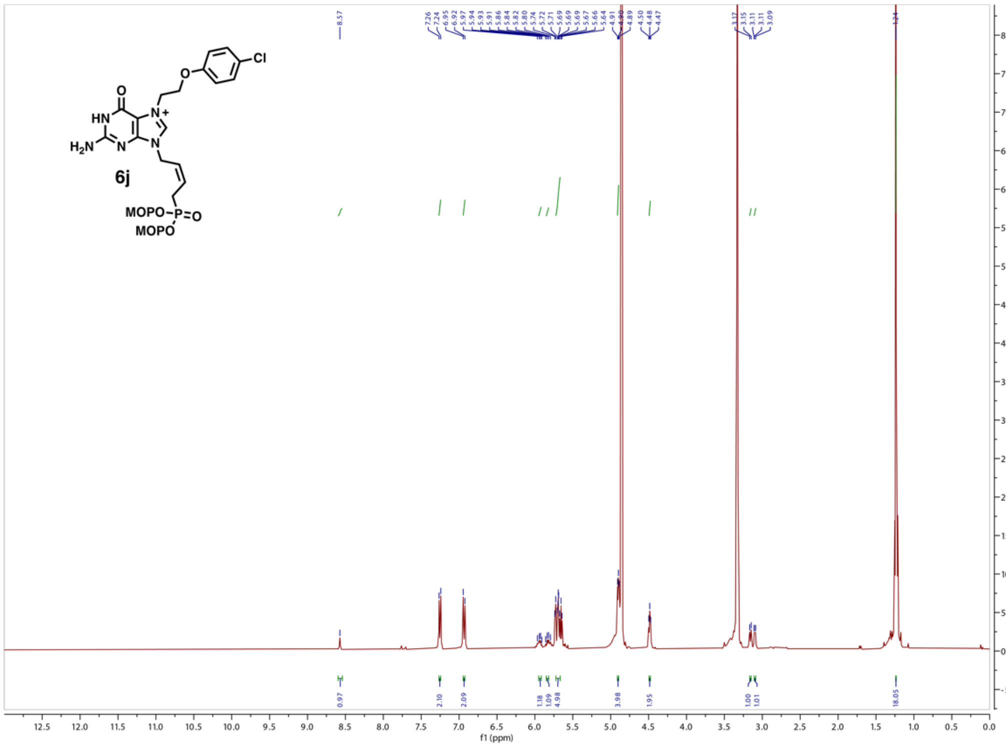

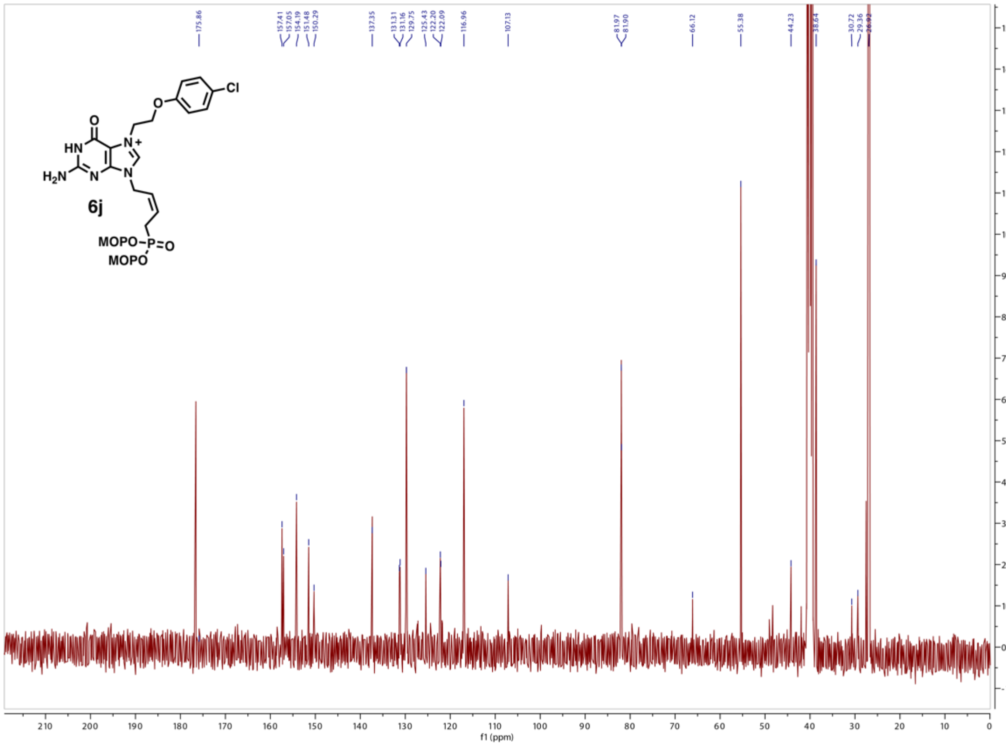

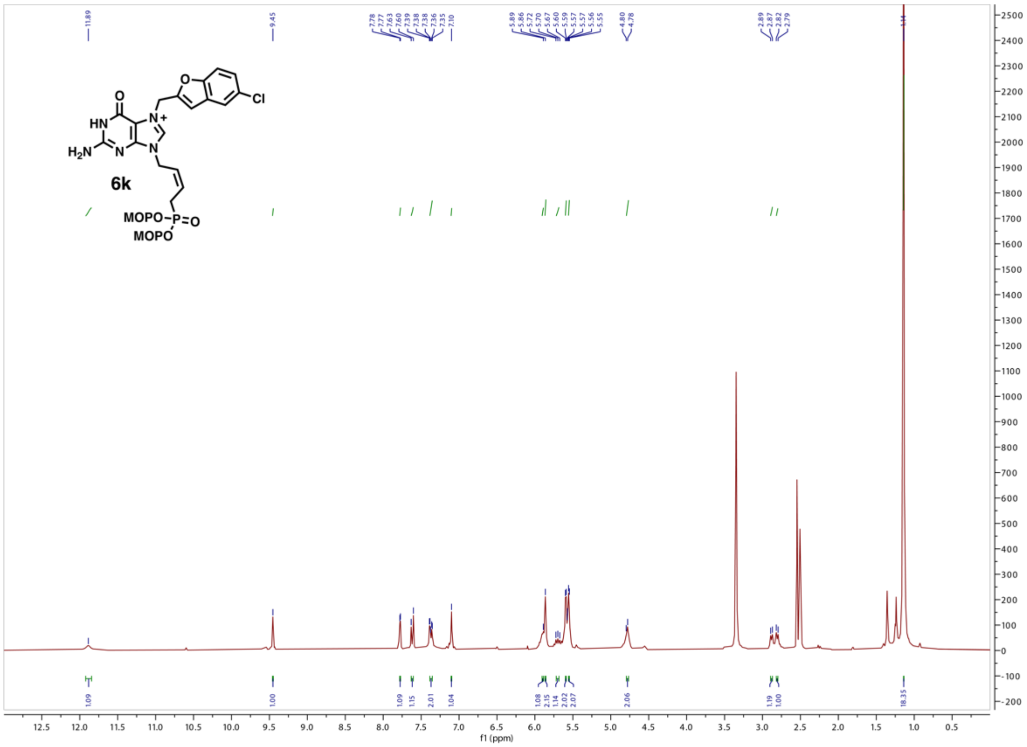
6k
MOPO
MOPO
f1 (ppm)

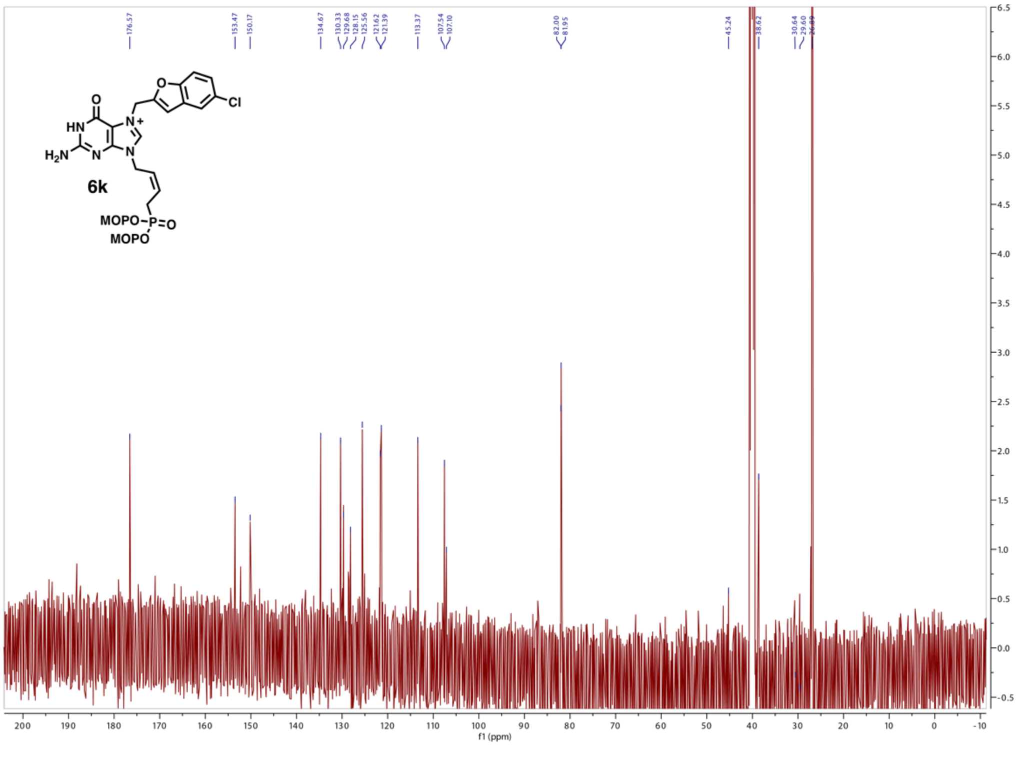
6k
MOPO
MOPO
f1 (ppm)

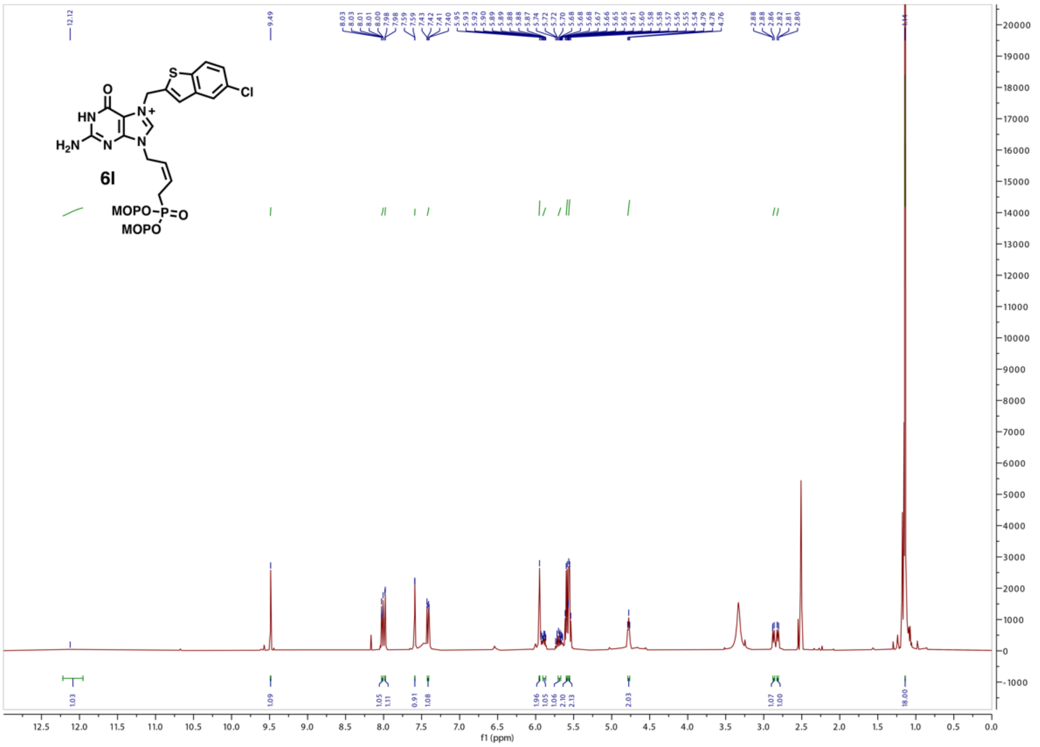
6l
MOPO
MOPO
f1 (ppm)

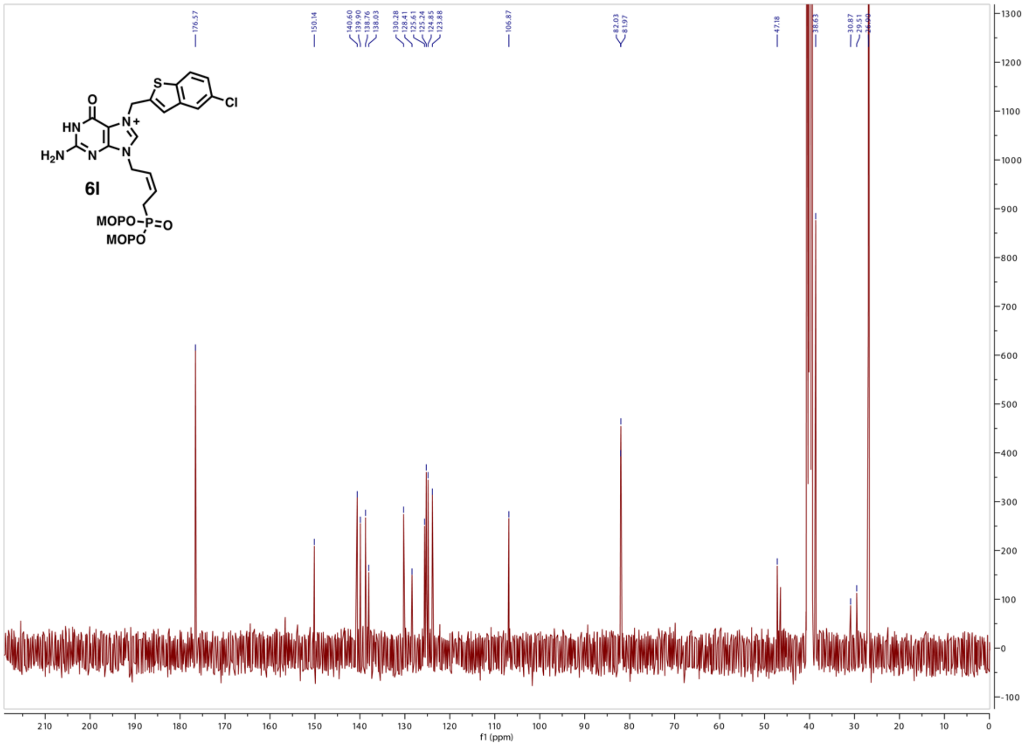
6l
MOPO
MOPO
f1 (ppm)

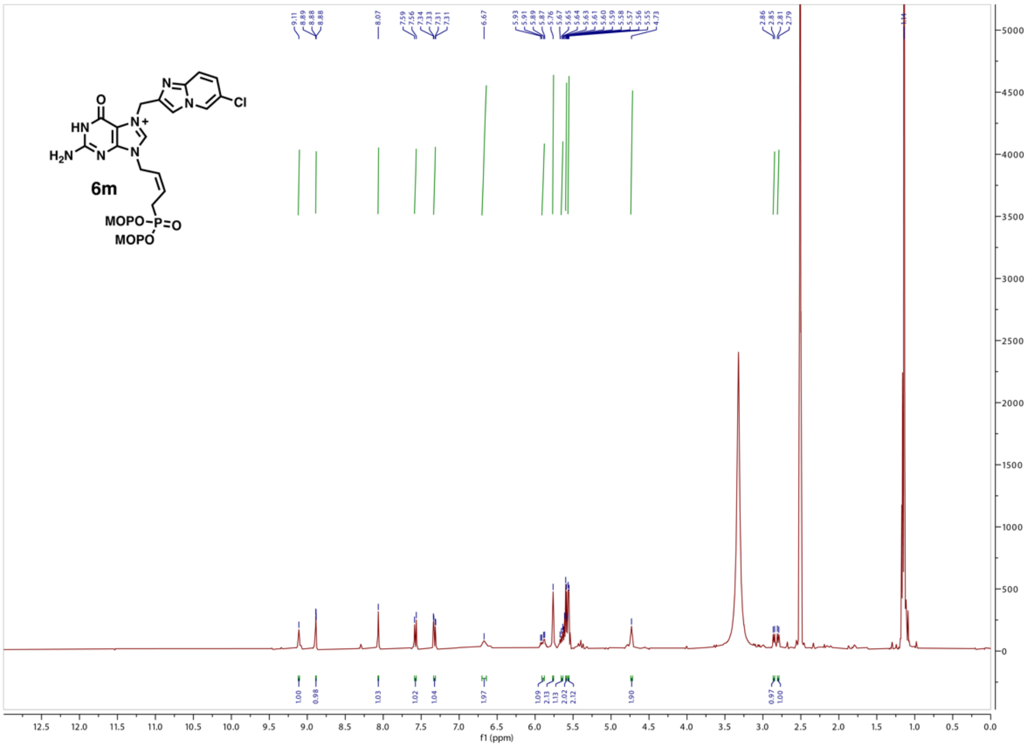

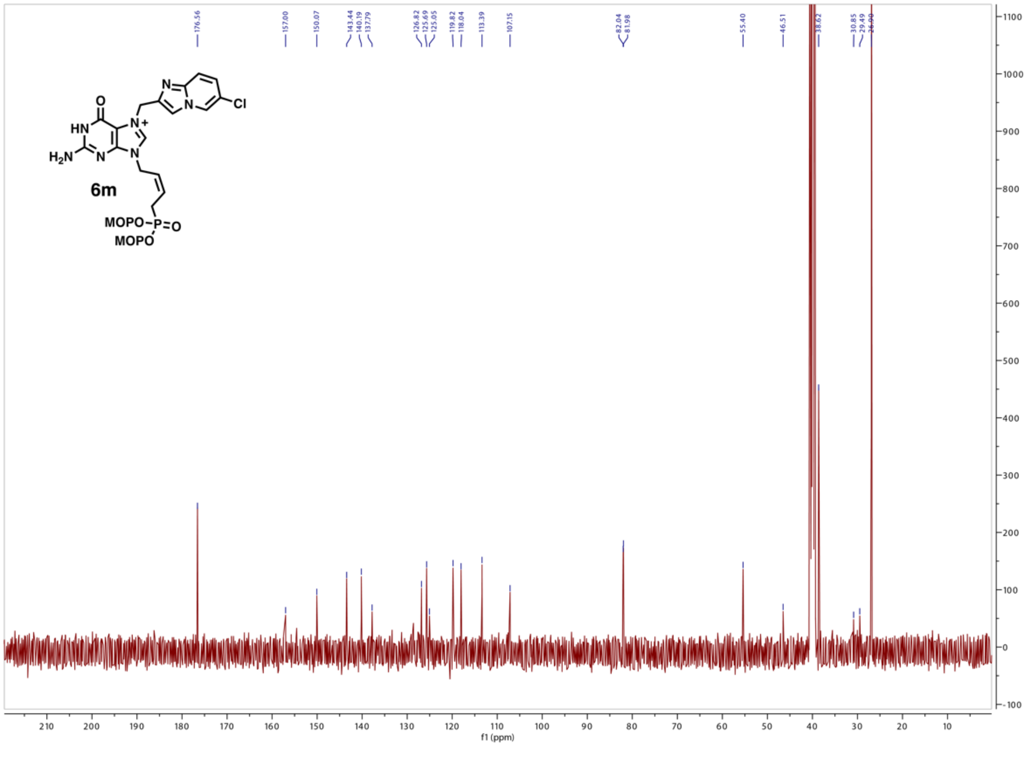

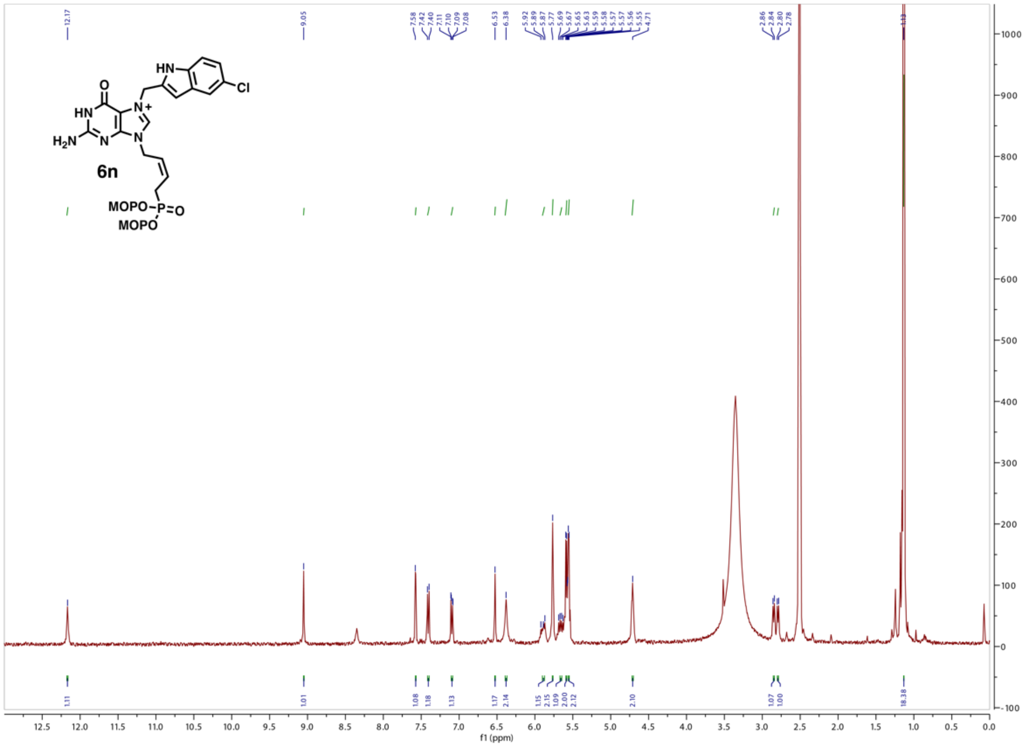
6n
MOPO
f1 (ppm)

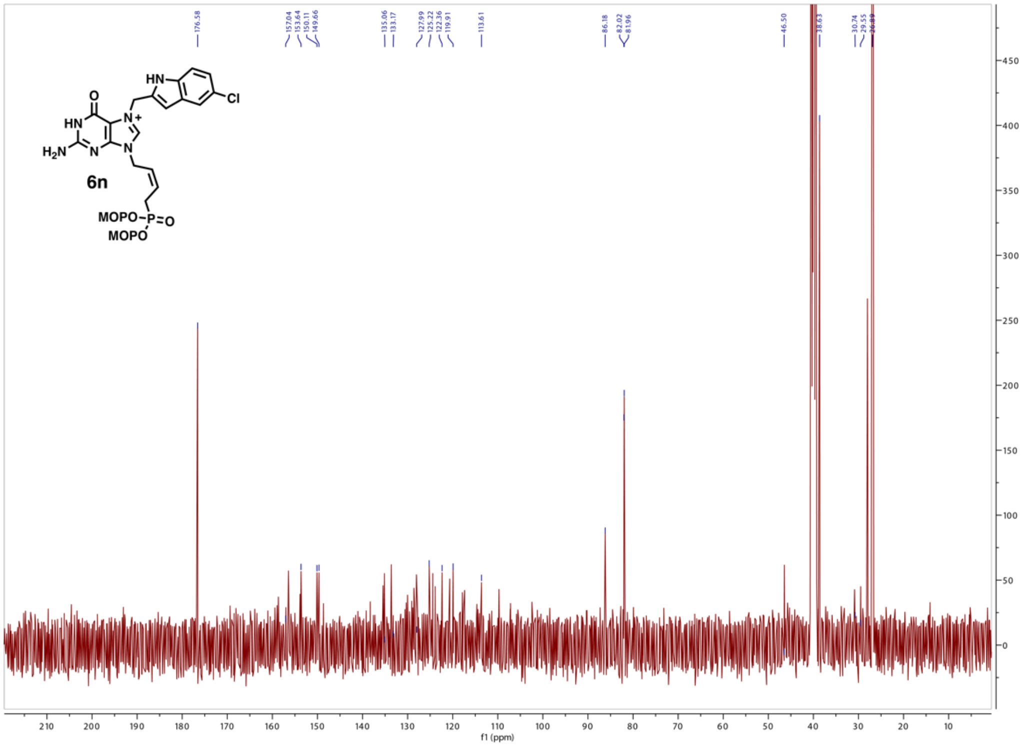
6n
f1 (ppm)

**6a:**

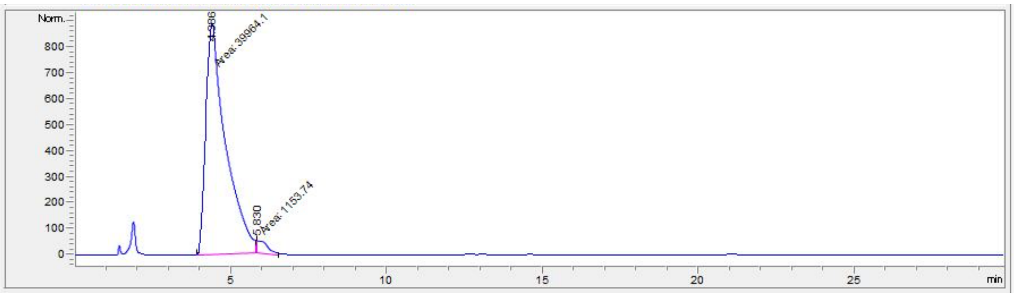


| # | Time | Area | Height | Width | Area% | Symmetry |
|---|---|---|---|---|---|---|
| 1 | 4.386 | 39964.1 | 889.6 | 0.7488 | 97.194 | 0.451 |
| 2 | 5.83 | 1153.7 | 47 | 0.4088 | 2.806 | 0 |

**6b:**

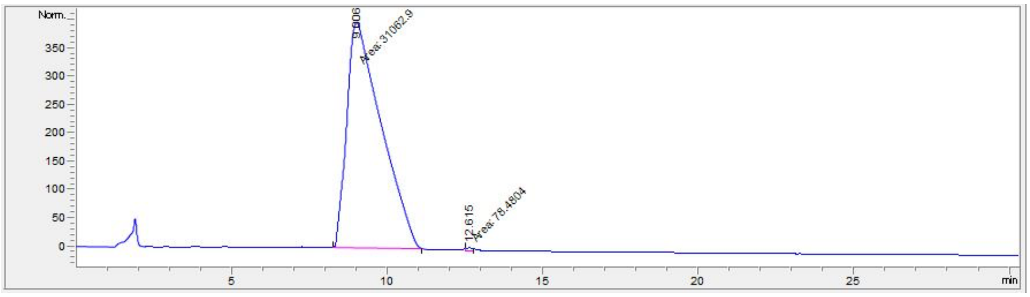


| # | Time | Area | Height | Width | Area% | Symmetry |
|---|---|---|---|---|---|---|
| 1 | 9.006 | 31062.9 | 402.4 | 1.2865 | 99.748 | 0.398 |
| 2 | 12.615 | 78.5 | 6.2 | 0.2106 | 0.252 | 0.556 |

**6c:**

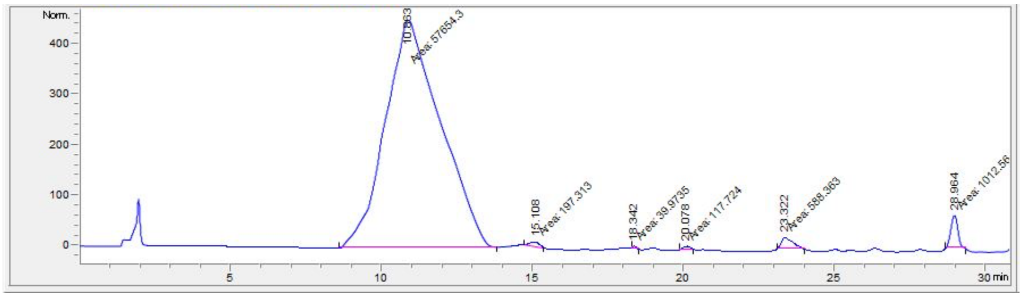


| # | Time | Area | Height | Width | Area% | Symmetry |
|---|---|---|---|---|---|---|
| 1 | 10.863 | 57654.3 | 450.9 | 2.1311 | 96.719 | 0.703 |
| 2 | 15.108 | 197.3 | 9.5 | 0.3465 | 0.331 | 1.312 |
| 3 | 18.342 | 40 | 5.4 | 0.1238 | 0.067 | 0.266 |
| 4 | 20.078 | 117.7 | 7.2 | 0.2707 | 0.197 | 1.009 |
| 5 | 23.322 | 588.4 | 21.8 | 0.449 | 0.987 | 0.343 |
| 6 | 28.964 | 1012.6 | 63.9 | 0.2639 | 1.699 | 1.077 |

**6d:**

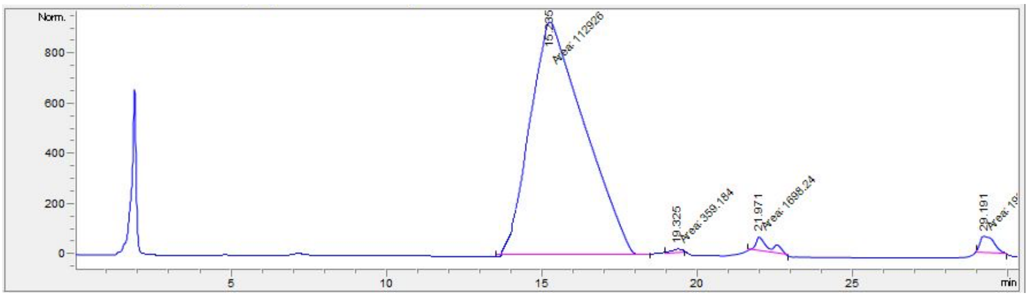

| # | Time | Area | Height | Width | Area% | Symmetry |
|---|---|---|---|---|---|---|
| 1 | 15.235 | 112925.7 | 926.9 | 2.0305 | 96.536 | 0.567 |
| 2 | 19.325 | 359.2 | 13.9 | 0.4304 | 0.307 | 1.256 |
| 3 | 21.971 | 1698.2 | 55.6 | 0.5087 | 1.452 | 0.312 |
| 4 | 29.191 | 1994.2 | 64.6 | 0.5147 | 1.705 | 0.332 |

**6e:**

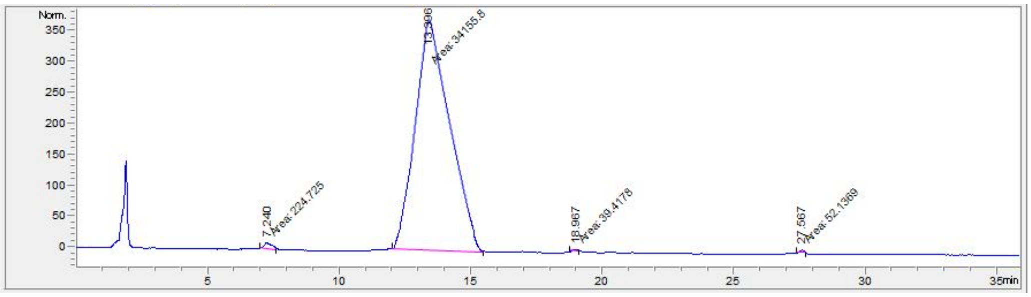

| # | Time | Area | Height | Width | Area% | Symmetry |
|---|---|---|---|---|---|---|
| 1 | 7.24 | 224.7 | 10.4 | 0.3603 | 0.652 | 0.516 |
| 2 | 13.396 | 34155.8 | 372.6 | 1.5279 | 99.083 | 0.665 |
| 3 | 18.967 | 39.4 | 3.1 | 0.2086 | 0.114 | 0.759 |
| 4 | 27.567 | 52.1 | 4.7 | 0.1852 | 0.151 | 0.758 |

**6f:**

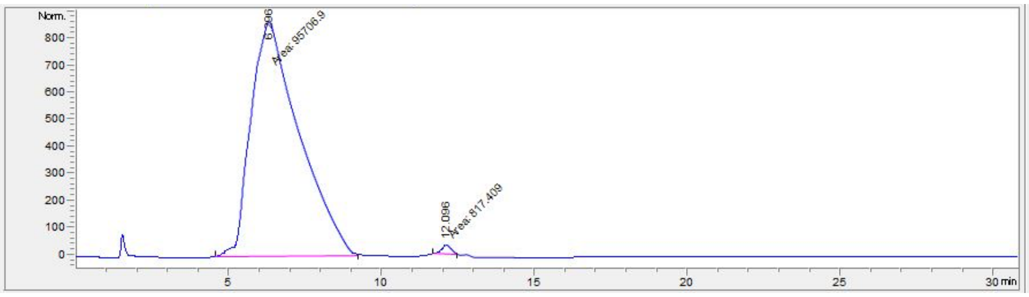

| # | Time | Area | Height | Width | Area% | Symmetry |
|---|---|---|---|---|---|---|
| 1 | 6.296 | 95706.9 | 866.4 | 1.8411 | 99.153 | 0.572 |
| 2 | 12.096 | 817.4 | 35.8 | 0.3809 | 0.847 | 0.802 |

**6g:**

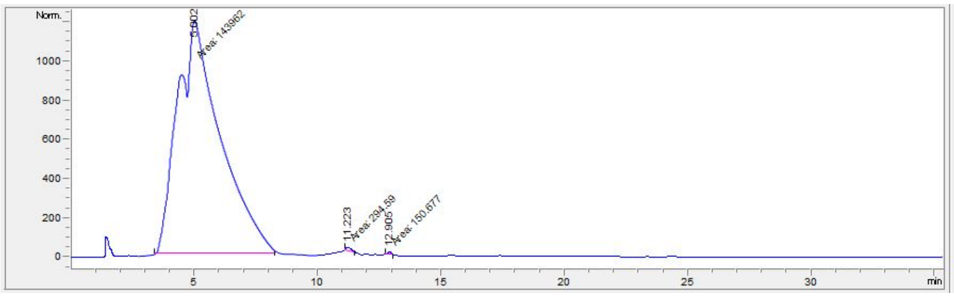


| # | Time | Area | Height | Width | Area% | Symmetry |
|---|---|---|---|---|---|---|
| 1 | 5.002 | 143961.8 | 1181.3 | 2.0311 | 99.692 | 0.646 |
| 2 | 11.223 | 294.6 | 18.2 | 0.2701 | 0.204 | 0.288 |
| 3 | 12.905 | 150.7 | 12.8 | 0.1959 | 0.104 | 1.327 |

**6h:**

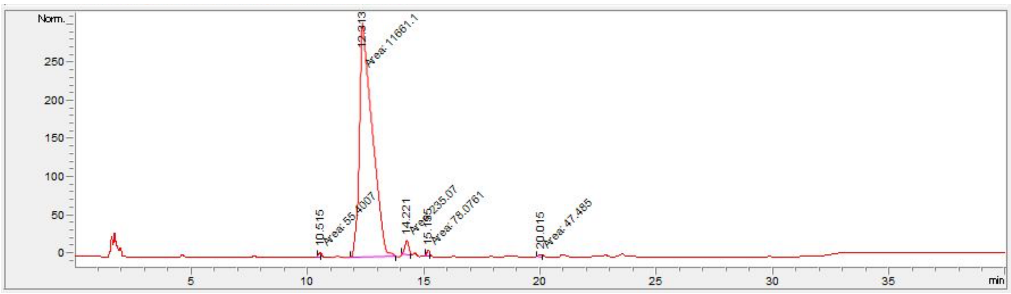


| # | Time | Area | Height | Width | Area% | Symmetry |
|---|---|---|---|---|---|---|
| 1 | 10.515 | 55.4 | 7.2 | 0.1278 | 0.459 | 0.787 |
| 2 | 12.313 | 11661.1 | 303.8 | 0.6398 | 96.555 | 0.314 |
| 3 | 14.221 | 235.1 | 19.6 | 0.2003 | 1.946 | 0.972 |
| 4 | 15.155 | 78.1 | 8 | 0.1621 | 0.646 | 0.869 |
| 5 | 20.015 | 47.5 | 3.6 | 0.2197 | 0.393 | 1.089 |

**6i:**

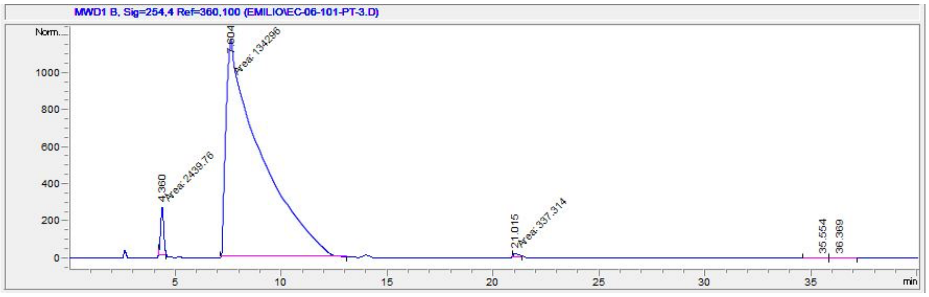


| # | Time | Area | Height | Width | Area% | Symmetry |
|---|---|---|---|---|---|---|
| 1 | 4.36 | 2439.8 | 258.7 | 0.1572 | 1.780 | 1.077 |
| 2 | 7.604 | 134296.1 | 1183.5 | 1.8912 | 97.966 | 0.163 |
| 3 | 21.015 | 337.3 | 19.8 | 0.2833 | 0.246 | 0.282 |
| 4 | 35.554 | 4.3 | 1.6E-1 | 0.3716 | 0.003 | 1.647 |
| 5 | 36.369 | 6.6 | 1.8E-1 | 0.4684 | 0.005 | 1.099 |

**6j:**

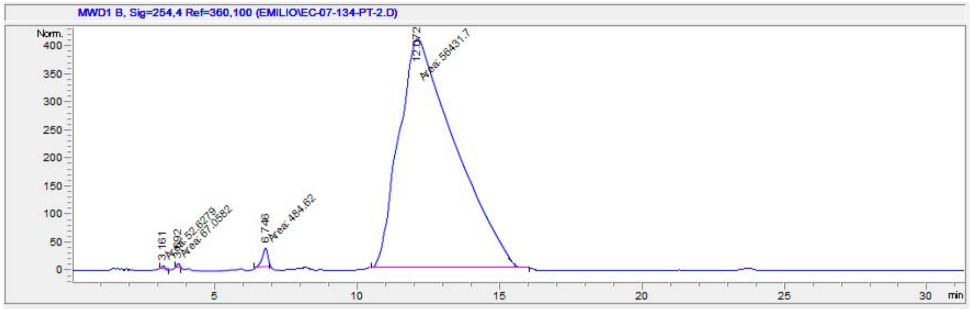


| # | Time | Area | Height | Width | Area% | Symmetry |
|---|---|---|---|---|---|---|
| 1 | 3.161 | 52.6 | 7 | 0.1256 | 0.092 | 1.312 |
| 2 | 3.692 | 67.1 | 8.7 | 0.1283 | 0.118 | 1.301 |
| 3 | 6.746 | 484.6 | 34.7 | 0.2324 | 0.850 | 1.471 |
| 4 | 12.072 | 56431.7 | 408.3 | 2.3035 | 98.940 | 0.487 |

**6k:**

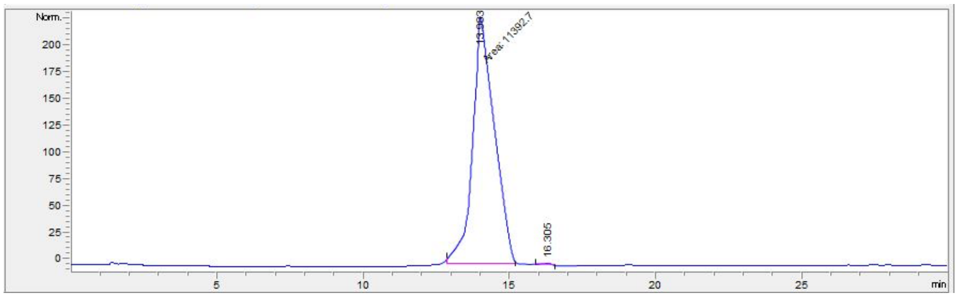


| # | Time | Area | Height | Width | Area% | Symmetry |
|---|---|---|---|---|---|---|
| 1 | 13.983 | 11392.7 | 219.8 | 0.8641 | 99.685 | 0.552 |
| 2 | 16.305 | 36 | 1.7 | 0.2861 | 0.315 | 1.726 |

**6l:**

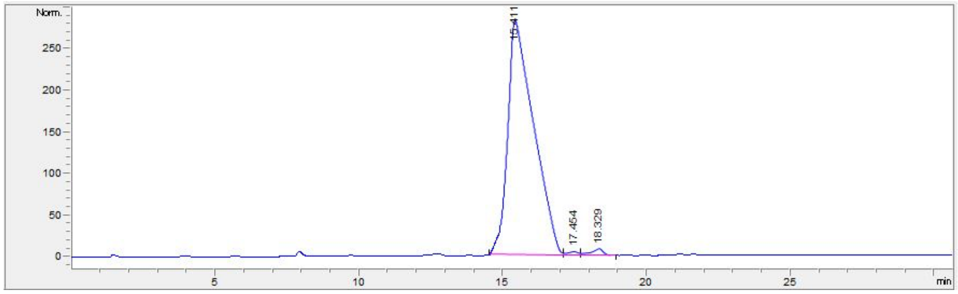


| # | Time | Area | Height | Width | Area% | Symmetry |
|---|---|---|---|---|---|---|
| 1 | 15.411 | 17839 | 283.4 | 0.8383 | 98.021 | 0.387 |
| 2 | 17.454 | 113.9 | 4.4 | 0.3658 | 0.626 | 1.335 |
| 3 | 18.329 | 246.4 | 8.2 | 0.4166 | 1.354 | 1.921 |

**6m:**

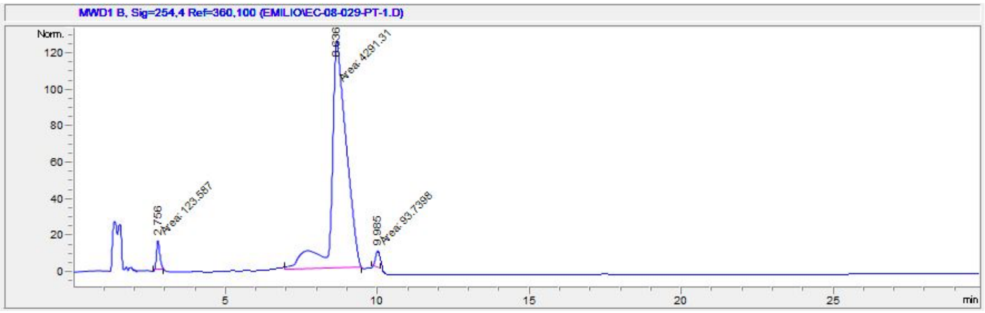


| # | Time | Area | Height | Width | Area% | Symmetry |
|---|---|---|---|---|---|---|
| 1 | 2.756 | 123.6 | 16.1 | 0.1281 | 2.741 | 0.796 |
| 2 | 8.636 | 4291.3 | 125.6 | 0.5694 | 95.180 | 0.627 |
| 3 | 9.985 | 93.7 | 9.1 | 0.1718 | 2.079 | 0.946 |

**6n:**

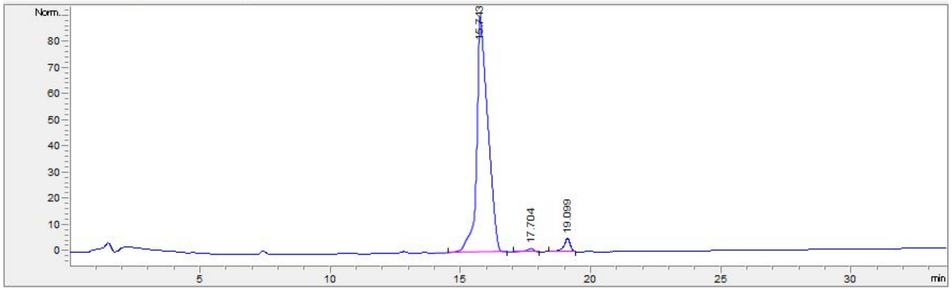


| # | Time | Area | Height | Width | Area% | Symmetry |
|---|---|---|---|---|---|---|
| 1 | 15.743 | 2667.9 | 89.3 | 0.4225 | 96.000 | 0.466 |
| 2 | 17.704 | 27.8 | 1.3 | 0.2945 | 1.001 | 2.015 |
| 3 | 19.099 | 83.4 | 5 | 0.248 | 2.999 | 1.37 |

**7a:**

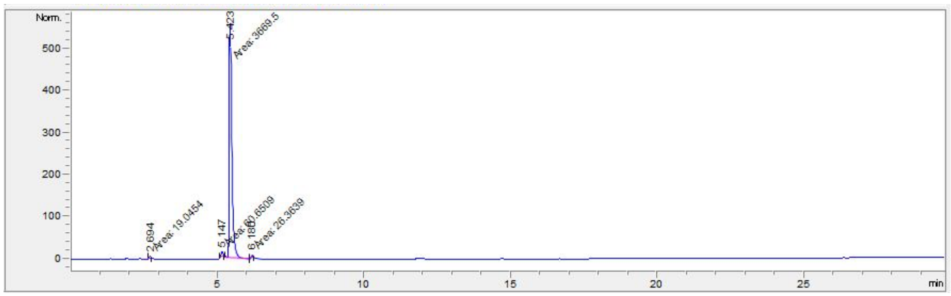


| # | Time | Area | Height | Width | Area% | Symmetry |
|---|---|---|---|---|---|---|
| 1 | 2.694 | 19 | 4.6 | 0.069 | 0.504 | 0.861 |
| 2 | 5.147 | 60.7 | 11.4 | 0.0888 | 1.606 | 1.079 |
| 3 | 5.423 | 3669.5 | 559.7 | 0.1093 | 97.191 | 0.641 |
| 4 | 6.18 | 26.4 | 6.6 | 0.0664 | 0.698 | 0.948 |

**7b:**

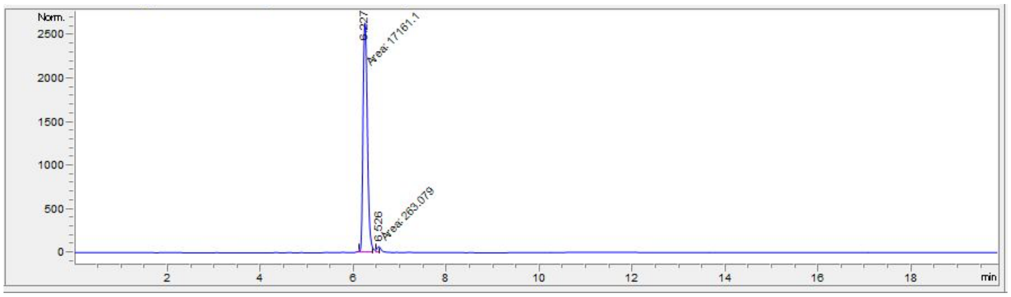


| # | Time | Area | Height | Width | Area% | Symmetry |
|---|---|---|---|---|---|---|
| 1 | 6.227 | 17161.1 | 2637.4 | 0.1084 | 98.490 | 0.733 |
| 2 | 6.526 | 263.1 | 64.1 | 0.0684 | 1.510 | 0.783 |

**7c:**

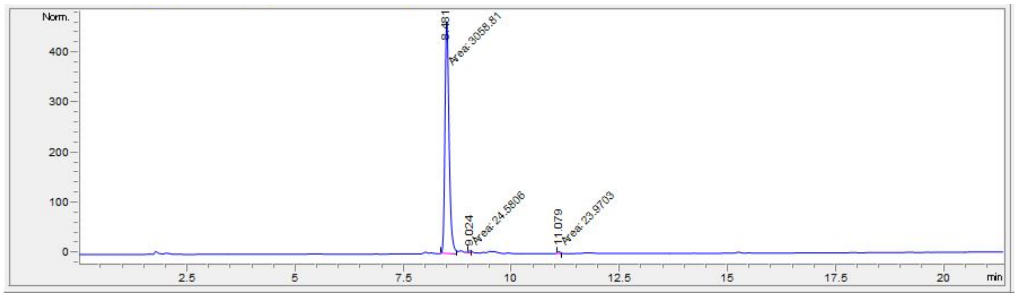


| # | Time | Area | Height | Width | Area% | Symmetry |
|---|---|---|---|---|---|---|
| 1 | 8.481 | 3058.8 | 459.5 | 0.1109 | 98.438 | 0.739 |
| 2 | 9.024 | 24.6 | 6.4 | 0.0641 | 0.791 | 0.455 |
| 3 | 11.079 | 24 | 5.1 | 0.0781 | 0.771 | 0.343 |

**7d:**

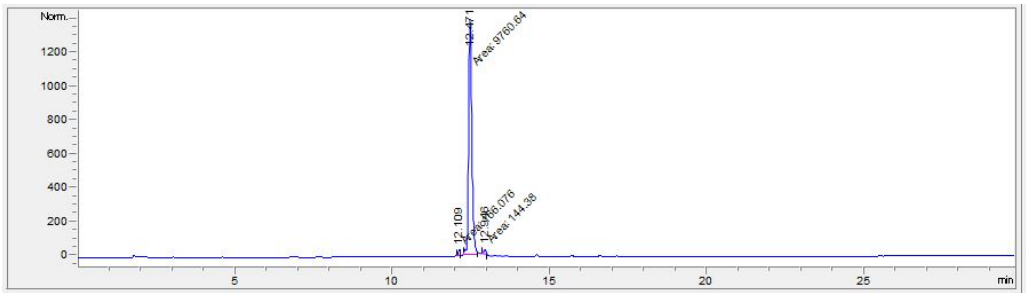


| # | Time | Area | Height | Width | Area% | Symmetry |
|---|---|---|---|---|---|---|
| 1 | 12.109 | 166.1 | 35.3 | 0.0784 | 1.649 | 0.94 |
| 2 | 12.471 | 9760.6 | 1368.1 | 0.1189 | 96.917 | 0.769 |
| 3 | 12.946 | 144.4 | 28.4 | 0.0849 | 1.434 | 0.942 |

**7e:**

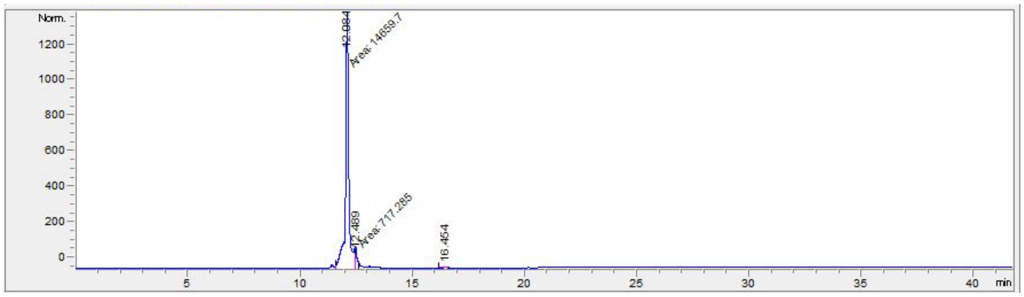


| # | Time | Area | Height | Width | Area% | Symmetry |
|---|---|---|---|---|---|---|
| 1 | 12.084 | 14659.7 | 1356.6 | 0.1801 | 95.251 | 0.99 |
| 2 | 12.489 | 717.3 | 118 | 0.1013 | 4.661 | 0 |
| 3 | 16.454 | 13.6 | 1.9 | 0.1114 | 0.088 | 0.908 |

**7f:**

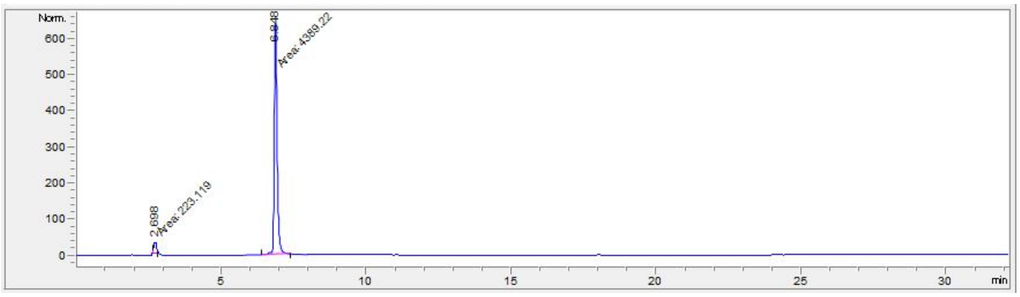


| # | Time | Area | Height | Width | Area% | Symmetry |
|---|---|---|---|---|---|---|
| 1 | 2.698 | 223.1 | 32.2 | 0.1155 | 4.837 | 0.638 |
| 2 | 6.848 | 4389.2 | 642.4 | 0.1139 | 95.163 | 0.721 |

**7g:**

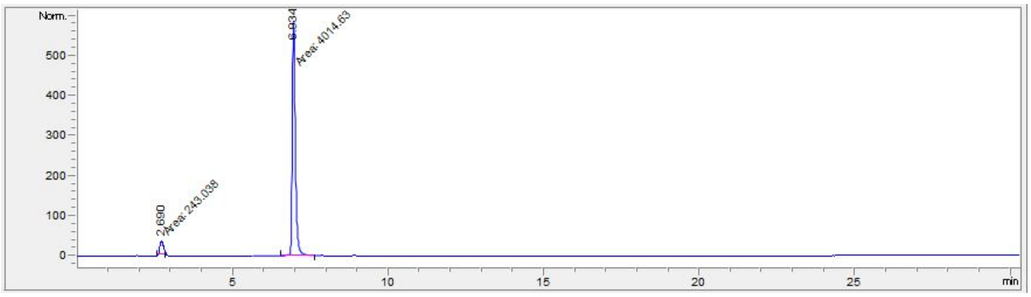


| # | Time | Area | Height | Width | Area% | Symmetry |
|---|---|---|---|---|---|---|
| 1 | 2.69 | 243 | 33.1 | 0.1225 | 5.708 | 0.715 |
| 2 | 6.934 | 4014.6 | 585.7 | 0.1142 | 94.292 | 0.688 |

**7h:**

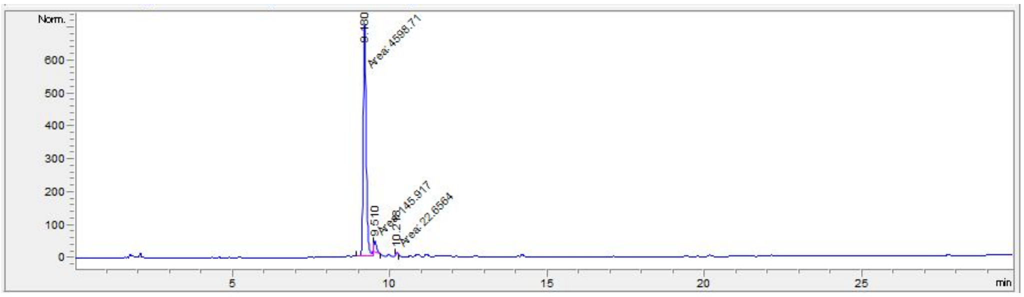


| # | Time | Area | Height | Width | Area% | Symmetry |
|---|---|---|---|---|---|---|
| 1 | 9.18 | 4598.7 | 705.3 | 0.1087 | 96.464 | 0.777 |
| 2 | 9.51 | 145.9 | 33.2 | 0.0732 | 3.061 | 0.273 |
| 3 | 10.218 | 22.7 | 6.5 | 0.0582 | 0.475 | 0.928 |

**7i:**

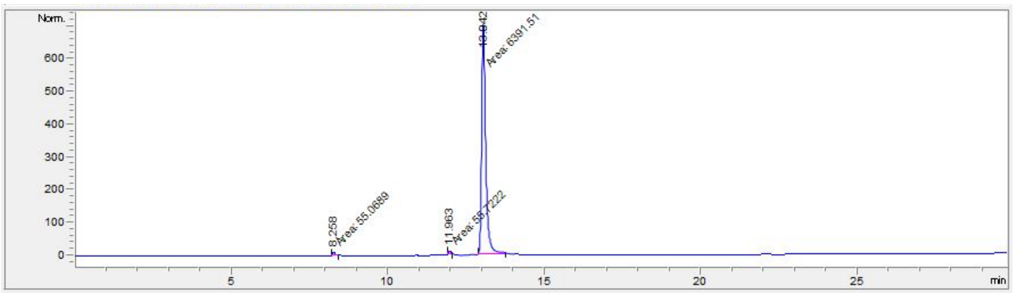


| # | Time | Area | Height | Width | Area% | Symmetry |
|---|---|---|---|---|---|---|
| 1 | 8.258 | 55.1 | 9.3 | 0.0989 | 0.847 | 0.613 |
| 2 | 11.963 | 55.7 | 12.3 | 0.0756 | 0.857 | 0.65 |
| 3 | 13.042 | 6391.5 | 702.7 | 0.1516 | 98.296 | 0.626 |

**7j:**

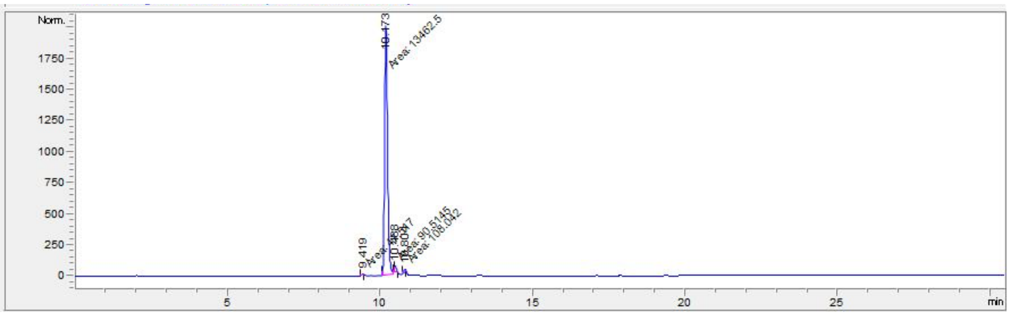


| # | Time | Area | Height | Width | Area% | Symmetry |
|---|---|---|---|---|---|---|
| 1 | 9.419 | 47.5 | 11.7 | 0.068 | 0.347 | 0.96 |
| 2 | 10.173 | 13462.5 | 2005.4 | 0.1119 | 98.205 | 0.715 |
| 3 | 10.488 | 90.5 | 44.1 | 0.0342 | 0.660 | 0 |
| 4 | 10.804 | 108 | 26.4 | 0.0681 | 0.788 | 0.715 |

**7k:**

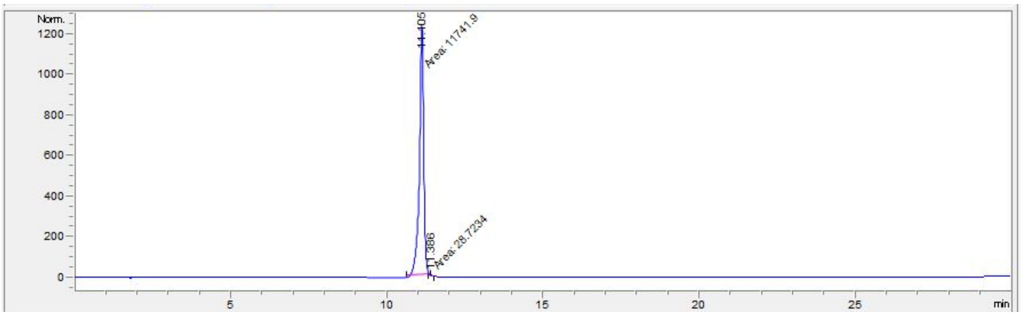


| # | Time | Area | Height | Width | Area% | Symmetry |
|---|---|---|---|---|---|---|
| 1 | 11.105 | 11741.9 | 1231.9 | 0.1589 | 99.756 | 1.35 |
| 2 | 11.386 | 28.7 | 6.8 | 0.0709 | 0.244 | 0 |

**7l:**

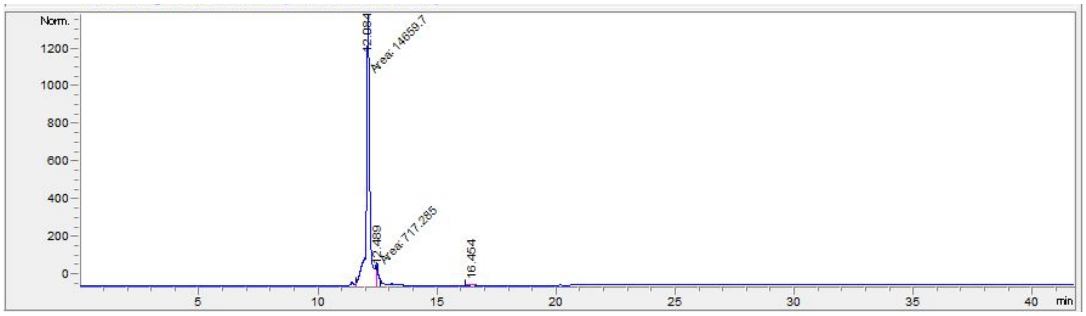


| # | Time | Area | Height | Width | Area% | Symmetry |
|---|---|---|---|---|---|---|
| 1 | 12.084 | 14659.7 | 1356.6 | 0.1801 | 95.251 | 0.99 |
| 2 | 12.489 | 717.3 | 118 | 0.1013 | 4.661 | 0 |
| 3 | 16.454 | 13.6 | 1.9 | 0.1114 | 0.088 | 0.908 |

**7m:**

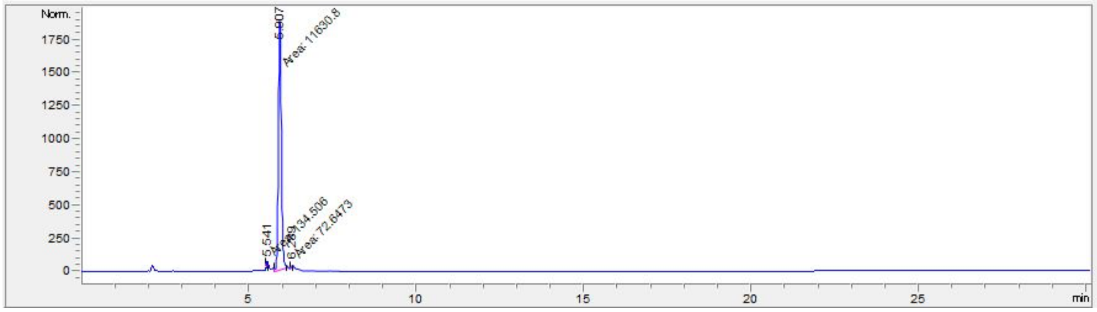


| # | Time | Area | Height | Width | Area% | Symmetry |
|---|---|---|---|---|---|---|
| 1 | 5.541 | 134.5 | 39.1 | 0.0574 | 1.136 | 0.686 |
| 2 | 5.907 | 11630.8 | 1893.1 | 0.1024 | 98.250 | 0.948 |
| 3 | 6.289 | 72.6 | 20 | 0.0605 | 0.614 | 1.005 |

**7n:**

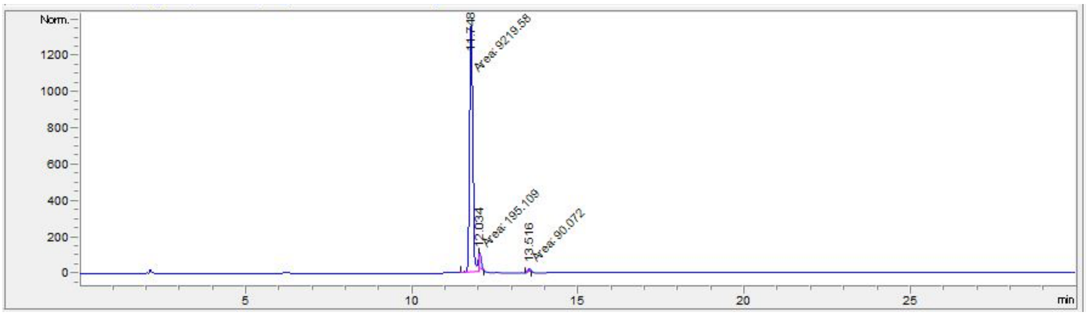


| # | Time | Area | Height | Width | Area% | Symmetry |
|---|---|---|---|---|---|---|
| 1 | 11.748 | 9219.6 | 1361.5 | 0.1129 | 97.000 | 0.735 |
| 2 | 12.034 | 195.1 | 80.3 | 0.0405 | 2.053 | 0 |
| 3 | 13.516 | 90.1 | 17.8 | 0.0844 | 0.948 | 0.932 |

### C. X-ray Crystallography Data

**Table S1: Crystallography Data Collection and Refinement Statistics**

| **Data Collection** | eIF4E-EC6012 |
|---|---|
| PDB Code | 8SX4 |
| SpaceGroup | $P2_1$ |
| Unit Cell a, b, c (Å) | 49.125, 74.328, 51.877 |
| Wavelength (Å) | 1.1271 |
| Resolution (Å)[1] | 2.0 (2.03-2.00) |
| Rsym (%)[2] | 7.4 (38.3) |
| <I/sI>[3] | 20.9 (4) |
| Completeness (%)[4] | 98.0 (99.2) |
| Redundancy | 5.2 (4.2) |
| **Refinement** | |
| Resolution (Å) | 2.0 |
| R-Factor (%)[5] | 19.7 |
| Rfree (%)[6] | 24.3 |
| Protein atoms | 2990 |
| Water Molecules | 163 |
| Unique Reflections | 24554 |
| R.m.s.d.[7] | |
| Bonds | 0.008 |
| Angles | 0.87 |
| MolProbity Score | 1.21 |
| Clash Score | 1.85 |
| Ligands | EC6012/M7GDP |
| RSCC[8] | 0.086/0.81 |
| RSR[8] | 0.15/0.17 |

[1]Statistics for highest resolution bin of reflections in parentheses.

[2]$R_{sym} = \Sigma_h \Sigma_j \mid I_{hj} - \langle I_h \rangle \mid / \Sigma_h \Sigma_j I_{hj}$, where $I_{hj}$ is the intensity of observation j of reflection h and $\langle I_h \rangle$ is the mean intensity for multiply recorded reflections.

[3]Intensity signal-to-noise ratio.

[4]Completeness of the unique diffraction data.

[5]R-factor = $\Sigma_h \mid |F_o| - |F_c| \mid / \Sigma_h |F_o|$, where $F_o$ and $F_c$ are the observed and calculated structure factor amplitudes for reflection h.

[6]$R_{free}$ is calculated against a 10% random sampling of the reflections that were removed before structure refinement.

[7]Root mean square deviation of bond lengths and bond angles.

[8] wwPDB validation service.
